# Structural basis for covalent inhibition of sulfatases by sulfamate warheads

**DOI:** 10.64898/2026.08.28.747710

**Authors:** Charles W.E. Tomlinson, Stefano Elli, Morgan Batiste-Simms, Zongjia Chen, Chris Taylor, Adam Dowle, Edwin A. Yates, Marc Nazaré, Martin A. Fascione, Lianne I. Willems, Spencer J. Williams, Conor J. Crawford, Alan Cartmell

## Abstract

The enzymatic removal of sulfate groups regulates processes ranging from steroid metabolism to carbohydrate degradation. Most sulfatases belong to the S1 family, whose members use a co-translationally installed formylglycine residue to hydrolyse sulfate esters. Arylsulfamates are potent covalent inhibitors of aryl and steroid sulfatases, including the clinical steroid sulfatase inhibitor Irosustat, yet the structure and stability of the inhibited complex remain unresolved. Arylsulfamates and carbohydrate sulfamates do not covalently inhibit many S1 carbohydrate sulfatases despite conservation of their sulfate-binding sites and formylglycine residue. Using enzyme kinetics, X-ray crystallography, molecular dynamics simulations and density functional theory calculations, we define the basis of these contrasting behaviours. High-resolution structures of the *Pseudomonas aeruginosa* arylsulfatase PaAtsA treated with two arylsulfamates reveal a long-lived tetrahedral, O-linked α-hydroxysulfamate adduct attached to formylglycine. Molecular simulations show that replacing sulfate with sulfamate disrupts the favourable Ca^2+^–oxyanion interaction and alters ligand binding geometry. The permissive hydrophobic binding site of PaAtsA accommodates this rearrangement while retaining a trajectory compatible with nucleophilic attack. By contrast, in the *Bacteroides thetaiotaomicron* carbohydrate sulfatase BT1636^3S-Gal^, sulfate-to-sulfamate substitution weakens binding and displaces the sulfamate from a reactive pose near the catalytic nucleophile due to a restrictive active site with conserved sugar binding. These findings define the structure and persistence of the arylsulfamate-derived covalent intermediate and explain why sulfamate warheads are tolerated by aryl sulfatases but not carbohydrate sulfatases.

## Introduction

Sulfation is the process by which sulfate groups are added to organic molecules. The enzymatic addition of sulfate occurs for all major classes of biological molecules including lipids^1^, steroids^2^, proteins^3^, and carbohydrates^4^ and profoundly alters their physical and biological properties. The sulfate moiety is added by sulfotransferase enzymes, which use 3′-phosphoadenosine-5′-phosphosulfate (PAPS) as the sulfate donor^5^ and is removed by enzymes called sulfatases^6^, which are essential for modifying and recycling sulfated molecules.

Sulfatase enzymes are catalogued on the SulfAtlas database and are grouped into four families, S1, S2, S3, and S4^7^. Of the >160,000 sequences catalogued on the database, ∼148,000 (∼92%) belong to the S1 family. Members of this family are distributed throughout all domains of life and catalyse the desulfation of steroids, lipids, and carbohydrates^6,8^. The other three families are exclusively limited to bacteria and have only been shown to desulfate lipid and aryl substrates^6^. The S1 family, as well as being the largest, is also unique in employing a non-genetically encoded amino acid called formylglycine (FGly) as its catalytic nucleophile. FGly is formed co-translationally by the conversion of a Cys or Ser residue, in the consensus sequence **C/S**-X-A/P-X-R, by a formylglycine generating enzyme^9^ or an anaerobic sulfatase maturation enzyme,^10^ depending on the organism and whether the environment is aerobic or anaerobic.

Two, three-step catalytic mechanisms have been proposed for S1 sulfatases: the transesterification-elimination-hydration mechanism (TEH), and the addition-hydrolysis- elimination mechanism (AHE)^8,11^. In the TEH mechanism **(Figure 1a)** the resting state of the FGly residue is the hydrated gem-diol. A conserved Asp residue, acting as a general base, activates the Oγ1 hydroxyl group of the gem-diol to attack the sulfur atom of the sulfate monoester. Concurrently, a conserved His acts as a general acid to protonate the leaving group, resulting in cleavage of the S–O bond and formation of an α-hydroxysulfate intermediate on FGly. In the second step, a second conserved His deprotonates the Oγ2 hydroxyl group of the FGly, promoting an E2 elimination of sulfate and generating the FGly aldehyde. Finally, hydration of this aldehyde regenerates the gem-diol, completing the catalytic cycle. In the AHE mechanism **(Figure 1b)**, the resting state of FGly is the aldehyde. The sulfate monoester first adds to the aldehyde carbonyl, generating a covalent α- hydroxysulfate diester adduct. In the second step, nucleophilic attack by water at sulfur cleaves the sulfate-ester bond, with the leaving group protonated by a conserved His acting as a general acid, producing a covalent α-hydroxysulfate–enzyme intermediate. Finally, this intermediate undergoes an E2 elimination, analogous to the TEH pathway, to regenerate the FGly aldehyde. Arguably, of the two mechanisms, there is more evidence in support of the TEH pathway^8,11^.

**Figure 1.**
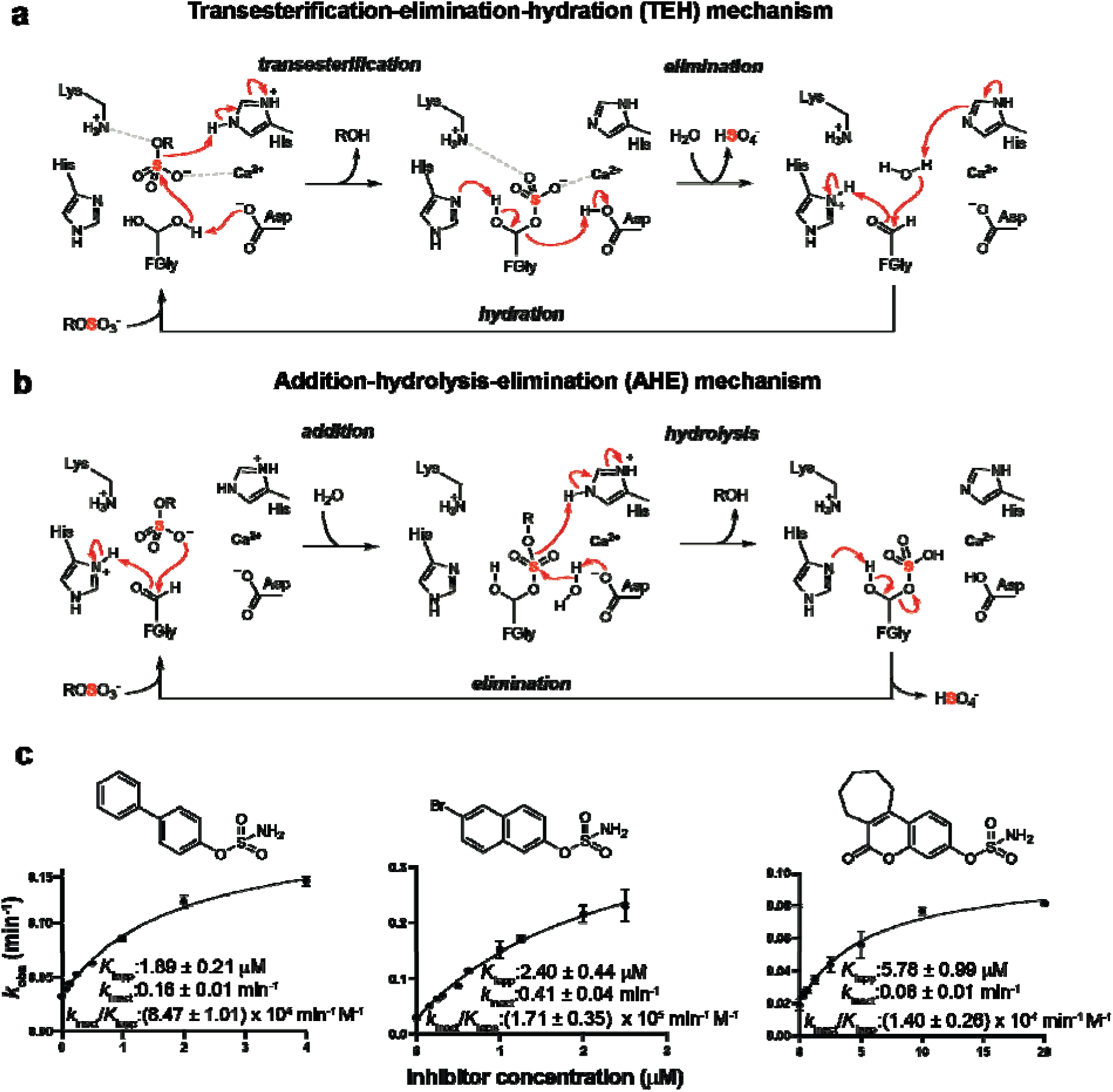
Proposed catalytic pathways for S1 sulfatases and inhibition of PaAtsA by arylsulfamates. **a.** Proposed transesterification–elimination–hydration mechanism. **b.** Proposed addition– hydrolysis–elimination mechanism. **c.** Determination of inhibition parameters for PaAtsA with 4-phenylphenyl sulfamate (PhPh-OSO_2_NH_2_), 6-bromonaphth-2-yl sulfamate, and 667 coumate (Irosustat), respectively. Enzyme assays were performed at 37 °C in 10 mM HEPES, pH 7.5, containing 150 mM NaCl and 1 mM 4-nitrophenyl sulfate. Data are from triplicate measurements and error bars represent the SD.

A well-known and clinically important member of the S1 sulfatase family in humans is steroid sulfatase (STS), which hydrolyses inactive sulfated steroid hormones to their active, unconjugated forms^12^. Sulfation of oestradiol by oestrogen sulfotransferase (SULT1E1) sequesters it into an inactive circulating pool^2^, from which it can be reactivated by STS. In the context of hormone-dependent breast cancers, inhibition of STS offers a strategy for reducing circulating oestrogen levels and thereby limiting tumour growth^13–15^. The involvement of steroid sulfation in cancer provided the impetus for the discovery of the arylsulfamates, a class of covalent sulfatase inhibitors possessing the general structure Ar– O–SO_2_NH_2_. Arylsulfamates inhibit both prokaryotic and eukaryotic S1 sulfatases^16^, including human STS^14^, and are thought to react irreversibly with the FGly nucleophile. The most advanced drug candidate is Irosustat, a first-in-class STS inhibitor that has progressed into phase II clinical trials, where it showed positive outcomes in postmenopausal women with oestrogen receptor–positive breast cancer (both early^17^ and late^18^ stages) as well as oestrogen receptor–positive endometrial cancer^19^. Despite this interest, the nature of the dead-end complex formed upon arylsulfamate inactivation of S1 sulfatases, and the mechanistic basis of its formation, is unknown.

Given the highly conserved sulfate-binding site observed across the S1 family^6,8^, sulfamate-based inhibitors might be anticipated to act as universal sulfatase inhibitors. However, subsequent work revealed that both aryl and carbohydrate sulfamates are ineffective against S1 carbohydrate sulfatases from human gut bacteria^20^. These bacterial enzymes are of biomedical interest because members of the human colonic microbiota (HCM) encode carbohydrate sulfatases that have been linked to ulcerative colitis^21–24^ and can drive colitis-like disease in susceptible mouse models^25,26^. At present, there are no selective inhibitors of HCM carbohydrate sulfatases^20^, representing an important gap in experimental capability and understanding, and constituting an unrealised opportunity for therapeutic development in inflammatory bowel disease.

Two major questions remain unanswered. First, why are arylsulfamates effective inhibitors of S1 aryl and steroid sulfatases but not of S1 carbohydrate sulfatases, despite the strong conservation of their sulfate-binding sites? Second, what is the structure of the covalent complex formed during arylsulfamate inhibition, and how stable is this adduct? To address these questions, we studied the S1_4 arylsulfatase PaAtsA, from *Pseudomonas aeruginosa* and the S1_20 carbohydrate sulfatase BT1636^3S-Gal^, from *Bacteroides thetaiotaomicron* VPI-5482 using enzyme kinetics, X-ray crystallography, molecular dynamic simulations, and density functional theory (DFT) calculations. We first examined how arylsulfates and arylsulfamates bind to and inhibit PaAtsA, then determined high-resolution structures of covalent complexes formed with two arylsulfamate inhibitors. These structures identify the inhibited state as an *O*-linked tetrahedral sulfamate adduct on FGly and, together with biochemical measurements and quantum-mechanical calculations, indicate that the adduct undergoes extremely slow breakdown. We next determined structures of BT1636^3S-Gal^ bound to sulfated glycan substrates and used molecular dynamics simulations to show that replacing sulfate with sulfamate disrupts Ca^2+^ coordination, weakens binding, and displaces the sulfur centre from the catalytic nucleophile. Finally, we examined whether increasing scaffold flexibility could overcome these geometric constraints and restore productive positioning of the sulfamate warhead within S1 sulfatase active sites.

## Results

### Binding and inhibition of PaAtsA by aryl sulfates and aryl sulfamates

The arylsulfamates 4-phenylphenyl-sulfamate (PhPh-OSO_2_NH_2_), 6-bromo-2-napthyl sulfamate, and 667-coumate sulfamate (Irosustat) exhibit irreversible inhibition with *k*_inact_/*K*_I_^app^values of (8.5 ± 1.0) × 10^4^, (1.7 ± 0.4) × 10^5^, and (1.4 ± 0.3) × 10^4^ min^-1^ M^-1^, respectively, and *K* ^app^ values in the range 2–6 μM **(Figure 1c)**. This demonstrates that a variety of aromatic structures can be accommodated in the PaAtsA binding site, consistent with previous reports^20^. Also consistent with previous observations^27^, sulfamate (NH_3_SO_3_) is not an inhibitor of PaAtsA **(Figure S1a)**, showing that the aryloxy component of arylsulfamates is required for inhibition.

To date, only a single structure has been reported of an arylsulfate with an S1 sulfatase:. the inactive C69A variant of the S1_1 glycolipid human sulfatase, ARSA, in complex with nitrocatechol sulfate (PDB: 1E2S), in which the ligand is partially disordered, limiting its utility^28^. To explore the binding of arylsulfamates and arylsulfates to an S1 arylsulfatase, we conducted molecular dynamic simulations of the inactive PaAtsA^C51S^ variant (derived from PDB 1HDH) in complex with both 4-phenylphenyl sulfate (PhPh-OSO_3_) and PhPh-OSO_2_NH_2_ **(Figure 2)**. The biphenyl group of PhPh-OSO_3_ **(Figure 2a)** is sandwiched between hydrophobic residues with Met72 and the methylene backbone of Glu74 on one side and Ile156 and Trp212 on the other. The methyl group of Thr160 sits beneath the first phenyl ring at ∼4 Å and Phe331 sits ∼3.5 Å above, and perpendicular, to the second. The sulfate group engages the Ca^2+^ ion via a head on electrostatic interaction through oxygen O1, O2 interacts with Lys113, while O3 makes no productive interaction. The sulfur atom lies within ∼3.5 Å of the Oγ of Ser51, a distance compatible with nucleophilic attack **(Figure 2a**, and **Figure S1b)**, while His211, the catalytic acid, and Lys375, coordinate the scissile sulfo-ester oxygen. This positioning of the sulfate group is in excellent agreement with the 1.3 Å PaAtsA-sulfate product complex^29^ (PDB: 1HDH), providing good support for the MD simulation **(Figure 2)**.

**Figure 2.**
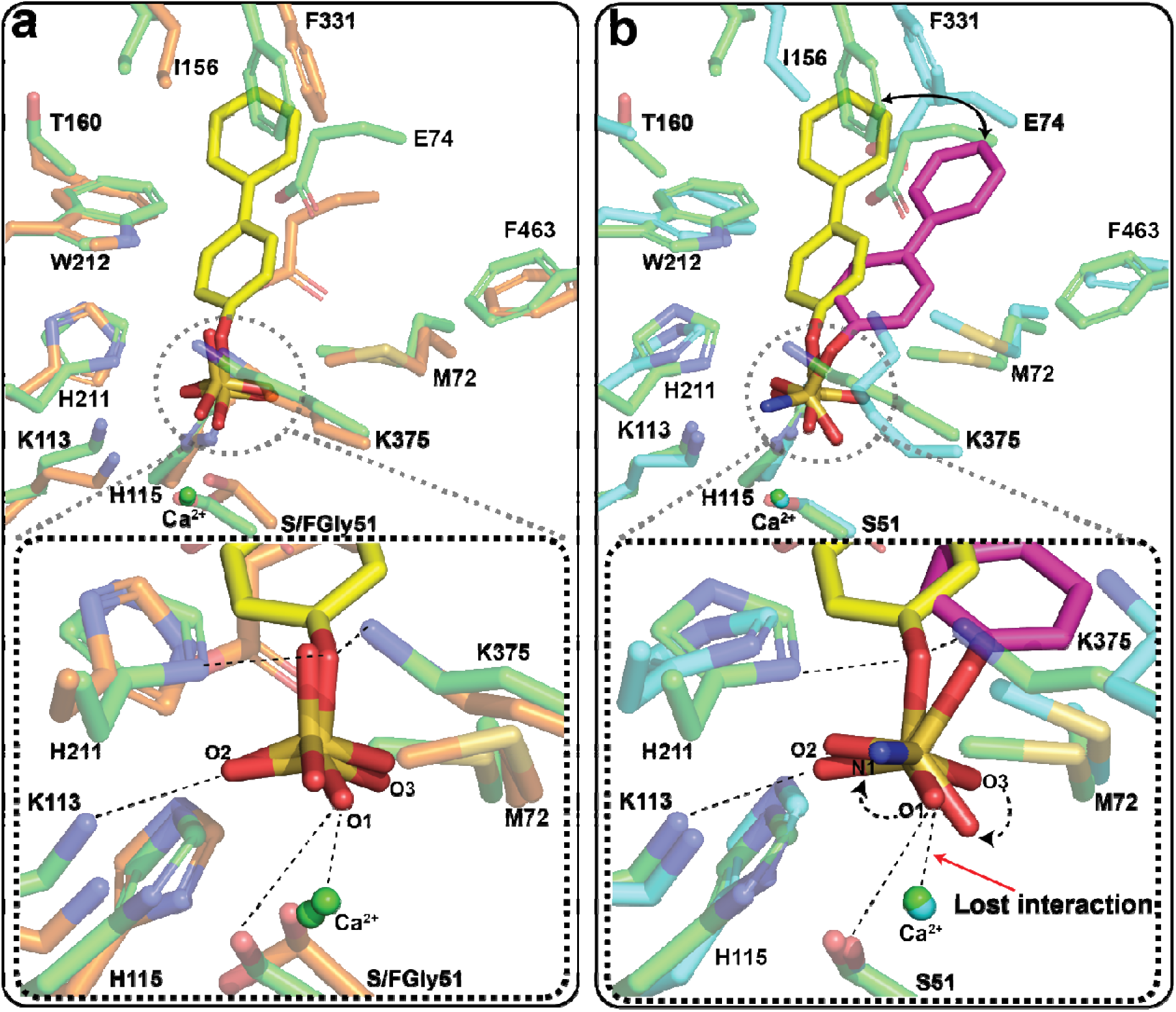
Molecular dynamics simulations of 4-phenylphenyl sulfate and 4-phenylphenyl sulfamate bound to PaAtsA. **a.** Snapshot at 120 ns of PaAtsA bound to 4-phenylphenyl sulfate. The sulfate ion from the PaAtsA-sulfate crystal structure (PDB 1HDH) is overlaid for comparison. The close agreement between the crystallographic and simulated sulfate positions supports the validity of the simulation. **b.** Snapshot at 120 ns of 4-phenylphenyl sulfamate bound to PaAtsA. The simulated 4-phenylphenyl sulfate pose (in yellow) from panel **a** is overlaid to highlight the relative changes in ligand orientation.

MD simulations with PhPh-OSO_2_NH_2_ **(Figure 2b)** predict a substantially different binding mode from that of PhPh-OSO_3_. The biphenyl group is slightly rotated, relative to that observed for PhPh-OSO_3_, although the sulfur atom occupies a similar position in both complexes, 3.5-4.0 Å from Cβ of Ser51 (**Figure S1b**). This distance is compatible with nucleophilic attack by the hydroxyl group of FGly51 in the first step of the TEH mechanism. In the sulfamate, the sulfate oxygen that coordinates Ca^2+^ is replaced by an NH_2_ group. The sulfamate group therefore rotates to avoid an unfavourable interaction between this NH_2_ group and Ca^2+^, abrogating the Ca^2+^-oxyanion interaction present in the arylsulfate complex. The hydrogen-bonding interaction between *O*2 and Lys113 is retained. The final *O*3 oxygen is also rotated, but it makes no interaction with the enzyme in either sulfate or sulfamate complex models. Most interactions involving the proximal biphenyl ring are maintained. However, the distal ring shifts away from Ile156, Thr160 and Glu74 and towards Phe463, becoming sandwiched between Phe463 and Phe330. Consistent with this altered binding mode, the S23–O22–C5–C4 dihedral angle (₂) differs markedly between the two ligands: approximately -120° for PhPh-OSO_3_ and 60° for PhPh-OSO_2_NH_2_ **(Figure S2a)**.

MD simulations indicate that the free energy of binding (ΔG^PB^), estimated using the MMPBSA approximation^30,31^, are 10.9 and 33.0 kcal mol^-1^ for PhPh-OSO_3_ and PhPh- OSO_2_NH_2_, respectively. Although both calculated values are positive, their relative magnitudes indicate weaker binding of the neutral sulfamate than of the negatively charged sulfate, highlighting the importance of electrostatic interactions. Per-residue decomposition of ΔG^PB^ provides a structural basis for this difference (**Figure S2b**). Although most interactions are retained in the altered arylsulfamate binding mode, interactions with Lys113 and Lys375 are weakened. In particular, the distances to Lys375 increase by 2-4 Å **(Figure S1b)**, and this residue no longer makes a significant favourable contribution to the calculated binding free energy **(Figure S2b)**. These changes are also associated with modest differences in the hydrogen bonding networks formed by PhPh-OSO_3_ and PhPh-OSO_2_NH_2_ within PaAtsA **(Table 1)**. Overall, the simulations suggest that the active site can accommodate either a sulfate or sulfamate group when presented on an aryl scaffold, although the difference in charge and structure of the sulfamate group leads the arylsulfamate to bind less favourably and in a different pose.

**Table 1.** Hydrogen bond occupancies between PhPh-O-SO_3_ or PhPh-O-SO_2_-NH_2_ and arylsulfatase PaAtsA (PDB 1HDH), predicted by MD simulation.

| PhPh-O-SO<sub>3</sub> |  | PhPh-O-SO<sub>2</sub>-NH<sub>2</sub> |  |
| --- | --- | --- | --- |
| H-bond | PoP % | H-bond | PoP % |
|  |  | H211-Nδ1---H-N-S-4PhPh | 38 |
| K113-Nζ-H---O-S-4PhPh | 74 | K113-Nζ-H---O-S-4PhPh | 23 |
| S51-OγH---O-S-4PhPh | 5.7 | S51-OγH---O-S-4PhPh | 13 |
| K375-Nζ-H---O-S-4PhPh | 12 | K375-Nζ-H---O-S-4PhPh | 11 |
|  |  | W212-Nε1-H---O-S-4PhPh | 3 |

### The nature of the sulfamate dead-end complex

Arylsulfamates are proposed to inhibit arylsulfatases irreversibly by forming a covalent adduct to the nucleophilic FGly residue, although direct evidence for this mechanism is lacking. To investigate the nature of the dead-end complex, we first used mass spectrometry. Wildtype PaAtsA (20 μM) was incubated overnight with or without Irosustat (1 mM). Activity assays revealed that approximately 90% of the PaAtsA was inhibited upon incubation with Irosustat **(Figure S3a)**. The samples were then analysed by peptide-level and intact-protein mass spectrometry. Peptide analysis identified both the formyl and hydrated forms of the catalytic FGly residue (representing the aldehyde and gem-diol forms, respectively) in roughly equal proportions **(Figure S3b)** but no mass shift consistent with a sulfamate-derived adduct was detected in the Irosustat-treated sample. Intact-protein mass spectrometry also provided no evidence of covalent modification; however, in this case only the hydrated gem-diol form of FGly was detected in both samples suggesting that this is the predominant resting state for the enzyme **(Figure S3c)**.

We therefore turned to X-ray crystallography to visualize the dead-end complex. Wildtype PaAtsA was crystallised with arylsulfamate inhibitors using two approaches: (i) co-crystallisation with Irosustat, and (ii) incubation with 6-bromo-2-napthyl sulfamate for approximately 3 days, followed by size exclusion chromatography to remove unbound species and crystallisation of the purified protein. Both approaches yielded high-resolution electron density maps with unambiguous density for a tetrahedral adduct attached to FGly.

The Irosustat complex, solved at 1.6 Å, contained a covalent adduct at full occupancy, with the coumarin group also present, but in a reversed orientation in the active site **(Figure 3ai)**. By contrast, the complex obtained after preincubation with 6-bromo-2- naphthyl sulfamate, solved at 1.5 Å, contained an FGly adduct modelled at 85% occupancy **(Figure 3ai)**. The retained aromatic coumarin derived from Irosustat interacts with a constellation of hydrophobic residues, including Met72, Ile156, Trp212, Phe331, Phe328, Phe463, and Phe533 **(Figure S4a)**, similar to that described above for PhPh-OSO_3_. The lactone group of the coumarin interacts with both the adduct and the putative catalytic acid His211. In the complex derived from 6-bromo-2-napthyl sulfamate, a water molecule occupies the corresponding position **(Figure 3ai)**.

To establish whether the adduct was linked to FGly through nitrogen or oxygen, we modelled both possibilities and compared their fit to the electron-density maps and the resulting atomic B-factors. At this resolution, incorrect assignment of N and O can be detected because atoms within the same covalent group should refine with comparable B-factors; a pronounced mismatch therefore supports the alternative atom assignment. For both complexes, the *O*-linked model provided the better fit. The N and O atoms of the *O*- linked adduct refined to similar B-factors, whereas exchanging their assignments to produce an *N*-linked model resulted in a higher B-factor for the atom modelled as oxygen, indicating that nitrogen occupies this position **(Figure 3aii,bii)**. The *N*-linked models also produced weak negative features in the mFobs−DFc difference maps, which were absent from the corresponding *O*-linked models. Together, these analyses support assignment of an *O*- linked sulfamate FGly adduct.

To further investigate the structural assignment, we analysed the two adduct alternatives by density functional theory (DFT) using a reduced model of the PaAtsA active site (see Materials and Methods). The calculated internal energy content of the *O*-linked adduct was 305.6 kcal mol^-1^ lower than that of the *N*-linked isomer **(Table 2)**. Moreover, the geometry predicted for the tetrahedral FGly–OSO_2_NH_2_ unit agreed more closely with the crystallographic model than did that predicted for the alternative *N*-linked FGly–NH-SO_3_^−^ structure. For the *O*-linked adduct, DFT gave an FGly O–S bond distance of 1.66 Å, S=O bond distances of 1.46 and 1.47 Å, and an S–N bond distance of 1.55 Å. These values agree reasonably well with the corresponding distances of 1.58, 1.44, 1.44 and 1.64 Å, respectively, in the crystallographic model derived from the 6-bromo-2-naphthyl sulfamate complex **(Figure 3c)**. By contrast, the DFT-optimised *N*-linked structure had an N–S bond distance of 1.73 Å and three similar S–O bond distances of 1.46, 1.47 and 1.48 Å, which were in poor agreement with the corresponding distances in the *N*-linked crystallographic model: 1.59 Å for N–S and 1.50, 1.44 and 1.43 Å for the three S–O bonds **(Figure 3d)**. Collectively, the crystallographic and computational results identify the covalent complex as a tetrahedral sulfamate adduct linked through oxygen to the hydrated form of FGly.

**Figure 3.**
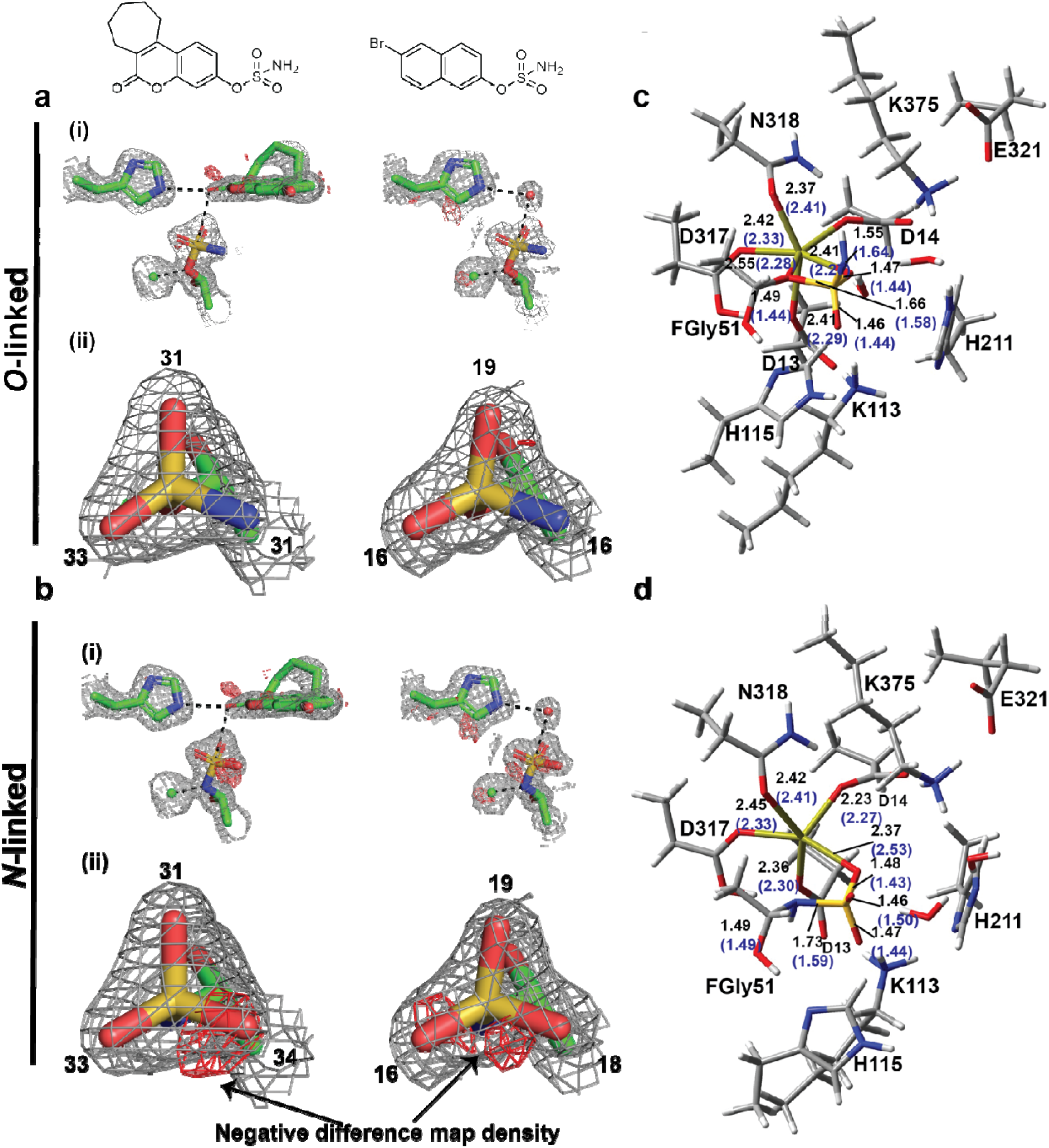
Structural identification of the O-linked covalent adduct formed by arylsulfamates on PaAtsA. Stick representations of the covalent adducts on PaAtsA with the corresponding 2mFobs– DFc and mFobs–DFc maps contoured at 1.5 σ and 3.0 σ, respectively. **a.** Models of the *O*-linked adducts. **b.** Corresponding models of the alternative N-linked adducts. For each complex, **(i)** shows the covalent adduct and its interactions with the catalytic acid His211, Ca^2+^, water molecules and the retained leaving group, whereas **(ii)** shows a view directly from directly above the covalent adduct with atomic B-factors indicated for the relevant atoms. **c,d** Density functional theory models for the O-linked and N-liked adducts, respectively. Bond lengths shown in black are derived from the quantum-mechanical calculations; blue values shown in parentheses are from the corresponding *O-*linked or *N-* linked crystallographic models.

**Table 2.** DFT-calculated energies of the O-linked and N-linked adducts. The electronic energy (E^B3LYP^), zero point energy (E^ZP^) and their sum (E^B3LYP^ + E^ZP^) are reported in hartree (see Materials and Methods).

| Energy (hartree) | FGly-O-SO<sub>2</sub>-NH<sub>2</sub> | FGly-NH-SO<sub>3</sub> |
| --- | --- | --- |
| $E^{B3LYP}$ | -4141.2810 | -4140.7828 |
| $E^{ZP}$ | 1.2311 | 1.2199 |
| $E^{B3LYP} + E^{ZP}$ | -4140.0499 | -4139.5629 |

**Table 3.** Hydrogen bond occupancies between 3SLacNAc and 3SNH_2_ LacNAc and the carbohydrate sulfatase BT1636^3S-Gal^, that were predicted by MD simulation.

| 3SLacNAc BT1636<sup>3S-Gal</sup> |  | 3SNH<sub>2</sub>LacNAc BT1636<sup>3S-Gal</sup> |  |
| --- | --- | --- | --- |
| H-bond | PoP % | H-bond | PoP % |
| H177-Nε2-H---O4-Gal | 63 | E100-COO---HO6-Gal | 95 |
| E334-COO---HO2-Gal | 58 | Q173-Oε1---HN-SO2-Gal | 66 |
| E100-COO---HO6-Gal | 58 | Y284-OH---O2-Gal | 50 |
| R353-NH<sup>+</sup>---O2-Gal | 54 | E334-COO---HO6-GlcNAc | 41 |
| Q173-Nε2H---O6-Gal | 50 | G281-O---HN-GlcNAc | 38 |
| E100-COO---HO3-GlcNAc | 42 | Q173-O---HO4-Gal | 18 |
| N98-NδH---O-S-O-Gal | 33 |  |  |
| K147-NH---O-S-O-Gal | 31 |  |  |

### The FGly–OSO_2_NH_2_ covalent adduct undergoes slow turnover

Three observations suggested that the covalent adduct may not be a dead-end product. First, overnight incubation of PaAtsA with Irosustat resulted in approximately 90% inhibition rather than complete loss of activity **(Figure S3a)**. Second, in the structure obtained with 6- bromo-2-naphthyl sulfamate, the *O*-linked adduct was best modelled at 85% occupancy, with a water molecule positioned above the sulfur centre **(Figure 3a(i))**. By comparison, the Irosustat-derived adduct was best modelled at full occupancy, with its retained aryloxy group capping the adduct and the lactone positioned above the sulfur, potentially shielding it from bulk solvent **(Figure 3a(i)).** Finally, the adduct could not be detected by MS analysis of the protein, either intact or after proteolysis. These observations suggest that the *O*-linked adduct may undergo hydrolysis if accessible to water. Interestingly, the C51S variant, PaAtsA^C51S^, which cannot undergo the proposed elimination step in the TEH and AHE mechanisms, cleaves 4NP-SO_3_ at a low rate **(Figure S4b)**. A structure of PaAtsA^C51S^ with bound sulfate, at 50% occupancy (derived from reaction with 4NP-SO_3_), revealed a water molecule coordinated by the catalytic acid and positioned above the sulfate, appropriately orientated for nucleophilic attack suggesting hydrolysis as the catalytic mechanism **(Figure S4c)**. A similarly positioned water molecule is observed in the 6-bromo-2-naphthyl sulfamate-derived complex **(Figure 3a(i))**.

Together, these observations suggest that the *O*-linked adduct is not a dead-end complex but is slowly turned over to regenerate active enzyme. To test this hypothesis, frozen PaAtsA used to obtain the 6-bromo-2-naphthyl sulfamate-derived crystal structure was thawed and assayed periodically over 4 weeks. The inhibited enzyme progressively regained activity, with the largest increase occurring during the first week **(Figure 4a)**. Recovery was evident both as an absolute increase in activity and relative to an uninhibited control, which gradually lost activity over the same period. These results demonstrate slow breakdown of the covalent adduct with recovery of enzymatic activity.

**Figure 4.**
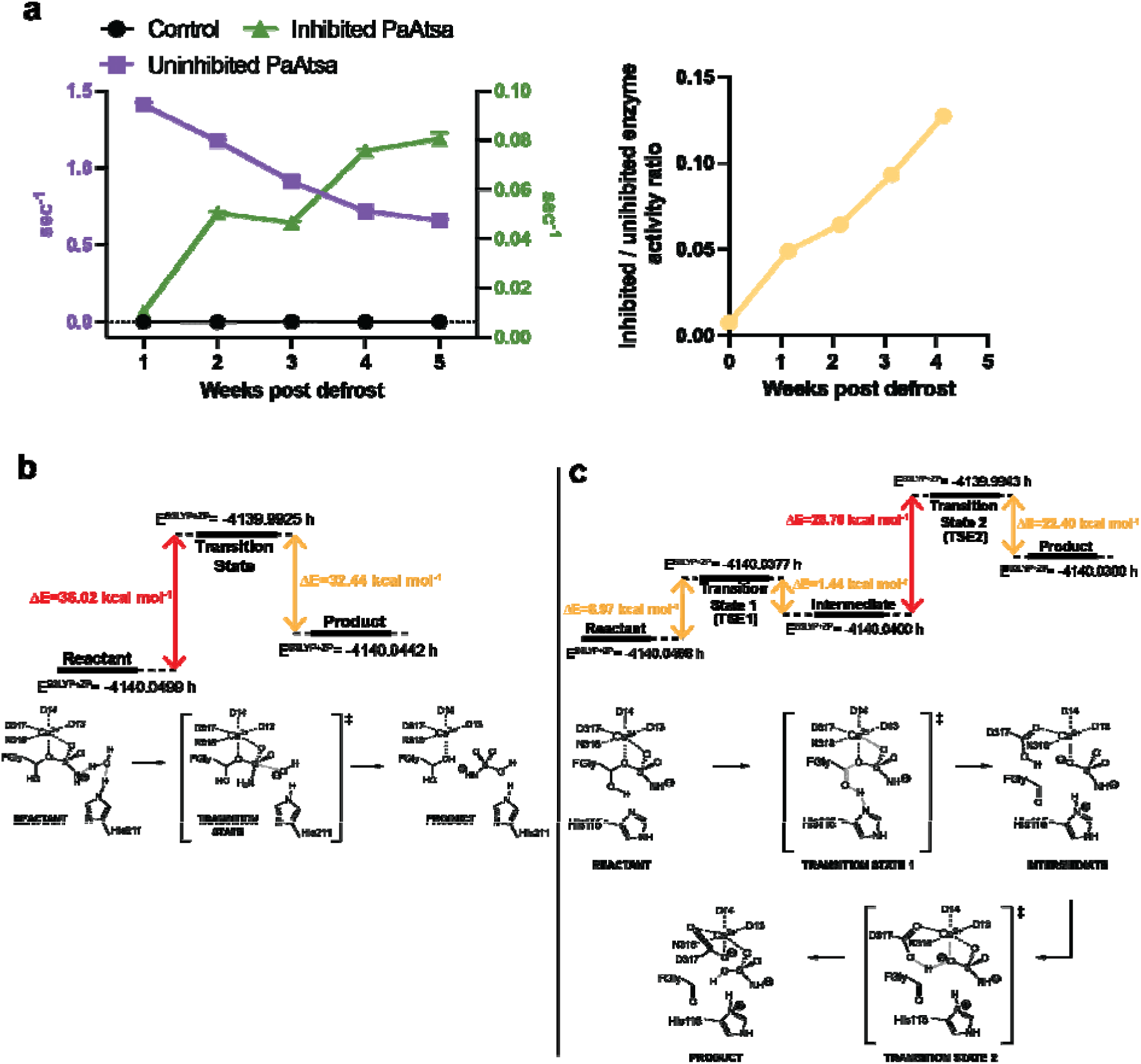
Time-dependent breakdown of the covalent adduct through two potential mechanisms. **a.** Activity of uninhibited PaAtsA and PaAtsA containing the α-hydroxysulfamate adduct over four weeks. Activity was measured using 1 mM 4-nitrophenyl sulfate in 10 mM HEPES, pH 7.5, containing 150 mM NaCl. **b.** Proposed substitution hydrolysis pathway for breakdown of the O-linked PaAtsA intermediate. Top: DFT-derived internal energy profile showing the reactant, transition state (TS) and product. Internal energies (E^B3LYP+ZP^) are reported in hartree, and energy differences between states are reported in kcal mol^-1^; the estimated activation energy is underlined in red. Bottom: reaction scheme for the substitution hydrolysis pathway, with the reactant, transition state and product underlined. **c.** Proposed two-step elimination pathway for breakdown of the O-linked PaAtsA intermediate. Top: DFT-derived internal energy profile showing the reactant, transition states (TSE1 and TSE2), the intermediate and product. Internal energies and energy differences between states are reported as described above. The estimated activation energy is in red. Bottom: reaction scheme for the two-step elimination pathway, with the reactant, transition states, intermediate and product indicated.

We used DFT calculations with a truncated active-site model to compare the energy profiles for substitution and elimination hydrolysis pathways of the FGly-OSO_2_NH_2_ adduct (**Figure 4b,c)**. In the case of hydrolysis by substitution, the reaction was predicted to proceed through a single-step nucleophilic substitution at sulfur by water, with an activation energy of 36.0 kcal mol^-1^ (**Figure 4b)**. Alternatively, elimination of the adduct was predicted to occur in two steps, through transition states TSE1 and TSE2. The higher barrier, associated with TSE2, was 28.7 kcal mol^-1^ (**Figure 4c).** In the first step of the elimination pathway, C_β1_ of FGly changes from *sp*^3^ to *sp*^2^ hybridization to form the aldehyde, generating an intermediate in which a ^−^OSO_2_NH^−^ dianion remains coordinated to Ca^2+^ (Ca^2+^–O distance, 2.33 Å). In the second step, proton transfer from Asp317 to the Ca^2+^ coordinated oxygen results in formation of the product. The resulting HO-SO_2_-NH^−^ monoanion occupies the Ca^2+^ coordination site distal to FGly, leaving the proximal site vacant.

The calculated geometries of the reactant, transition states, intermediate and products for both pathways are shown in **Figure S5(i)-(iv) and Figure S6(i)-(v)**, while their energies and fundamental vibrational frequencies are reported in **Tables S1** and **S2**. The calculations predict high barriers for both pathways consistent with the slow breakdown of the adduct, but with elimination favoured.

### Crystal structures of the S1_20 enzyme BT1636^3S-Gal^ bound with substrate and product

Our previous work showed that the sulfamate glycomimetics Gal *O*2- and *O*3-sulfamate do not bind to or inhibit BT1636^3S-Gal^ ^20^. To investigate why, we determined high-resolution crystal structures of an inactive Ser77 variant bound to the substrates 3S-LacNAc, 3S- Lewis^x^, and 3S-Lewis^a^, and the product LacNAc **(Figure 5a)**. All four ligands adopted the same mode of Gal binding in the 0 subsite, and sulfate recognition was conserved across the complexes with the three sulfated substrates. The sulfate moiety of the substrates sits in the S site and coordinates Ca^2+^, completing its octahedral coordination geometry. It also interacts with Ser77, the precursor of the catalytic FGly residue, and with Lys147, Lys353, and His252, with His252 proposed to act as general acid/base catalyst. The axial C4 hydroxyl of Gal, which distinguishes Gal from its C4 epimer Glc, is recognised by His177, and the C2 hydroxyl interacts with Glu334 and Arg353 **(Figure 5a)**. The *O*6 of Gal forms hydrogen bonds with Glu100 and Gln173. These features are broadly consistent with those reported for a lower-resolution structure^32^.

**Figure 5.**
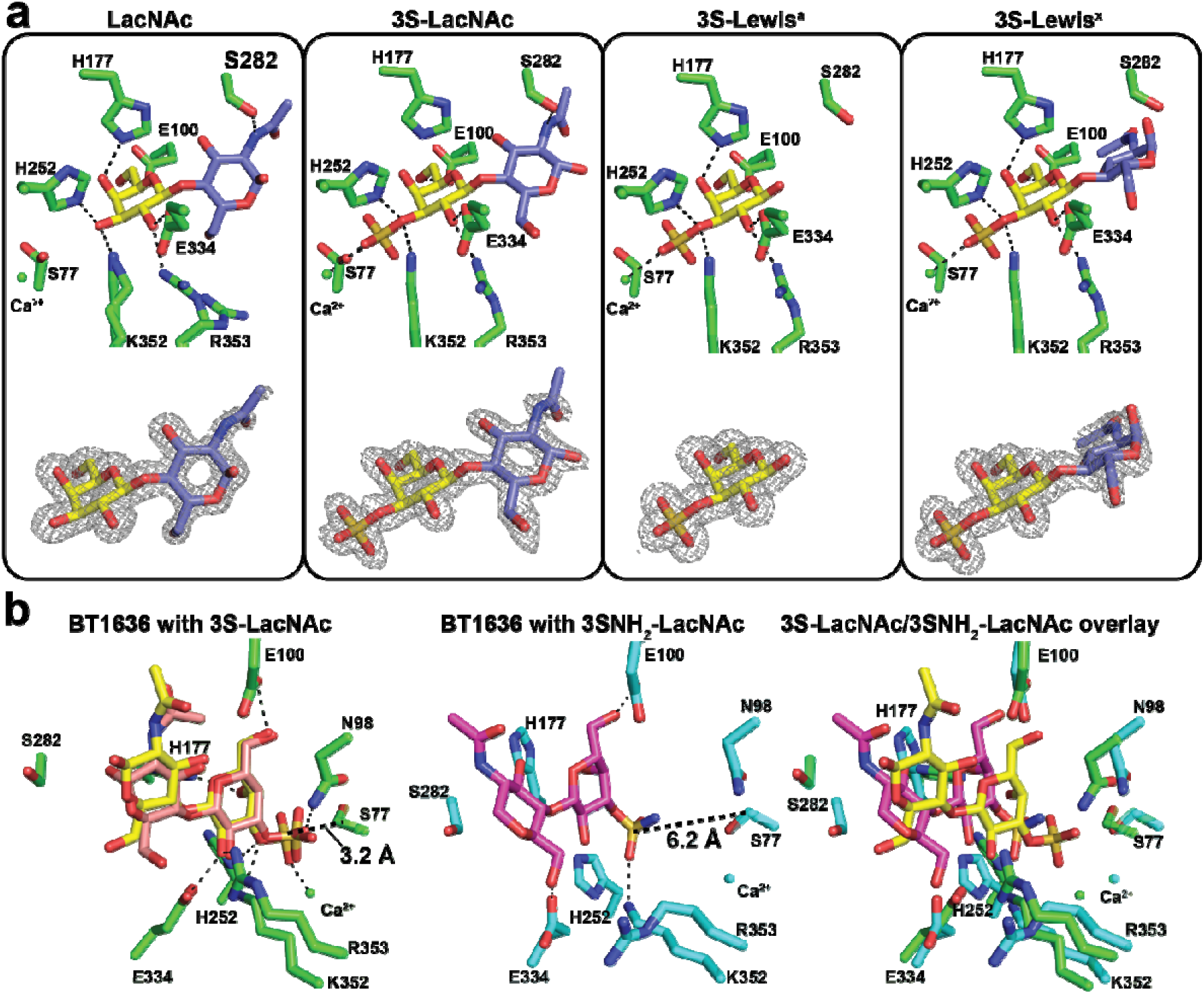
Interrogating the binding of sulfated substrates and a sulfamate substrate mimic to BT1636^3S-Gal^ using with high resolution X-ray crystallography and molecular dynamics. **a.** Stick representations of BT1636^3S-Gal^ bound to LacNAc, 3S-LacNAc, 3S-Lewis^a^, and 3S-Lewis^x^, determined at resolutions of 1.4, 1.45, 1.41, and 1.42 Å, respectively. The corresponding 2mFobs–DFc electron density maps are shown below each structure and contoured at 1 σ. **b.** Left, snapshot at 83 ns of BT1636^3S-Gal^ bound to 3S-LacNAc. The simulated ligand closely matches the superimposed 3S-LacNAc conformation from the 3S-Lewis^X^ crystal structure. Middle: snapshot at 110 ns BT1636^3S-Gal^ bound to O3-sulfamated LacNAc (3S-NH_2_-LacNAc). Right: overlay of the simulated 3S-LacNAc and 3S-NH_2_-LacNAc binding modes.

On the other hand, significant variation was observed in the orientation of the +1 GlcNAc residue. This residue was unresolved in previous structures, whereas here, clear electron density for GlcNAc was observed in the LacNAc, 3S-LacNAc, and 3S-Lewis^x^ complexes **(Figure 5a)**. In the 3S-Lewis^x^ structure, neither the α1,3-linked L-fucose substituent nor its linkage could be resolved, and in the 3S-Lewis^a^ structure, neither the β1,4-linked D-GlcNAc nor the α1,4-linked L-fucose could be resolved. The principal difference among the three resolved GlcNAc residues is the degree of rotation relative to Gal **(Figure 5a)**. The Gal–GlcNAc (C1–O–C4–C5) torsion angles are -102.4°, -116.8°, and - 159.1° for LacNAc, 3S-LacNAc, and 3S-Lewis^x^, respectively. In the LacNAc complex, this orientation may permit an interaction between the *N*-acetyl NH with Ser282 at approximately 2.9 Å. No other interactions with GlcNAc are observed, and the potential Ser282 interaction is progressively lengthened to approximately 3.4 and 5.9 Å in the 3S-LacNAc and 3S-Lewis^x^ complexes, respectively. These structures suggest that accommodation of the α1,3-linked L-fucose substituent of Lewis^x^ within BT1636^3S-Gal^ requires rotation of D-GlcNAc, whereas the D-GlcNAc and L-Fuc residues of Lewis^a^ are highly disordered. Despite this variation in the +1 subsite, the orientation and interactions of Gal, and its labile sulfate group, are conserved across all complexes.

### Carbohydrate sulfamates cannot be accommodated by the S1_20 enzyme BT1636^3S-Gal^

Because no complex of BT1636^3S-Gal^ with an aryl or carbohydrate sulfamate could be obtained (which is consistent with their reported lack of binding^20^), we performed MD simulations with 3S-LacNAc and its *O*3 sulfamate derivative (3SNH_2_-LacNAc). Docking followed by MD simulation positioned 3S-LacNAc in close alignment with the 3S-LacNAc and 3S-Lewis^X^ crystallographic structures, providing confidence in the model **(Figure 5b)**. The calculations indicated that binding is driven primarily by the 3S-Gal moiety, with GlcNAc making a smaller but significant contribution. Of the calculated binding free energy of -48.2 kcal mol^-1^, more than 30 kcal mol^-1^ was attributed to 3S-Gal, with coordination of the sulfate group to Ca^2+^ providing a major contribution **(Figure S7)**.

Replacement of the negatively charged sulfate with the larger, neutral sulfamate group causes 3SNH_2_-LacNAc to adopt a binding pose distinct from that of the native substrate **(Figure 5b)**. The altered binding pose in the MD simulation reduced the calculated binding free energy, from −48.2 kcal mol^-1^ for 3S-LacNAc to 3.3 kcal mol^-1^ for 3SNH_2_- LacNAc. This change reflects the loss of the large favourable contribution from the Ca^2+^- coordinated 3S-Gal sulfate group; instead, GlcNAc provides the principal contribution to binding **(Figure S7)**. Critically, in the calculated model, the sulfur atom of 3SNH_2_-LacNAc is positioned approximately 6 Å from Cβ of Ser77, compared with approximately 3 Å for 3S- LacNAc. This increased distance is incompatible with nucleophilic attack by the hydroxyl group of the FGly-gem diol **(Figure 5b)**. The change in binding pose is also accompanied by loss of the interactions between Gal *O*2 and Glu334 and Arg353, although His177, which recognises *O*4, retains a favourable contribution **(Figure S8)**. These findings provide insight that help to explain the inability of Gal *O*3-sulfamates to bind to or inhibit BT1636^3S-Gal^.

### Flexible sulfamate probes bind but do not covalently modify FGly in a carbohydrate sulfatase

Given the evidence that the conformational stringency of carbohydrate sulfate binding by the S1 sulfatase BT1636^3S-Gal^ prevents carbohydrate and aromatic sulfamates from adopting a reactive orientation within the sulfate-binding site, we postulated that presenting a sulfamate warhead on a small, flexible scaffold might engender a reactive binding pose. We therefore designed a focused library (compounds **1**–**13**, **Figure 6a**) in which we varied both the intrinsic reactivity of the sulfamate, parameterised by the p*K*_a_ value of the alcohol or *N*- hydroxy leaving group leaving group, and the structure of that leaving group, which may influence how the sulfamate is presented within the active site. Leaving-group p*K*_a_ values ranged from 6 to 17 and included simply alkyl (**1, 2**), fluoroalkyl (**3**–**6**) and *N*- hydroxysuccinimidyl (NHS) (**7**) sulfamates (**Figure 6a**). Presentation geometry was varied using a series of flexible, hydroxylated scaffolds (**8**–**13**) designed to mimic the hydroxyl groups and secondary sulfate of the natural substrate, 3-*O*-sulfate galactose, while providing conformational freedom unavailable to the more rigid sugar scaffold. These included the achiral secondary glycerol-2-sulfamate **10** and its primary regioisomer **11**.

**Figure 6.**
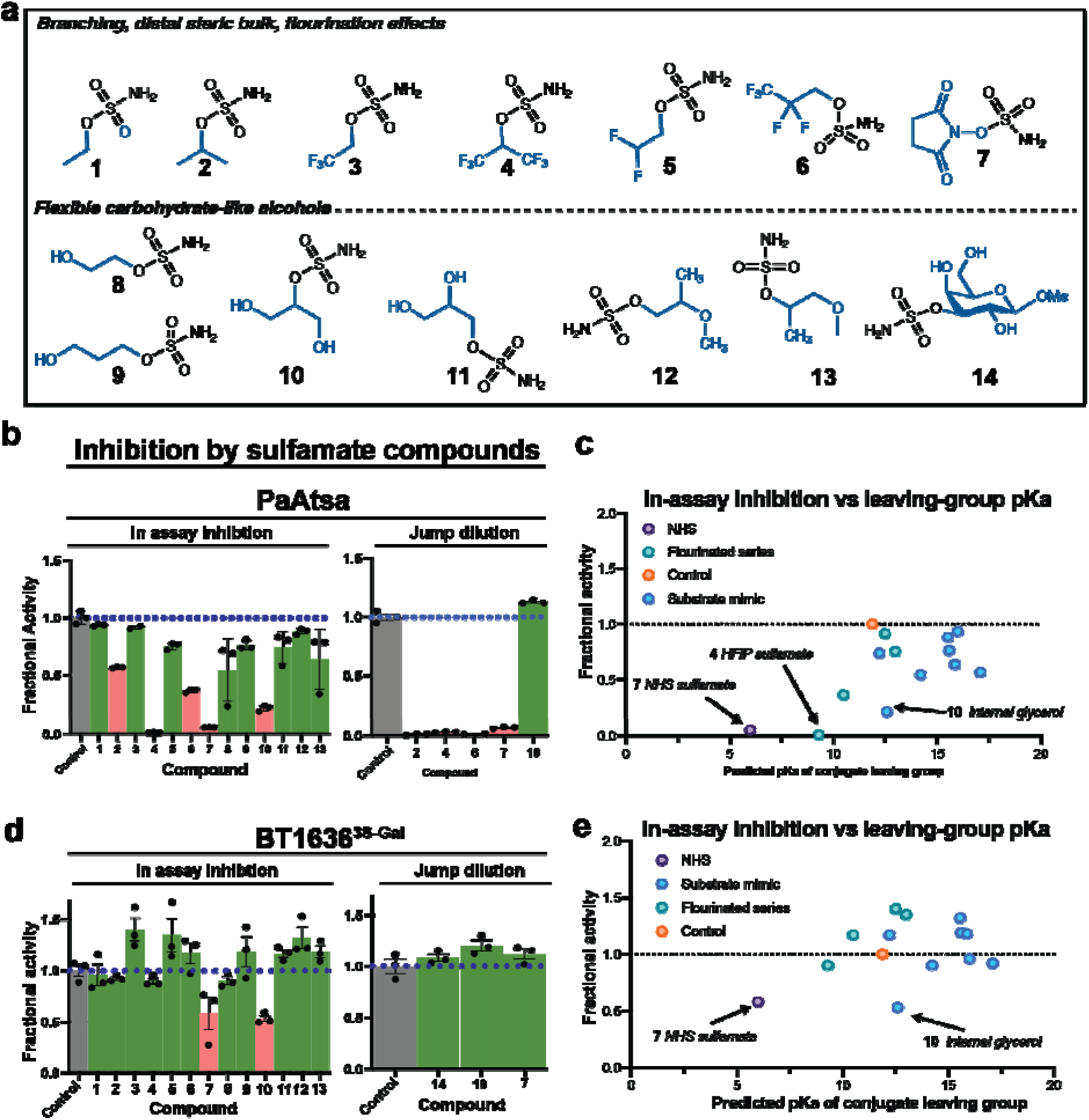
Flexible warhead-bearing scaffolds produce reversible inhibition of BT1636^3S-Gal^. **a.** Flexible scaffolds bearing sulfamate and their corresponding □-galactose derivative. **b.** Left, compounds were screened at 500 μM after overnight incubation. Green indicates little or no inhibition and red indicates a loss of activity. Right, PaAtsa (50 μM) was incubated overnight with **2, 4, 6, 7** and **10** (10 mM), then diluted 1000-fold before activity measurement. Assays contained 100 mM BTP, pH 7.5, 150 mM NaCl, 5 mM CaCl_2_ and 1 mM BODIPY-labelled 3S-Gal substrate. Data are means of triplicate assays; error bars represent SEM. **c.** Inhibition of PaAtsa from the initial screen plotted against the predicted p*K*_a_ of the conjugate acid of each sulfamate leaving group. **d.** Left, compounds were screened at 500 μM after overnight incubation. Green indicates little or no inhibition and red indicates a substantial loss of activity. Right, BT1636^3S-Gal^ (20 μM) was incubated overnight with **14**, **10** or **7** (10 mM), then diluted 1000-fold before activity measurement. Assays contained 100 mM BTP, pH 7.5, 150 mM NaCl, 5 mM CaCl₂ and 1 μM BODIPY-labelled 3S-Gal substrate. Data are means of triplicate assays; error bars represent SEM. **e.** Inhibition of BT1636^3S-Gal^ from the initial screen plotted against the predicted p*K*_a_ of the conjugate acid of each sulfamate leaving group.

We first tested the library against PaAtsA and observed that five compounds (**2, 4, 6, 7,** and **10**) caused 50%, or greater, inhibition in our in-assay format **(Figure 6b)**. Follow up jump dilution experiments, where PaAtsA was incubated with 10 mM compound overnight, then assayed by diluting 1000-fold into substrate solution, indicated **2, 4, 6,** and **7** were covalent sulfamate inhibitors whilst **10** functioned as a competitive inhibitor. Screening of the library against BT1636^3SGal^ identified two compounds with significant inhibition, glycerol-2- sulfamate **10** and NHS sulfamate **7** (**Figure 6c**). However, jump dilution experiments with BT1636^3SGal^, carried out as for PaAtsA above, fully restored activity (**Figure 6c**). Therefore, both compounds act as weak competitive inhibitors of BT1636^3SGal^ and do not form a persistent covalent adduct with the FGly.

## Discussion

Arylsulfamates are potent covalent inhibitors of S1 aryl and steroid sulfatases^14,16,27,33^, but the structure of the inhibited complex and the basis of its persistence have remained uncertain.^33,34^ Our crystallographic data identify the covalent complex formed in PaAtsA as a tetrahedral, *O*-linked α-hydroxysulfamate adduct of FGly51. Assignment of the linkage is supported by the electron-density maps, refinement behaviour and atomic B-factors, as well as by DFT calculations showing that the *O*-linked structure is more stable than the alternative *N*-linked isomer and more closely reproduces the experimental crystallographic geometry. These structures therefore provide direct evidence that arylsulfamates inhibit PaAtsA through covalent modification of the catalytic FGly residue. Interestingly, mass spectrometric analysis did not detect this adduct, suggesting that the α-hydroxysulfamate linkage is unstable under the conditions used for both peptidic digest and intact-protein mass spectrometry.

Bojarova *et al.* showed by linear free energy Brønsted analysis that ArO–S bond cleavage occurs during the first irreversible chemical step of inactivation. Several chemical mechanisms were proposed to account for these data, including an E1cb-type process involving the reactive intermediate sulfimine (HN=SO_2_) and concerted nucleophilic substitution at sulfur by an FGly hydroxyl group^27^. The *O*-linked structure observed here defines the product of covalent modification but does not establish whether a discrete intermediate is formed or whether bond cleavage and nucleophilic attack are concerted. Bojarova *et al.* also found that inactivation required 3 to 6 arylsulfamate molecules per enzyme molecule, with the higher stoichiometries observed for arylsulfamates containing better leaving groups^27^. They suggested that this behaviour might arise from non-adduct forming processes, such as E1cb elimination to form sulfenimine, which could diffuse from the active site before reacting with FGly. Thus, our structural results define the product of productive covalent modification but do not resolve the sequence of chemical events leading to its formation or the competing processes that determine inactivation efficiency.

Although arylsulfamates are generally described as irreversible sulfatase inhibitors, PaAtsA inactivated by 6-bromo-2-naphthyl sulfamate progressively recovered activity over several weeks. The *O*-linked α-hydroxysulfamate adduct can therefore undergo slow breakdown to regenerate active enzyme. Breakdown could occur either by elimination to regenerate the aldehyde form of FGly or by nucleophilic substitution at sulfur by water. The positioning of a water molecule above the sulfur centre in the PaAtsA adduct structure, together with the residual sulfatase activity of the C51S variant, is consistent with the possibility of nucleophilic substitution at sulfur.

DFT calculations predict high activation energy barriers for both pathways, consistent with the observed persistence of the adduct, but favour the two-step elimination pathway over direct substitution by water. It may well be that the sulfamate adduct lowers the rate of elimination to the point where hydrolysis becomes a less likely but competing mechanism for adduct turnover. Although high, the activation energies calculated here are consistent with those reported for ‘slow’ reactions. Callahan *et al*^35^ argue that activation enthalpies (ΔH^‡^) for the uncatalyzed hydrolysis of urea greater than about 20 kcal mol^-1^ correspond to ‘slow’ reactions (t_1/2_ between 33 to 1200 years) whereas the much faster urease catalysed reaction with t_1/2_ of 0.02 s has a ΔH^‡^ value of 10.5 kcal mol^-1^. Thus, the activation energy values found here for the degradation of the *O*-linked intermediate of PaAtsA are consistent with the process being ‘slow’, with the calculated fraction of molecules at 300 K (Boltzmann factor^36^) with a mean kinetic energy greater than 30 kcal mol^-1^ being 1.39 x 10^-22^, whereas at 10 kcal mol^-1^ this rises to a fraction of 5.18 x 10^-8^.

Whereas arylsulfamates are effective inhibitors of arylsulfatases such as PaAtsA, neither carbohydrate sulfamates and arylsulfamates covalently inactivate the bacterial S1 carbohydrate sulfatase BT1636^3S-Gal^. Gal *O*3-sulfamate does not inhibit BT1636^3S-Gal^, as observed here and previously by Crawford *et al.*^20^ A trisaccharide sulfamate has also been reported to inhibit the human S1_6 glycosaminoglycan *endo*-sulfatase Sulf1 competitively, but persistent covalent inhibition was not demonstrated^37,38^. Comparison of PaAtsA and BT1636^3S-Gal^ provides a structural explanation for these contrasting behaviours. In PaAtsA, replacement of sulfate by sulfamate removes the favourable Ca^2+^–oxyanion interaction. The MD simulations indicate that the sulfamate group rotates to avoid an unfavourable interaction between its NH_2_ group and the metal ion. The permissive hydrophobic pocket accommodates this rearrangement through changes in the orientation of the aromatic scaffold while maintaining the sulfur atom in a position compatible with attack by the FGly gem diol. PaAtsA therefore provides a model for how the flexible substrate-binding pockets of aryl- and steroid-recognising S1 sulfatases may tolerate the altered geometry of a sulfamate warhead.

By contrast, the crystal structures of BT1636^3S-Gal^ show highly conserved recognition of the *O*3-sulfated Gal residue across several glycan substrates, despite structural variation. MD simulations predict that replacing sulfate with sulfamate disrupts this conserved binding mode by eliminating the favourable contribution of Ca^2+^ coordination, The resulting rearrangement displaces the sulfur centre away from the catalytic nucleophile and prevents productive attack by FGly. A screen of alternative sulfamate scaffolds tested whether conformational freedom and variation of leaving group ability could overcome these geometric constraints. While many of these compounds were good inhibitors of the more permissive PaAtsA, and although glycerol 2-sulfamate **10** and NHS sulfamate **7** produced competitive inhibition of BT1636^3S-Gal^, activity was fully restored upon dilution, demonstrating that neither compound formed a persistent covalent adduct. Scaffold flexibility may therefore permit weak binding to this carbohydrate sulfatase but is insufficient to provide the combination of affinity and precise sulfamate positioning required for reaction with FGly in the carbohydrate sulfatase. These results suggest that a successful sulfamate inhibitor of a carbohydrate sulfatase must retain sufficient substrate-like interactions to bind productively while allowing the sulfamate group to adopt a geometry compatible with both the bound Ca^2+^ ion and nucleophilic attack.

This work identifies the long-lived product of arylsulfamate inactivation as an *O*-linked α-hydroxysulfamate adduct of FGly and shows that this adduct undergoes very slow breakdown, making its apparent irreversibility dependent on the timescale of observation. The contrasting behaviour of PaAtsA and BT1636^3S-Gal^ arises from differences in the conformational freedom permitted by their substrate-binding sites, with the permissive aryl- binding pocket of PaAtsA accommodating repositioning of the sulfamate that retains a productive trajectory for nucleophilic attack, whereas the stringent carbohydrate-recognition network of BT1636^3S-Gal^ does not. The failure of flexible sulfamate scaffolds to produce persistent inhibition further shows that conformational freedom alone is insufficient. Effective covalent inhibitors of carbohydrate sulfatases will require simultaneous optimisation of substrate-recognition interactions, sulfamate orientation relative to Ca^2+^ and the trajectory of attack by FGly.

---

**Figure S1. Activity of PaAtsA in the presence of sulfate and sulfamate ions and interaction distances of 4-phenylphenyl sulfate and 4-phenylphenyl sulfamate during molecular dynamics simulations.**

**a.** Activity of PaAtsA against 1 mM 4-nitrophenyl sulfate in the presence of increasing concentrations of sodium sulfate and sodium sulfamate. **b.** Distances of key interactions of 4-phenylphenyl sulfate over the course of the simulation.

**Figure S2. Torsion angles and MMPBSA free energy plot per residue for 4- phenylphenyl sulfate and 4-phenylphenyl sulfamate bound to PaAtsA.**

**a.** Time-dependent variation of the dihedral angles ϕ_1_, ϕ_2_, ϕ_3_, shown in the left, middle and right plots. (Top) Dihedral angles ϕ_1_, ϕ_2_, ϕ_3_ that characterise the conformation of PhPh-OSO_3_ (left), and PhPh-SO_2_NH_2_ (right) are defined. Sulfur, oxygen, nitrogen, and hydrogen atoms are represented by yellow, orange/magenta, red, blue, and white. Carbon atoms in PhPh- SO_3_ and PhPh-OSO_2_NH_2_ are represented in orange and magenta, respectively. **b.** Histogram of the Poisson Boltzmann free energy of binding per residue decomposition ΔG_i_^MMPBSA^ (kcal mol^-1^) when PhPh-OSO_3_ (left) and PhPh-OSO NH (right) are in bound state with PaAtsA. Contributions greater than 0.1 kcal mol^-1^ are reported.

**Figure S3. Activity and molecular mass of PaAtsA before and after treatment with Irosustat.**

**a.** Activity of PaAtsA against 1 mM 4-nitrophenyl sulfate with and without overnight incubation with 1 mM Irosustat. Buffer conditions were 10 mM HEPES pH 7.5 with 150 mM NaCl. Data are technical triplicates. **b.** LC-MS extracted ion chromatograms for 3+ charge states of formylglycine (*m/z* 506.910) and gem-diol (*m/z* 512.241) modified forms of trypsin-derived PaAtsA peptide LTDFHTASTGSPTR. **c.** Deconvoluted intact mass spectra obtained from non-treated (top) and Irosustat-treated (bottom) PaAtsA. The lower mass signal is consistent with loss of N-terminal Met and gem-diol conversion from FGly (theoretical mass = 61,962 Da). The higher mass signal is consistent with the same species plus a single O- gluconoylation within the protein sequence (theoretical mass = 62,140 Da).

**Figure S4. Interaction of PaAtsA with the coumarin leaving group of Irosustat and the activity of PaAtsA^C51S^ against 4-nitrophenyl sulfate and its covalent sulfate adduct.**

**a.** The interaction of the coumarin scaffold leaving group of Irosustat in the PaAtsA binding site. Protein is in green and the Irosustat coumarin scaffold in yellow. **b.** Activity of PaAtsA^C51S^ against 10 mM 4-nitrophenyl sulfate. It is calculated that each molecule of enzyme has turned over 40 molecules of 4-nitrophenyl sulfate over the time course. Reactions were carried out in triplicate in 10 mM HEPES pH 7.5 with 150 mM NaCl. **c.** Crystal structure of PaAtsA^C51S^ covalently bound in complex with sulfate from prolonged incubation with 4-nitrophenyl sulfate. The electron density map is the 2mFobs-DFc map contoured to 1 σ. Numbers are the B factors for the relevant oxygen atoms.

**Figure S5. Geometries, energies and key vibrational states along the proposed substitution pathway for breakdown of the covalent adduct.**

Carbon, oxygen, nitrogen, sulfur, calcium and hydrogen atoms are shown as grey, red, blue, yellow, gold, and white sticks, respectively. Structures represent the **(i)** reactant, **(ii)** transition state, **(iii)** apparent intermediate and **(iv)** product of the proposed substitution hydrolysis pathway. Straight and curved arrows denote selected interatomic distances and bond angles, respectively. Distances are given in Å and angles in degrees. For the reactant state, values derived from the crystallographic and DFT models are shown in blue and black, respectively.

**Figure S6. Geometries, energies and key vibrational states along the proposed elimination pathway for breakdown of the covalent adduct.**

Carbon, oxygen, nitrogen, sulfur, and calcium, and hydrogen atoms are shown as grey, red, blue, yellow, gold, and white sticks, respectively. Structures represent the **(i)** reactant, **(ii)** first transition state (TSE1), **(iii)** intermediate, **(iv)** second transition state (TSE2) and **(v)** product. Straight and curved arrows denote selected interatomic distances and bond angles, respectively. Distances are given in Å and angles in degrees. Values derived from the crystallographic and DFT models are shown in blue and black, respectively. Selected distances within the FGly–OSO_2_NH^2^ moiety are shown in bold in (iii) and (iv).

**Figure S7. MMPBSA Free energy plot per residue for 3S-LacNAc and 3SNH_2_-LacNAc bound to BT1636^3S-Gal^.**

Histogram of the Poisson Boltzmann free energy of binding per residue decomposition ΔG_i_^MMPBSA^ (kcal mol^-1^) when 3S-LacNAc. (left) and 3SNH_2_-LacNAc (right) are in bound state with BT1636^3S-Gal^. Contributions greater than 0.1 kcal mol^-1^ are reported.

**Figure S8. Distances of key interactions in the 3S-LacNAc and 3S-NH_2_-LacNAc complexes with BT1636^3S-Gal^.**

Distances between selected enzyme and ligand atoms were monitored during the simulations shown in **Figure 5b**. Black traces represent 3S-LacNAc and red traces represent 3S-NH_2_-LacNAc.

## Methods

### Synthesis of chemical compounds

The following compounds were prepared as described previously: 4-phenylphenyl sulfamate^39^, 6-bromo-2-napthyl sulfamate^20^ and methyl 3-O-sulfamoyl-β-D-galactoside^20^. Full details of the synthetic procedures for compounds synthesised for this work can be found in the Supporting Information.

### Recombinant protein production

Genes were amplified by PCR using the appropriate primers and cloned into pET28b with N- His_6_ using NheI/XhoI restriction sites. Recombinant proteins were expressed in *Escherichia coli* strains TUNER (DE3) (Novagen) and cultured to mid-exponential phase in LB supplemented with 50 μg/mL kanamycin at 37 °C and 180 rpm. Cells were then cooled to 16 °C, and gene expression was induced by the addition of 0.1 mM isopropyl β-D-1- thiogalactopyranoside; cells were then cultured for another 16 h at 16 °C and 180 rpm. The cells were then centrifuged at 5,000 × *g* for 10 mins and resuspended in 20 mM HEPES, pH 7.4, with 500 mM NaCl then were sonicated on ice. Recombinant protein was purified by immobilized metal ion affinity chromatography using a cobalt-based matrix (Talon, Takara Biosciences) and, after a wash with resuspension buffer, eluted with a step gradient of 10-, 50-, and 100-mM imidazole in resuspension buffer. Proteins were analysed by SDS-PAGE gel for purity. Proteins were then concentrated in centrifugal concentrators with a molecular mass cutoff of 30 kDa, loaded onto a 16/60 S200 Superdex size exclusion column, and eluted into 10mM HEPES pH 7.4 with 150mM NaCl. Fractions from this were then subject SDS-PAGE analysis and fractions judged to >95% pure pooled and again concentrated in centrifugal concentrators with a molecular mass cutoff of 30 kDa. Protein concentrations were determined by measuring absorbance at 280 nm using the molar extinction coefficient calculated by ProtParam on the ExPasy server (web.expasy.org/protparam/).

### HPAEC based enzyme desulfation assays

BT1636^3S-Gal^ (30 nM) was assayed against 1 μM BODIPY-labelled 3S-Gal as a fluorescent substrate. In initial screens, recombinant enzyme was incubated overnight with 1 mM inhibitor, before addition of substrate and subsequent following of reaction progress (final inhibitor concentration 500 μM). Jump dilution experiments were carried out by incubation of 20 μM enzyme with 10 mM inhibitor overnight, before a 500x dilution into the reaction (final protein 40 nM, inhibitor concentration 20 μM). Progress of the reactions were measured by ThermoScientific/Dionex ICS-6000 HPAEC with a Vanquish Fluorescence detector, with separation of sulfated substrate and unsulfated product over a Dionex CarboPac PA-200 3x50mm BioLC guard column at a flow rate of 1 ml/min for 3 minutes. The mobile phase of 100 mM sodium hydroxide with 0.2 M sodium acetate was increased linearly to 1 M sodium acetate over 0.25 min, held until 2 min when the column is re-equilibrated back into 100 mM sodium hydroxide with 0.2 M sodium acetate for the remainder of the run. Sampling was carried out eight times across a 24 h period (ThermoScientific/Dionex AS-AP autosampler), and relative peak area assessed using Chromeleon 7. GraphPad prism was used for subsequent data processing.

### PaAtsA desulfation assays

PaAtsA was assayed at 50 nM, with 1 mM 4-nitrophenyl sulfate as substrate in black wall/clear bottom microplates (Greiner). In initial reactions, PaAtsA was incubated overnight with 1mM inhibitor, before addition of substrate and monitoring of reaction progress (final inhibitor concentration 500 μM). Jump dilution experiments were carried out by incubation of 50 μM enzyme with 10 mM inhibitor overnight, before a 1000x dilution into the reaction (final protein 50 nM, inhibitor concentration 10 μM). The reaction progress was monitored continuously at 400 nm (BMG FLUOstar Omega), and linear rates calculated using GraphPad Prism.

### Irreversible inhibition constant calculations for PaAtsA

Irreversible inhibitors were assessed using a continuous method in which rate of inhibition was calculated from a reaction progress curve, according to protocols laid out by Mons *et al*.^40^ PaAtsA, 5 nM, was introduced to a solution containing 1 mM 4-nitrophenyl sulfate, and a varied molar quantity of the inhibitor to be tested. The reaction progress was monitored over 2 h following absorbance at 405 nm. The resulting curves were fit using **equation 1** (where F_t_ is observed signal, F_0_ is signal at t=0, v_i_ is initial velocity, and v_s_ is final velocity) to yield a *k*_obs_ value for each inhibitor concentration. Plots of *k*_obs_ vs [I] were fit to determine *K*_I_ and *k*_inact_.

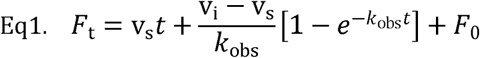

### X-ray crystallography experiments

Sparse matrix (JCSG & SG2, Molecular Dimensions) and systematic (PACT, Molecular Dimensions) screens were set up in 96-well sitting drop SwissCI MRC 2-drop plates (400-nL drops) using an SPT mosquito crystallisation robot. All proteins were in 10 mM HEPES pH 7.5, 150 mM NaCl. For the sulfamate adduct derived from Irosustat, co-crystallisation was carried out using PaAtsA at a final concentration of 29.4 mg/ml, with 2 mM irosustat and a final concentration of 2% DMSO. Large block form crystals were identified in a condition with 0.2 M MgCl_2_.6H_2_O, 0.1 M Tris pH 8.0 and 20% w/v PEG 6000, and cryo-protected with 20% PEG 200. For the sulfamate adduct derived from 6-bromonaphth-2-yl sulfamate PaAtsA was incubated with 6-bromonaphth-2-yl sulfamate for 16 h at room temp, followed by 48 h at 4°C, after IMAC purification. The protein was then subject to size exclusion before being concentrated to 20 mg/ml and crystallised in 20% PEG 6000, 0.1 M Tris pH 8.0 and 0.2 CaCl_2_; crystals were cryo-protected with 20% PEG 200. For the sulfate adduct derived from 4-nitrophenyl-sulfate, PaAtsA^C51S^ was incubated with 4-nitrophenyl-sulfate, after IMAC purification, for 16 h at room temp, followed by 48 h at 4 °C. The protein was then subject to size exclusion chromatography, then was concentrated to 20 mg/ml and crystallised in 20% w/v PEG 3,350 with 0.04 M citric acid, and 0.06 M BIS-TRIS propane, pH 6.4 and cryoprotected with 20% PEG 200. Wild-type variants of BT1636^3S-Gal^ were used that had a serine at the catalytic formylglycine position (*E. coli* cannot convert Ser to FGly). The protein was crystallised at 30 mg/ml in 40% 2-methyl-2,4-pentanediol (MPD), 5% PEG 8000 and 0.1 M sodium cacodylate, pH 6.5. Crystals were then serially soaked with 20 mM LacNAc, Lewis^a^, or Lewis^x^ dissolved in the reservoir solution overnight. For colonic mucin oligosaccharide soaks, treatment was the same but 20 mg/ml was used as the soaking concentration. No additional cryoprotection was needed as it was afforded by the crystallisation condition. Data were collected at Diamond Light Source (Oxford) on beamlines I0-3 at 100 K. The data were integrated and scaled with auto processing software at Diamond via DIALS, Xia2 or autoproc with staraniso pipelines and merged with Aimless^41,42^. Five percent of observations were randomly selected for the R_free_ set. The phase problem was solved by molecular replacement using the program Phaser or Molrep with using the PDB 1HDH for PaAtsA and 7AN1 for BT1636^3S-Gal^. Models then underwent recursive cycles of model building in Coot^43^ and refinement cycles in Refmac5^44^. The models were validated using Coot^43^ and MolProbity^45^. Carbohydrates were made using Jigand^46^. Structure Figures were made using PyMOL (The PyMOL Molecular graphics system, Version 2.0 Schrodinger, LLC.) and all other programs used were from the CCP4 suite^47^. The data processing and refinement statistics are reported in **(Supplementary tables S4 and S5)**

### Proteomic Analysis

Protein solution was diluted 1:1 (v:v) with aqueous 100 mM ammonium bicarbonate. Proteolytic digestion was performed with the addition of 0.2 μg of sequencing grade trypsin (V5111 – Promega) and incubation at 37 °C for 16 h. Resulting peptides were loaded onto EvoTip Pure tips for desalting and as a disposable trap column for nanoUPLC using an EvoSep One system. A pre-set EvoSep 100 SPD gradient was used with an 8 cm EvoSep C_18_ Performance column (8 cm x 150 μm x 1.5 μm). The nanoUPLC system was interfaced to a timsTOF HT mass spectrometer (Bruker) with a CaptiveSpray ionisation source (Source). Positive PASEF-DDA, ESI-MS and MS^2^ spectra were acquired using Compass HyStar software (version 6.2, Bruker). Instrument source settings were: capillary voltage, 1,600 V; dry gas, 3 l/min; dry temperature; 180 °C. Spectra were acquired between *m/z* 100- 1,700. TIMS settings were: 1/K0 0.6-1.60 V.s/cm^2^; Ramp time, 100 ms; Ramp rate 9.42 Hz. Data dependant acquisition was performed with 10 PASEF ramps and a total cycle time of 1.17 s. An intensity threshold of 2,500 and a target intensity of 20,000 were set with active exclusion applied for 0.4 min post precursor selection. Collision energy was interpolated between 20 eV at 0.6 V.s/cm^2^ to 59 eV at 1.6 V.s/cm^2^. Data were searched using FragPipe (v22.0) against the expected amino acid sequence of PaAtsA allowing for variable modification of Met by oxidation plus wildcard mass addition within the range of -1 Da to +500 Da. A 20 ppm error tolerance was set for both MS1 and MS2. A 1% FDR cut-off was applied to peptide matches. Detection of formylglycine +27.995 Da and gem-diol + 43.999 Da was observed within the expected peptide sequence LTDFHTASTGSPTR. Protein N-terminal gluconoylation (+178.048 Da) on the peptide GSSHHHHHHSSGLVPR was also identified. Extracted ion chromatograms for identified peptidoforms of LTDFHTASTGSPTR were obtained using Bruker DataAnalysis (v6.0).

### Intact Protein Mass Spectrometry

Samples were diluted 10-fold in 0.1% trifluoroacetic acid and eluted from a MSPac DS-10 Desalting Cartridge at a flow rate of 0.2 mL/min (Thermo Scientific), held at 30 °C, driven by an IClass UPLC (Waters Corp.). Mass acquisition was via an Orbitrap Fusion Tribrid mass spectrometer (Thermo Scientific) fitted with a heated electrospray ionization source. The spray voltage was held static at 3500 V with the sheath gas set at 25 (arbitrary units) and the auxiliary gas at 5. The sweep gas was set to 0. Ion transfer tube temperature was maintained at 300 °C with the application mode configured for Intact Proteins. Measurements were carried out in the Orbitrap at a resolution of 7500. The scan range was *m/z* 800-3000 with an AGC target of 1.2 × 10^6^. The RF lens was set to 150% and source fragmentation was enabled at 60 V. Acquired spectra were de-convoluted to average neutral masses using Thermo BioPharma Finder software (v5.1) implementing the ReSpect algorithm.

### Molecular Modelling analysis

#### Conformational analysis of 4-Phenyl-phenyl sulfate, and 4-Phenyl-phenyl sulfamate

4-Phenyl-phenyl sulfate (PhPh-OSO_3_) and the 4-phenyl-phenyl sulfamate (PhPh-OSO_2_NH_2_) were obtained by molecular editing using Maestro release 2023 (Schrödinger Inc.). Both compounds were analysed for the prediction of their lowest energy conformation. PhPh- OSO_3_ is characterized by three principal degrees of freedom (dihedral angles) φ_1_ = O_24_-S_23_- O_22_-C_5_, φ_2_ = S_23_-O_22_-C_5_-C_4_, φ_3_ = C_1_-C_2_-C_11_-C_12_, while PhPh-OSO_2_NH_2_ is described by four dihedrals: φ_0_ = H_27_-N_26_-S_23_-O_22_, φ_1_ = N_26_-S_23_-O_22_-C_5_, φ_2_ = S_23_-O_22_-C_5_-C_4_, φ_3_ = C_1_-C_2_-C_11_-C_12_, according to the numbering system reported in **Figure S2**. The most stable conformations calculated at the quantum-mechanic (QM) level of theory HF/6-31G* gives φ_1_ = 59.6°, φ_2_ = 88.1°, φ_3_ = 136.2°, with a total energy E^HF/6-31G*^ + E^PZ^ = -1156.4220 hartree, and φ_0_ = 113.0°, φ_1_ = 180.0°, φ_2_ = 91.0°, φ_3_ = 134.1°, with a total energy (including the zero-point correction) E^HF/6-31G*^ + E^PZ^ = -1137.0712 hartree, for PhPh-OSO_3_ and PhPh-OSO_2_NH_2_, respectively. All these conformations are verified to be minima since all the calculated vibrational frequencies corresponds to real numbers. The Gaussian16 program was used for this QM analysis^48^. Point-like RESP charges were calculated for these optimized conformations of PhPh-OSO_3_ and PhPh-OSO_2_NH_2_ in accordance with the standard required by the Amber force-field. The Amber atom-types (parm10.dat) were selected for both these compounds using the antechamber application included in Ambertools 18.0^49^. The optimized conformations of these compounds were then submitted to molecular docking.

#### Molecular Docking of PhPh-OSO_3_, PhPh-OSO_2_NH_2_ on Arylsulfatase PaAtsA

Calculated geometries of the complexes between PhPh-OSO_3_, PhPh-OSO_2_NH_2_ and PaAtsA (PDB ID 1HDH) were obtained using automatic docking (Autodock 4.0)^50^. Since FGly51 is not recognized by the Amber Force Field, it was replaced at this stage by S51. The parameters of ligands (PhPh-OSO_3_, PhPh-OSO_2_NH_2_) and receptor (PaAtsA) were prepared using AutoDockTools4^50^, while Gesteiger charges were used for both ligands and receptor in these docking simulations^51^. The grid-box was centred on the Cε1 of His211, with dimensions of 80 × 80 × 80 grid-points. The genetic algorithm was used in all the docking runs, with parameters: *number of GA runs*, *population size*, *max number of energy evaluation*, and *max number of generations* set as 100, 20000, 5 × 10^7^, 600,000, respectively. At each run the docking solutions were clustered using a cut-off RMSD = 2.0 Å from the cluster centroid. The selected poses are characterized by the highest score, and their position between the residues: S51, E74, K113, H115, I156, T160, H211, W212, F331 (active site of PaAtsA). Two clusters characterized by population 85 and 15 poses and average binding energy -6.69 and -6.58 kcal mol^-1^, respectively were obtained docking PhPh-OSO_3_ on PaAtsA. Analogously, two clusters with population 99 and 1 and average binding energy -6.76 and -6.29 kcal mol^-1^, respectively, were generated upon docking PhPh- OSO_2_NH_2_ on PaAtsA. Poses that show the lowest binding energy: -6.91 and -7.25 kcal mol^-1^ and belong to the most populated clusters were selected to represent PhPh-OSO_3_ – PaAtsA, and PhPh-OSO_2_NH_2_ – PaAtsA complexes, respectively. The selected poses are also characterized by a suitable position of PhPh-OSO_3_ and PhPh-OSO_2_NH_2_ in the active site PaAtsA (**Table S3**), and were submitted to MD simulation in explicit solvent as described in the former paragraph.

### Quantum mechanical (QM) modelling

Reduced models of the O-linked and N-linked intermediates of PaAtsA were built from the corresponding X-ray resolved models and selecting all the residues that are within 3.0 Å from the FGly51-Cβ-Oγ_1_-O-SO_2_, or FGly51-Cβ-NH-SO_3_ moieties, respectively, as well as the amino acids that are involved in the TE mechanism, according with Marino T. *et al*.^11^. Therefore, both these reduced models include the Ca^+2^, the Cα (replaced by methyl group) and the atoms of the side chains of the following amino acids: Asp13, Asp14, FGly51, Lys113, His115, His211, Asp317, Asn318, Glu321, Lys375, as well as two co-crystallized water molecules. The molecular editing and the inclusion of the missing hydrogen atoms were done using Pymol 2.5.5. and Gaussview 3.07. To characterize both substitution hydrolysis and the elimination pathways, Asp317 and H211, were described in their protonated and anionic states, respectively, in both models. Upon editing, these reduced models of O-linked and N-linked intermediates of PaAtsA include 147 and 146 atoms, respectively, and are characterized by a total charge of 0 and -1, in that order, while their multiplicity is 1.

The geometry of *O*-linked and *N*-linked intermediate models of PaAtsA were then optimized using the DFT wavefunction with the B3LYP functional; a split-valence basis set was used that includes the triple-zeta 6-311+G(2d) for Ca^+2^ and S, and the double-zeta 6- 31+G(d) for H, C, N, O. The superfine integration grid (keyword Int=superfineGrid) was used in all the calculations. The search for minimum energy geometry points was done using the keyword ‘Optimization’ and stopping it when default threshold conditions on forces and step size were satisfied; the search for transition states (first order saddle points) were done stopping the optimization when very tight threshold conditions were satisfied. All the stationary point geometries explored, being minimum or transition states (TS), were verified by calculating the vibrational frequencies. The total internal energy of the explored geometries being minimum (stable complex), or transition state are calculated summing the electronic energy (E^B3LYP^) with the fundamental vibrational state (zero-point energy, E^ZP^). The Gaussian 16 program was used for all the QM calculations. The comparison of the internal energy content indicated that the O-linked is thermodynamically more stable than the N-linked intermediate (see result section).

To predict the mechanism by which the *O*-linked intermediate is degraded, according to the second step of the TE reaction (**Figure 1a**), the thermodynamic profile of the substitution hydrolysis and elimination pathways were determined, and reported in **Figure 4b**, and **Figure 4c**, respectively. The transition states of the substitution hydrolysis and elimination pathways were determined by manual editing of their initial conformation from of the optimized geometry of the O-linked PaAtsA intermediate and running a TS search according to the Optimization keyword in Gaussian 16. In all cases the TS geometry search was stopped until very tight conditions on forces and step size were satisfied, and the TS was confirmed by the existence of the one imaginary frequency (**Table S1**, **Table S2**). One TS and two TSs (TSE1, TSE2) were found in the substitution and elimination mechanisms, respectively, and reported in **Figure 4b, Figure S5(ii)**, and **Figure 4c**, **Figure S6(ii, iv)**, respectively. The DFT electronic energy (E^B3LYP^), zero-point energy (E^ZP^), internal energy (E^B3LYP+ZP^), and the frequency of the fundamental vibration (ν_0_), are reported in **TableS1**, and **TableS2** for the reactant, TS, intermediate and product of the substitution-hydrolysis and elimination mechanism, respectively. Starting from the TS of the substitution-hydrolysis (**Figure S5(ii)**), an IRC calculation in forward and reverse direction (separately) was done to verify that a minimum energy path connects this TS with the corresponding reactant and product states, that precede and follow it, respectively (**Figure S5**). In this case the forward IRC stops at a geometry that is intermediate between the TS and the hypothesised minimum (apparent intermediate), in fact the sulfur atom is between the tetrahedral and the penta-coordinated geometry, which correspond to an energy minimum (**Figure S5 (iii)**). Furthermore, re-optimizing the geometry of this apparent intermediate state upon moving one of the NH_2_- proton in the middle between the Oγ_1_ of FGly and the NH- moiety of the sulfamate group, the system converge to a minimum energy state that in our opinion correspond to a reasonable product state of the substitution-hydrolysis (**Figure S5 (iv)**). In fact, at best of our investigation, no transition state was found that connect this apparent intermediate (**Figure S5 (iii)**) with the product state (**Figure S5 (iv)**). Analogously, starting from the TSE1 and TSE2 that characterize the 1^st^ and 2^nd^ step in which the elimination mechanism decomposes, two IRC calculations were done: the former was found connecting the TSE1 with the reactant state (reverse) and with the intermediate state (forward). The latter IRC was found connecting the TSE2 with the intermediate state (reverse) and with the product state (forward); these sampled geometries are reported in **Figure S6 (i)-(v)**. All these IRC calculations were done using the following keywords: IRC=(RCFC, LQA, StepSize=17, RecalcFC=(Predictor=50), MaxCycle=200).

## Data availability statement

The crystal structure datasets generated have been deposited in the in the Protein Data Bank (PDB) under the following accession numbers: 33HO/pdb_000033ho, 33HP/pdb_000033hp, 33IL/pdb_000033il, 33HR/pdb_000033hr, 33IO/pdb_000033io, 33ID/pdb_000033id, and 33II/pdb_000033ii. Information on all other data and materials are contained within the main manuscript and Supplemental Information.

## Code availability statement

No new code was developed as part of this project

## Acknowledgements

We thank the York Structural Biology Laboratory X-ray Facility for access to structural infrastructure, and Dr Johan Turkenburg and Mr Sam Hart for assistance with sample preparation, diffraction screening, and data collection. We gratefully acknowledge Diamond Light Source for time on beamlines I03 and I04, under proposal MX39189. Computational work used the Viking Cluster, a high-performance computing facility provided by the University of York, and we gratefully acknowledge support from University of York IT Services and the Research IT team. We thank Dr Miloš Hricovíni (Institute of Chemistry, Slovak Academy of Sciences) for valuable and insightful discussions, and Dr Rosachiara Salvino (Istituto di Ricerche Chimiche e Biochimiche G. Ronzoni) for helpful discussions.

## Author contributions

CWET performed the enzymology. CWET and AC performed X-ray crystallography experiments and analysis. SE carried out molecular dynamics and density functional theory calculations. ZC, MBS, MN, LIW, CJC carried out chemical synthesis. AD, CT, and MAF carried out mass spectrometry analyses. EAY, SJW and AC devised the original drafts. All authors read and approved the manuscript.

## References

1 Dias, I. H. K. et al. Sulfate-based lipids: Analysis of healthy human fluids and cell extracts. Chem Phys Lipids 221, 53–64 (2019). 10.1016/j.chemphyslip.2019.03.009

2 Cole, G. B. et al. Specific estrogen sulfotransferase (SULT1E1) substrates and molecular imaging probe candidates. Proceedings of the National Academy of Sciences of the United States of America 107, 6222–6227 (2010). 10.1073/pnas.0914904107

3 Stewart, V. & Ronald, P. C. Sulfotyrosine residues: Interaction specificity determinants for extracellular protein-protein interactions. The Journal of biological chemistry 298, 102232 (2022). 10.1016/j.jbc.2022.102232

4 Luis, A. S. et al. Sulfated glycan recognition by carbohydrate sulfatases of the human gut microbiota. Nature chemical biology 18, 841–849 (2022). 10.1038/s41589-022-01039-x

5 Mistry, R. et al. Polysaccharide sulfotransferases: the identification of putative sequences and respective functional characterisation. Essays Biochem 68, 431–447 (2024). 10.1042/EBC20230094

6 Barbeyron, T. et al. Matching the Diversity of Sulfated Biomolecules: Creation of a Classification Database for Sulfatases Reflecting Their Substrate Specificity. PloS one 11, e0164846 (2016). 10.1371/journal.pone.0164846

7 Stam, M. et al. SulfAtlas, the sulfatase database: state of the art and new developments. Nucleic Acids Res 51, D647–D653 (2023). 10.1093/nar/gkac977

8 Hanson, S. R., Best, M. D. & Wong, C. H. Sulfatases: structure, mechanism, biological activity, inhibition, and synthetic utility. Angewandte Chemie 43, 5736–5763 (2004). 10.1002/anie.200300632

9 Appel, M. J. et al. Formylglycine-generating enzyme binds substrate directly at a mononuclear Cu(I) center to initiate O2 activation. Proceedings of the National Academy of Sciences of the United States of America 116, 5370–5375 (2019). 10.1073/pnas.1818274116

10 Berteau, O., Guillot, A., Benjdia, A. & Rabot, S. A new type of bacterial sulfatase reveals a novel maturation pathway in prokaryotes. The Journal of biological chemistry 281, 22464–22470 (2006). 10.1074/jbc.M602504200

11 Marino, T., Russo, N. & Toscano, M. Catalytic mechanism of the arylsulfatase promiscuous enzyme from Pseudomonas aeruginosa. Chemistry 19, 2185–2192 (2013). 10.1002/chem.201201943

12 Hernandez-Guzman, F. G., Higashiyama, T., Pangborn, W., Osawa, Y. & Ghosh, D. Structure of human estrone sulfatase suggests functional roles of membrane association. The Journal of biological chemistry 278, 22989–22997 (2003). 10.1074/jbc.M211497200

13 Reed, M. J., Purohit, A., Woo, L. W., Newman, S. P. & Potter, B. V. Steroid sulfatase: molecular biology, regulation, and inhibition. Endocr Rev 26, 171–202 (2005). 10.1210/er.2004-0003

14 Thomas, M. P. & Potter, B. V. Estrogen O-sulfamates and their analogues: Clinical steroid sulfatase inhibitors with broad potential. J Steroid Biochem Mol Biol 153, 160–169 (2015). 10.1016/j.jsbmb.2015.03.012

15 Stanway, S. J. et al. Steroid sulfatase: a new target for the endocrine therapy of breast cancer. Oncologist 12, 370–374 (2007). 10.1634/theoncologist.12-4-370

16 Bojarova, P. & Williams, S. J. Aryl sulfamates are broad spectrum inactivators of sulfatases: effects on sulfatases from various sources. Bioorg Med Chem Lett 19, 477–480 (2009). 10.1016/j.bmcl.2008.11.059

17 Palmieri, C. et al. IPET study: an FLT-PET window study to assess the activity of the steroid sulfatase inhibitor irosustat in early breast cancer. Breast Cancer Res Treat 166, 527–539 (2017). 10.1007/s10549-017-4427-x

18 Palmieri, C. et al. IRIS study: a phase II study of the steroid sulfatase inhibitor Irosustat when added to an aromatase inhibitor in ER-positive breast cancer patients. Breast Cancer Res Treat 165, 343–353 (2017). 10.1007/s10549-017-4328-z

19 Pautier, P. et al. A Phase 2, Randomized, Open-Label Study of Irosustat Versus Megestrol Acetate in Advanced Endometrial Cancer. Int J Gynecol Cancer 27, 258–266 (2017). 10.1097/IGC.0000000000000862

20 Crawford, C. J. et al. Arylsulfamates inhibit colonic Bacteroidota growth through a sulfatase-independent mechanism. Proceedings of the National Academy of Sciences of the United States of America 122, e2414331122 (2025). 10.1073/pnas.2414331122

21 Png, C. W. et al. Mucolytic bacteria with increased prevalence in IBD mucosa augment in vitro utilization of mucin by other bacteria. Am J Gastroenterol 105, 2420–2428 (2010). 10.1038/ajg.2010.281

22 Tsai, H. H., Dwarakanath, A. D., Hart, C. A., Milton, J. D. & Rhodes, J. M. Increased faecal mucin sulphatase activity in ulcerative colitis: a potential target for treatment. Gut 36, 570–576 (1995). 10.1136/gut.36.4.570

23 Chatzidaki-Livanis, M. & Comstock, L. E. Friend turned foe: a role for bacterial sulfatases in colitis. Cell host & microbe 17, 540–541 (2015). 10.1016/j.chom.2015.04.012

24 Corfield, A. P. et al. Colonic mucins in ulcerative colitis: evidence for loss of sulfation. Glycoconj J 13, 809–822 (1996). 10.1007/BF00702345

25 Hickey, C. A. et al. Colitogenic Bacteroides thetaiotaomicron Antigens Access Host Immune Cells in a Sulfatase-Dependent Manner via Outer Membrane Vesicles. Cell host & microbe 17, 672–680 (2015). 10.1016/j.chom.2015.04.002

26 Bloom, S. M. et al. Commensal Bacteroides species induce colitis in host- genotype-specific fashion in a mouse model of inflammatory bowel disease. Cell host & microbe 9, 390–403 (2011). 10.1016/j.chom.2011.04.009

27 Bojarova, P. et al. Direct evidence for ArO-S bond cleavage upon inactivation of Pseudomonas aeruginosa arylsulfatase by aryl sulfamates. Chembiochem : a European journal of chemical biology 9, 613–623 (2008). 10.1002/cbic.200700579

28 von Bulow, R., Schmidt, B., Dierks, T., von Figura, K. & Uson, I. Crystal structure of an enzyme-substrate complex provides insight into the interaction between human arylsulfatase A and its substrates during catalysis. Journal of molecular biology 305, 269–277 (2001). 10.1006/jmbi.2000.4297

29 Boltes, I. et al. 1.3 A structure of arylsulfatase from Pseudomonas aeruginosa establishes the catalytic mechanism of sulfate ester cleavage in the sulfatase family. Structure 9, 483–491 (2001). 10.1016/s0969-2126(01)00609-8

30 Miller, B. R., 3rd et al. MMPBSA.py: An Efficient Program for End-State Free Energy Calculations. J Chem Theory Comput 8, 3314–3321 (2012). 10.1021/ct300418h

31 Srinivasan, J., Miller, J., Kollman, P. A. & Case, D. A. Continuum solvent studies of the stability of RNA hairpin loops and helices. J Biomol Struct Dyn 16, 671–682 (1998). 10.1080/07391102.1998.10508279

32 Luis, A. S. et al. A single sulfatase is required to access colonic mucin by a gut bacterium. Nature 598, 332–337 (2021). 10.1038/s41586-021-03967-5

33 Potter, B. V. L. SULFATION PATHWAYS: Steroid sulphatase inhibition via aryl sulphamates: clinical progress, mechanism and future prospects. J Mol Endocrinol 61, T233–T252 (2018). 10.1530/JME-18-0045

34 Williams, S. J., Denehy, E. & Krenske, E. H. Experimental and theoretical insights into the mechanisms of sulfate and sulfamate ester hydrolysis and the end products of type I sulfatase inactivation by aryl sulfamates. The Journal of organic chemistry 79, 1995–2005 (2014). 10.1021/jo4026513

35 Callahan, B. P., Yuan, Y. & Wolfenden, R. The burden borne by urease. J Am Chem Soc 127, 10828–10829 (2005). 10.1021/ja0525399

36 Reif, F. Fundamentals of Statistical and Thermal Physics. 651 (1965).

37 Chiu, L. T. et al. Trisaccharide Sulfate and Its Sulfonamide as an Effective Substrate and Inhibitor of Human Endo-O-sulfatase-1. J Am Chem Soc 142, 5282–5292 (2020). 10.1021/jacs.0c00005

38 Schelwies, M. et al. Glucosamine-6-sulfamate analogues of heparan sulfate as inhibitors of endosulfatases. Chembiochem : a European journal of chemical biology 11, 2393–2397 (2010). 10.1002/cbic.201000401

39 Winum, J. Y. et al. Carbonic anhydrase inhibitors. Inhibition of cytosolic isozymes I and II and transmembrane, tumor-associated isozyme IX with sulfamates including EMATE also acting as steroid sulfatase inhibitors. Journal of medicinal chemistry 46, 2197–2204 (2003). 10.1021/jm021124k

40 Mons, E., Roet, S., Kim, R. Q. & Mulder, M. P. C. A Comprehensive Guide for Assessing Covalent Inhibition in Enzymatic Assays Illustrated with Kinetic Simulations. Curr Protoc 2, e419 (2022). 10.1002/cpz1.419

41 Evans, P. Scaling and assessment of data quality. Acta crystallographica. Section D, Biological crystallography 62, 72–82 (2006). 10.1107/S0907444905036693

42 Evans, P. R. An introduction to data reduction: space-group determination, scaling and intensity statistics. Acta crystallographica. Section D, Biological crystallography 67, 282–292 (2011). 10.1107/S090744491003982X

43 Emsley, P., Lohkamp, B., Scott, W. G. & Cowtan, K. Features and development of Coot. Acta crystallographica. Section D, Biological crystallography 66, 486–501 (2010). 10.1107/S0907444910007493

44 Murshudov, G. N. et al. REFMAC5 for the refinement of macromolecular crystal structures. Acta crystallographica. Section D, Biological crystallography 67, 355–367 (2011). 10.1107/S0907444911001314

45 Chen, V. B. et al. MolProbity: all-atom structure validation for macromolecular crystallography. *Acta crystallographica. Section D*, Biological crystallography 66, 12–21 (2010). 10.1107/S0907444909042073

46 Lebedev, A. A. et al. JLigand: a graphical tool for the CCP4 template-restraint library. Acta crystallographica. Section D, Biological crystallography 68, 431–440 (2012). 10.1107/S090744491200251X

47 Collaborative Computational Project, N. The CCP4 suite: programs for protein crystallography. Acta crystallographica. Section D, Biological crystallography 50, 760–763 (1994). 10.1107/S0907444994003112

48 Frisch, M. J. T., G. W.; Schlegel, H. B.; Scuseria, G. E.; Robb, M. A.; Cheeseman, J. R.; Scalmani, G.; Barone, V.; Petersson, G. A.; Nakatsuji, H.; Li, X.; Caricato, M.; Marenich, A. V.; Bloino, J.; Janesko, B. G.; Gomperts, R.; Mennucci, B.; Hratchian, H. P.; Ortiz, J. V.; Izmaylov, A. F.; Sonnenberg, J. L.; Williams-Young, D.; Ding, F.; Lipparini, F.; Egidi, F.; Goings, J.; Peng, B.; Petrone, A.; Henderson, T.; Ranasinghe, D.; Zakrzewski, V. G.; Gao, J.; Rega, N.; Zheng, G.; Liang, W.; Hada, M.; Ehara, M.; Toyota, K.; Fukuda, R.; Hasegawa, J.; Ishida, M.; Nakajima, T.; Honda, Y.; Kitao, O.; Nakai, H.; Vreven, T.; Throssell, K.; Montgomery, J. A., Jr.; Peralta, J. E.; Ogliaro, F.; Bearpark, M. J.; Heyd, J. J.; Brothers, E. N.; Kudin, K. N.; Staroverov, V. N.; Keith, T. A.; Kobayashi, R.; Normand, J.; Raghavachari, K.; Rendell, A. P.; Burant, J. C.; Iyengar, S. S.; Tomasi, J.; Cossi, M.; Millam, J. M.; Klene, M.; Adamo, C.; Cammi, R.; Ochterski, J. W.; Martin, R. L.; Morokuma, K.; Farkas, O.; Foresman, J. B.; Fox, D. J. Gaussian, Inc., Wallingford CT. Gaussian 16, Revision C.01. (2016).

49 Case, D. A. et al. AmberTools. J Chem Inf Model 63, 6183–6191 (2023). 10.1021/acs.jcim.3c01153

50 Morris, G. M. et al. AutoDock4 and AutoDockTools4: Automated docking with selective receptor flexibility. J Comput Chem 30, 2785–2791 (2009). 10.1002/jcc.21256

51 Gasteiger, J., Marsili, M. Iterative partial equalization of orbital electronegativity—a rapid access to atomic charges. Tetrahedron 36, 3219– 3228 (1980). 10.1016/0040-4020(80)80168-2

