## Supplemental information for "Structural basis for covalent inhibition of sulfatases by sulfamate warheads"

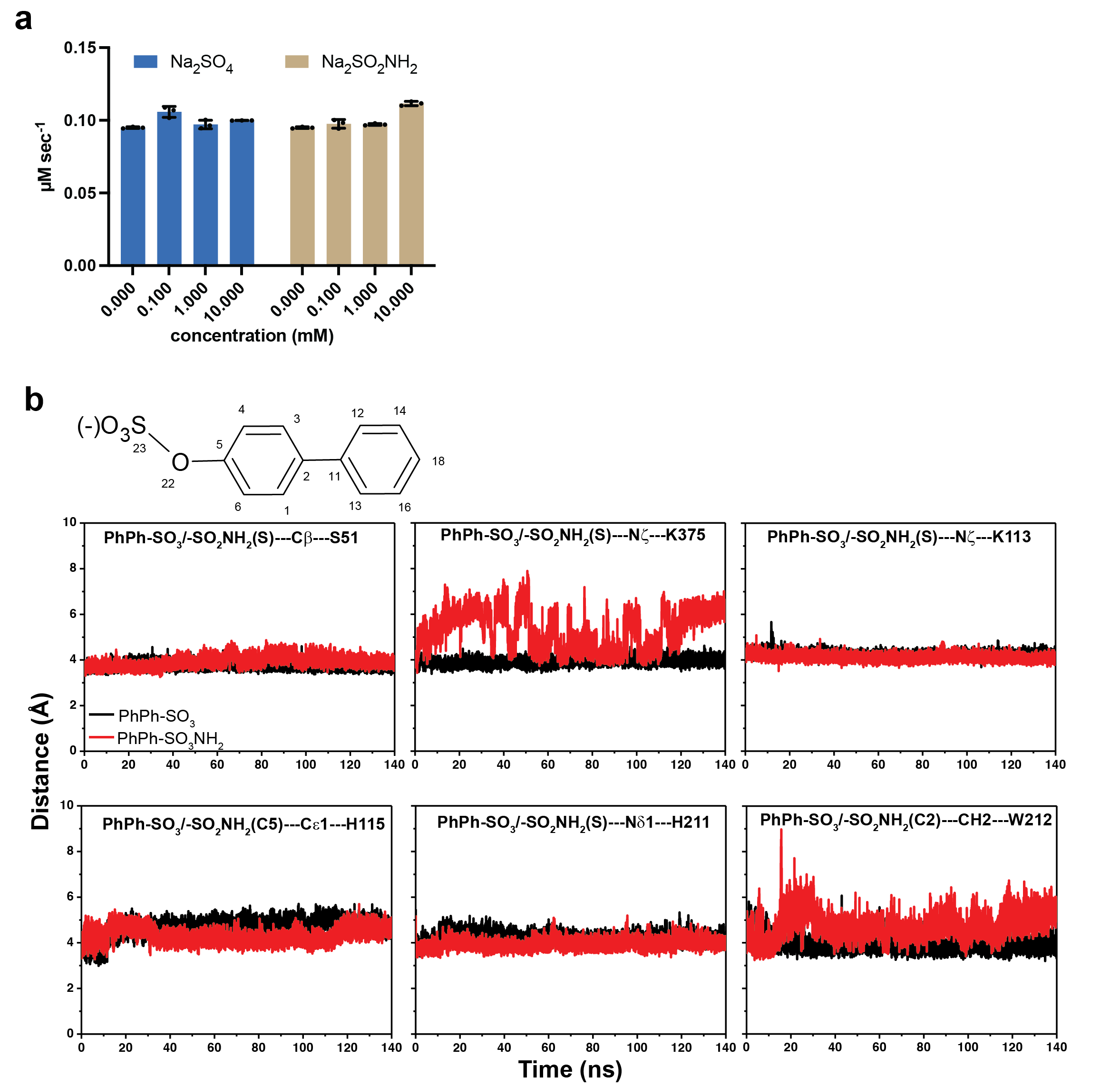

**Figure S1. Activity of PaAtsA in the presence of sulfate and sulfamate ions and interaction distances of 4-phenylphenyl sulfate and 4-phenylphenyl sulfamate during molecular dynamics simulations.**

**a.** Activity of PaAtsA against 1 mM 4-nitrophenyl sulfate in the presence of increasing concentrations of sodium sulfate and sodium sulfamate. **b.** Distances of key interactions of 4-phenylphenyl sulfate over the course of the simulation.

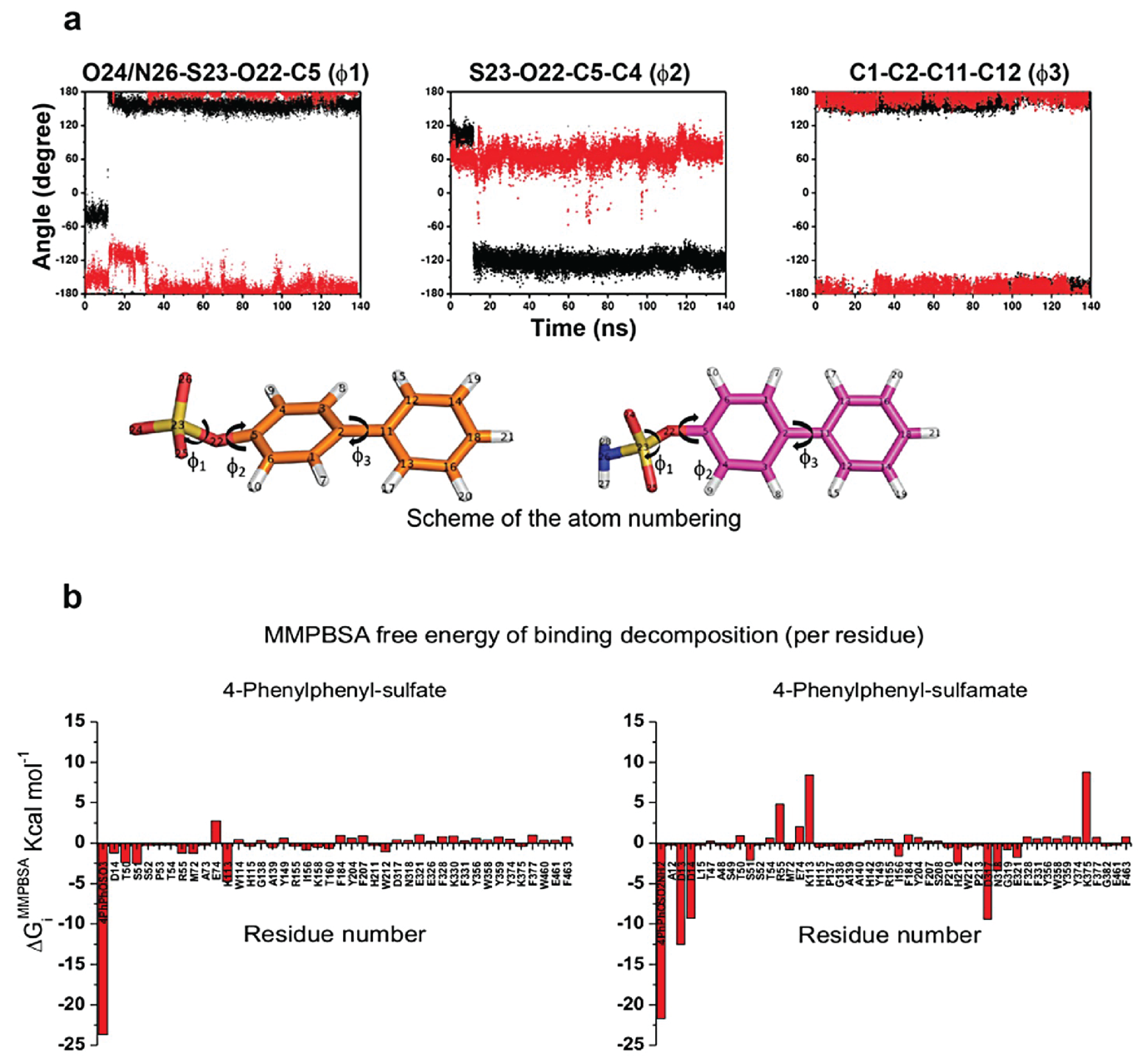

**Figure S2.** **Torsion angles and** **MMPBSA free energy plot per residue for 4-phenylphenyl sulfate and 4-phenylphenyl sulfamate bound to PaAtsA.**

**a.** Time-dependent variation of the dihedral angles φ_1_, φ_2_, φ_3_, shown in the left, middle and right plots. (Top) Dihedral angles φ_1_, φ_2_, φ_3_ that characterise the conformation of PhPh-OSO_3_ (left), and PhPh-SO_2_NH_2_ (right) are defined. Sulfur, oxygen, nitrogen, and hydrogen atoms are represented by yellow, orange/magenta, red, blue, and white. Carbon atoms in PhPh-SO_3_ and PhPh-OSO_2_NH_2_ are represented in orange and magenta, respectively. **b.** Histogram of the Poisson Boltzmann free energy of binding per residue decomposition ΔG_i_^MMPBSA^ (kcal mol^-1^) when PhPh-OSO_3_ (left) and PhPh-OSO_2_NH_2_ (right) are in bound state with PaAtsA. Contributions greater than 0.1 kcal mol^-1^ are reported.

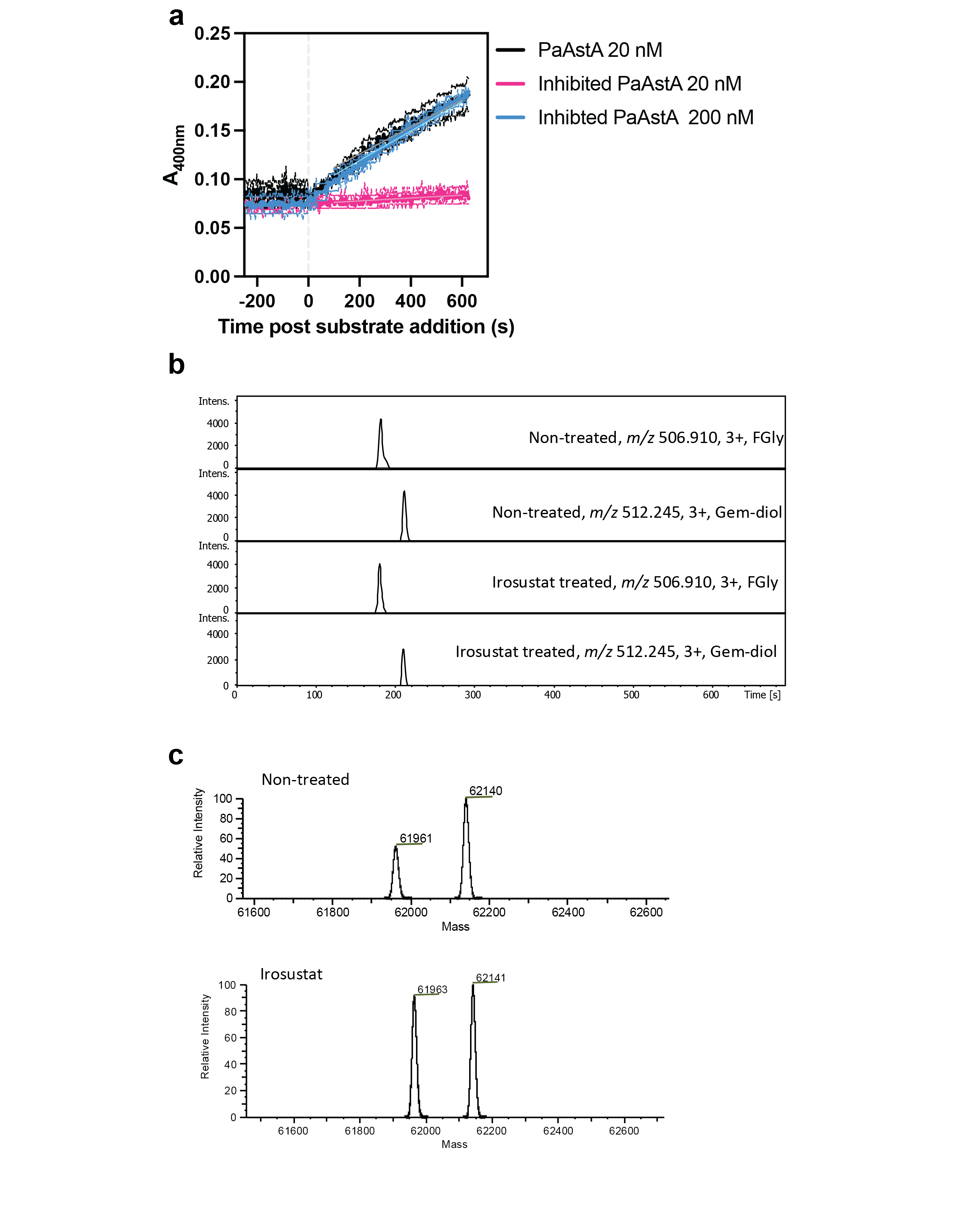

**Figure S3. Activity and molecular mass of PaAtsA before and after treatment with Irosustat.**

**a.** Activity of PaAtsA against 1 mM 4-nitrophenyl sulfate with and without overnight incubation with 1 mM Irosustat. Buffer conditions were 10 mM HEPES pH 7.5 with 150 mM NaCl. Data are technical triplicates. **b.** LC-MS extracted ion chromatograms for 3+ charge states of formylglycine (*m/z* 506.910) and gem-diol (*m/z* 512.241) modified forms of trypsin-derived PaAtsA peptide LTDFHTASTGSPTR. **c.** Deconvoluted intact mass spectra obtained from non-treated (top) and Irosustat-treated (bottom) PaAtsA. The lower mass signal is consistent with loss of N-terminal Met and gem-diol conversion from FGly (theoretical mass = 61,962 Da). The higher mass signal is consistent with the same species plus a single O-gluconoylation within the protein sequence (theoretical mass = 62,140 Da).

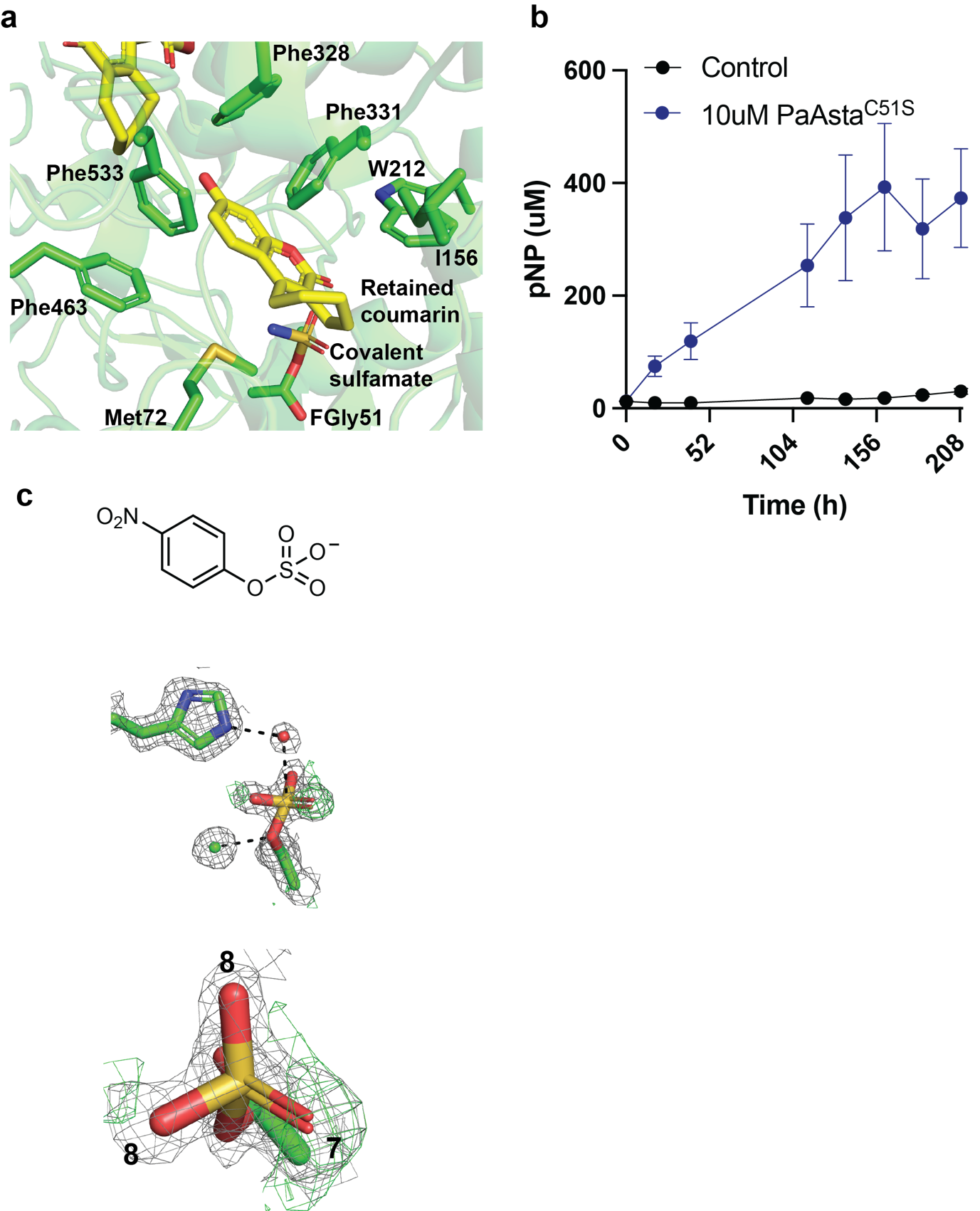

**Figure S4.** **Interaction of** **PaAtsA with the coumarin leaving group of Irosustat and the** **activity of PaAtsA^C51S^ against 4-nitrophenyl sulfate and its covalent sulfate adduct.**

**a.** The interaction of the coumarin scaffold leaving group of Irosustat in the PaAtsA binding site. Protein is in green and the Irosustat coumarin scaffold in yellow. **b.** Activity of PaAtsA^C51S^ against 10 mM 4-nitrophenyl sulfate. It is calculated that each molecule of enzyme has turned over 40 molecules of 4-nitrophenyl sulfate over the time course. Reactions were carried out in triplicate in 10 mM HEPES pH 7.5 with 150 mM NaCl. **c.** Crystal structure of PaAtsA^C51S^ covalently bound in complex with sulfate from prolonged incubation with 4-nitrophenyl sulfate. The electron density map is the 2mFobs-DFc map contoured to 1 σ. Numbers are the B factors for the relevant oxygen atoms.

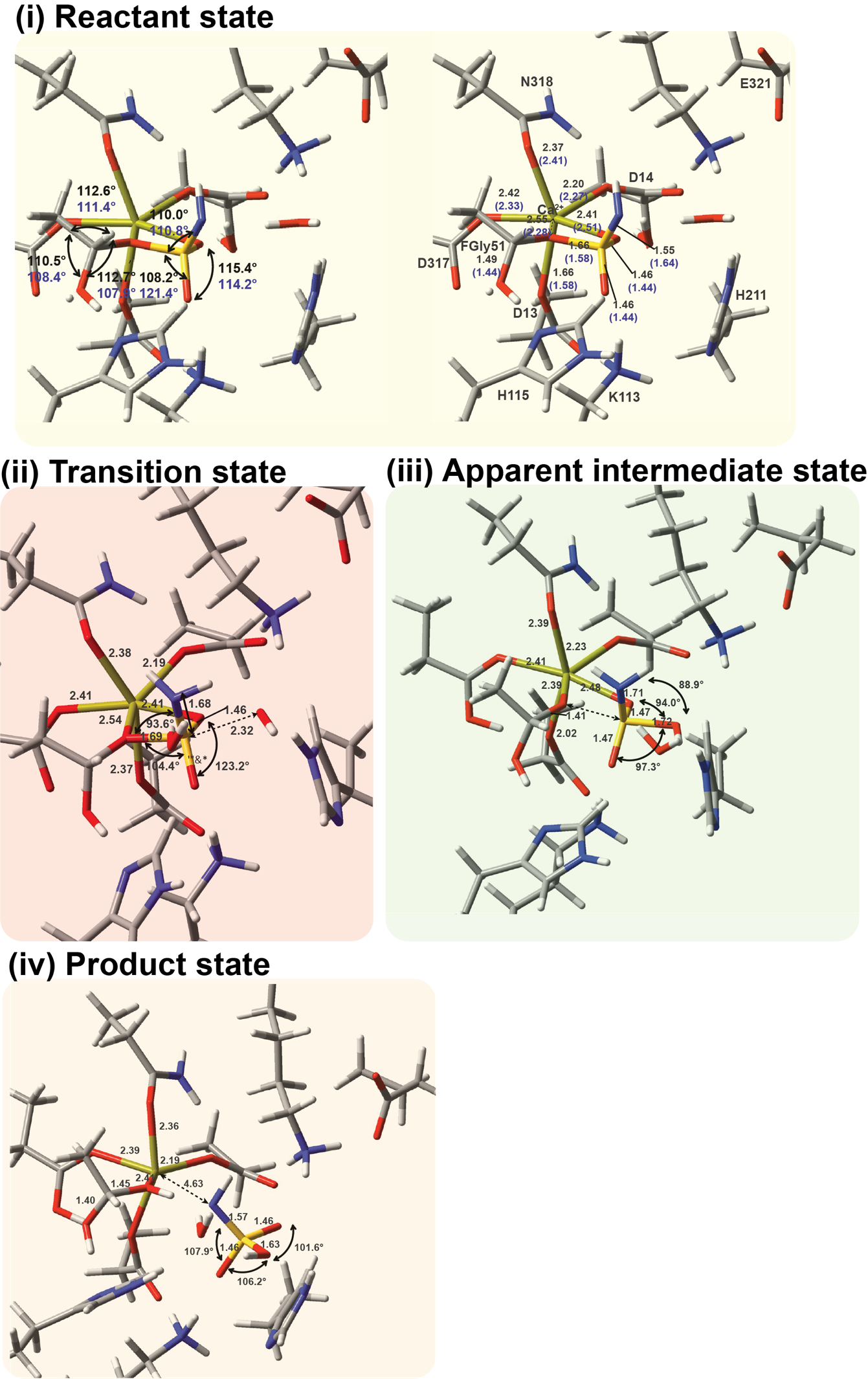

**Figure S5.** **Geometries, energies and key vibrational states along the proposed substitution pathway for breakdown of the covalent adduct.**

Carbon, oxygen, nitrogen, sulfur, calcium and hydrogen atoms are shown as grey, red, blue, yellow, gold, and white sticks, respectively. Structures represent the **(i)** reactant, **(ii)** transition state, **(iii)** apparent intermediate and **(iv)** product of the proposed substitution hydrolysis pathway. Straight and curved arrows denote selected interatomic distances and bond angles, respectively. Distances are given in Å and angles in degrees. For the reactant state, values derived from the crystallographic and DFT models are shown in blue and black, respectively.

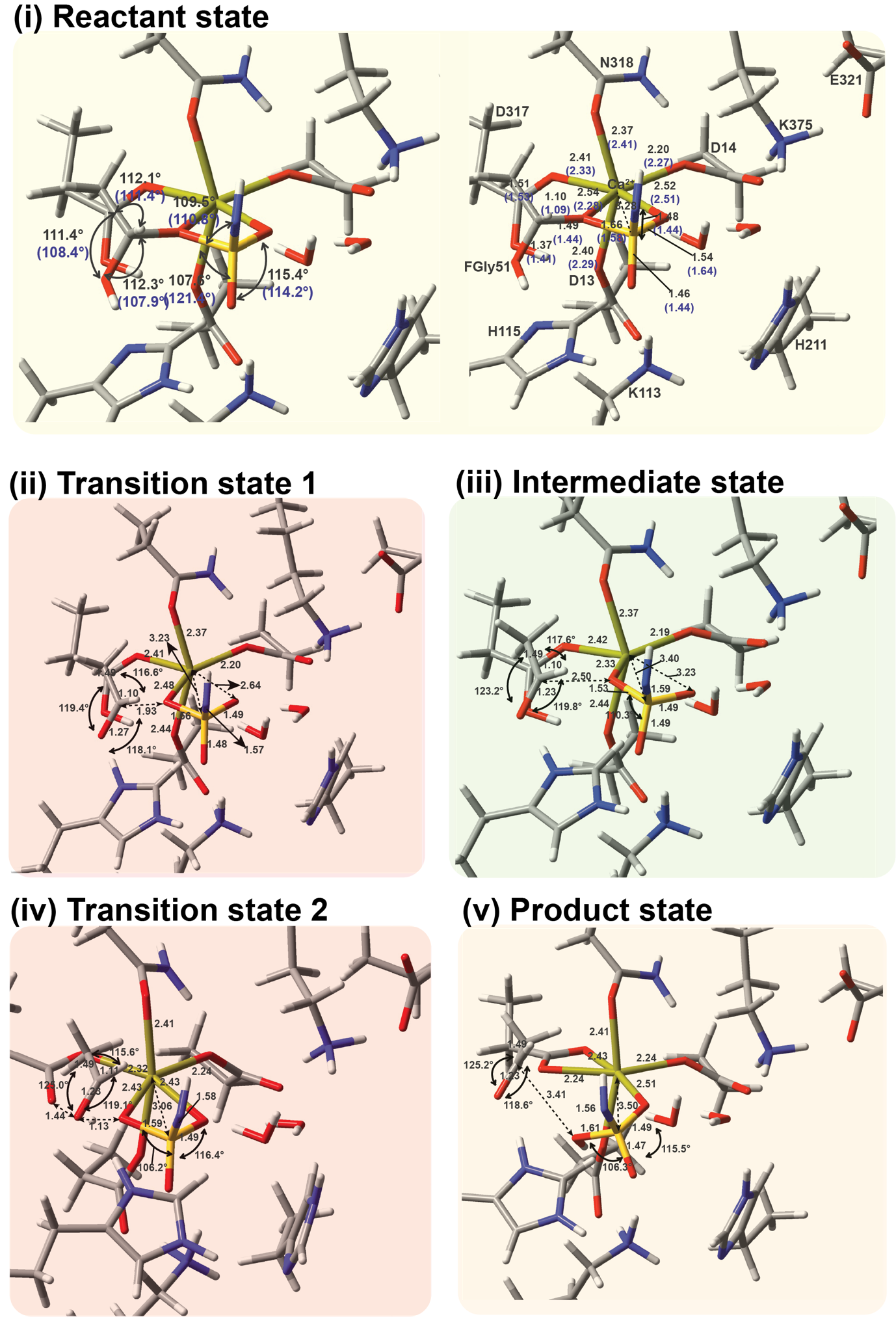

**Figure S6.** **Geometries, energies and key vibrational states along the proposed elimination pathway for breakdown of the covalent adduct.**

Carbon, oxygen, nitrogen, sulfur, and calcium, and hydrogen atoms are shown as grey, red, blue, yellow, gold, and white sticks, respectively. Structures represent the **(i)** reactant, **(ii)** first transition state (TSE1), **(iii)** intermediate, **(iv)** second transition state (TSE2) and **(v)** product. Straight and curved arrows denote selected interatomic distances and bond angles, respectively. Distances are given in Å and angles in degrees. Values derived from the crystallographic and DFT models are shown in blue and black, respectively. Selected distances within the FGly–OSO_2_NH^2^ moiety are shown in bold in (iii) and (iv).

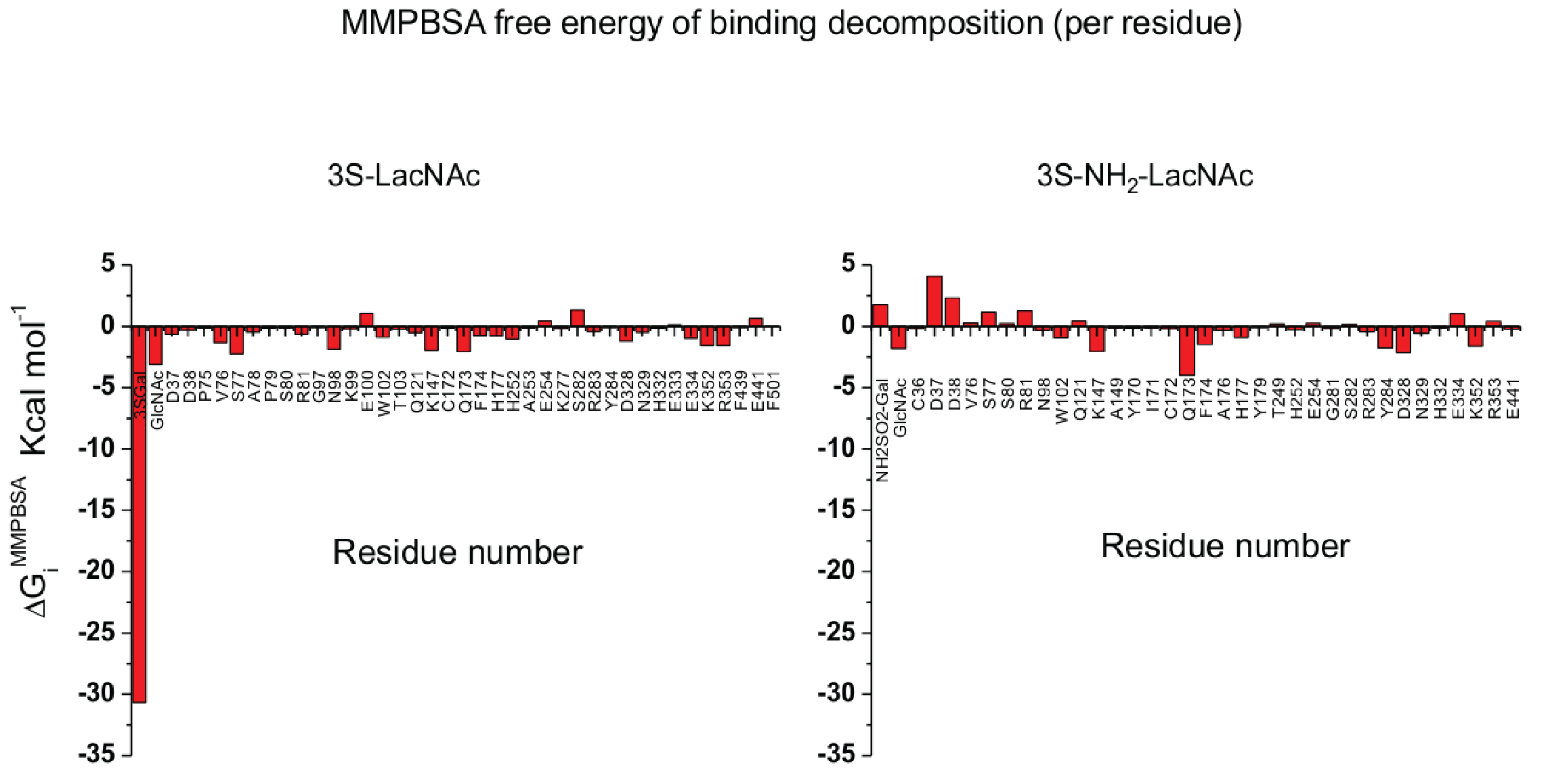

**Figure S7.** **MMPBSA Free energy plot per residue for 3S-LacNAc and 3SNH_2_-LacNAc bound to BT1636^3S-Gal^.**

Histogram of the Poisson Boltzmann free energy of binding per residue decomposition ΔG_i_^MMPBSA^ (kcal mol^-1^) when 3S-LacNAc. (left) and 3SNH_2_-LacNAc (right) are in bound state with BT1636^3S-Gal^. Contributions greater than 0.1 kcal mol^-1^ are reported.

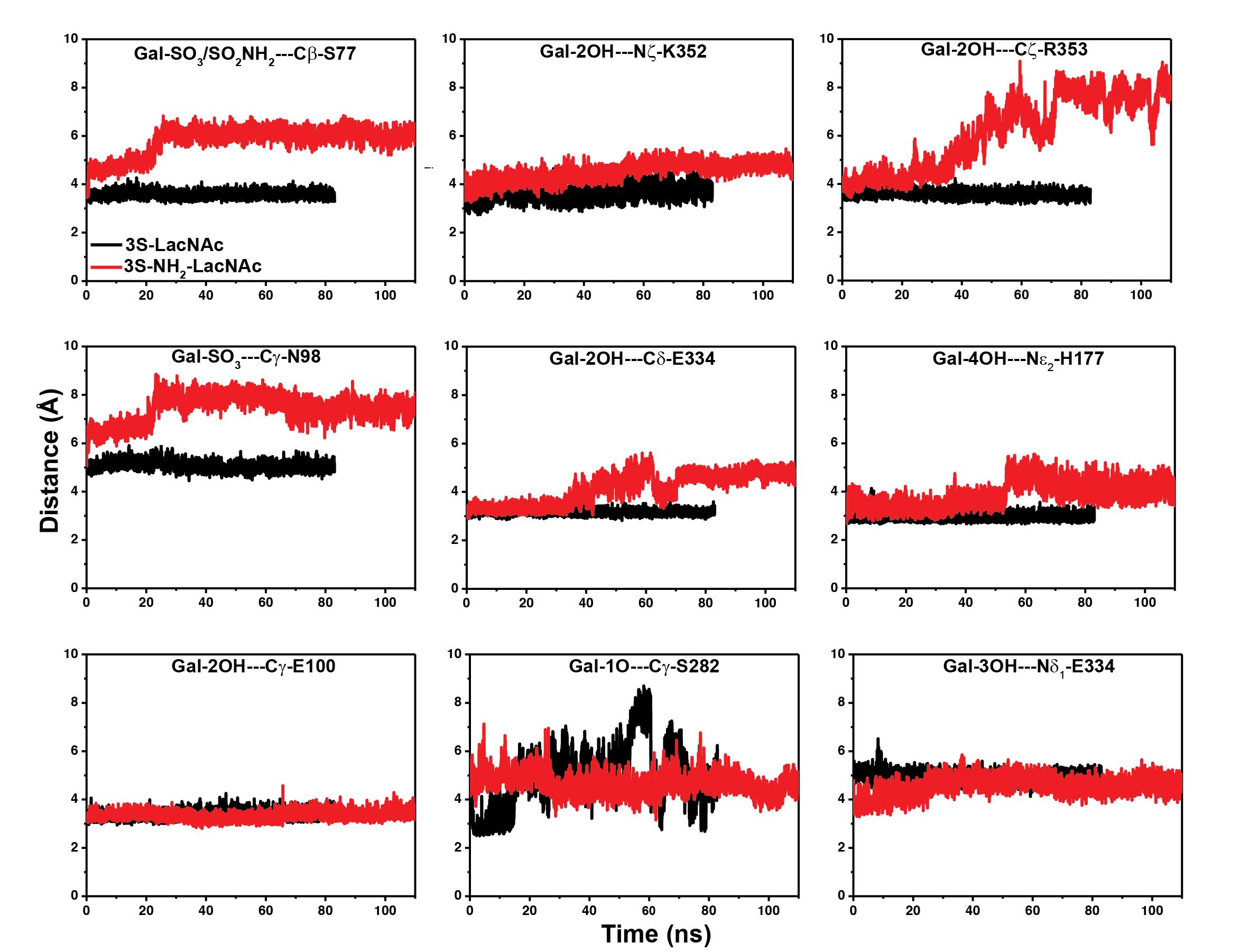

**Figure S8. Distances of key interactions in the 3S-LacNAc and 3S-NH_2_-LacNAc complexes with BT1636^3S-Gal^.**

Distances between selected enzyme and ligand atoms were monitored during the simulations shown in **Figure 5b**. Black traces represent 3S-LacNAc and red traces represent 3S-NH_2_-LacNAc.

**Table S1 Electronic DFT B3LYP energy (E^B3LYP^), zero-point energy (E^ZP^), total internal energy (E^B3LYP + ZP^), and the fundamental vibrational state (ν_0_) are reported for the optimized geometries of the O-linked intermediate of PaAsta according in its reactant, transition state, apparent intermediate, and product states, respectively, according to the SN2 hydrolysis mechanism. Energies are in Hartree, the vibrational states are in cm^-1^**

| **Energy (hartree)** | **Reactant** | **Transition state** | **Apparent intermediate** | **Product** |
| --- | --- | --- | --- | --- |
| E^B3LYP^ | -4141.2810 | -4141.2233 | -4141.2356 | -4141.2770 |
| E^ZP^ | 1.2311 | 1.2308 | 1.2319 | 1.2328 |
| E^B3LYP + ZP^ | -4140.0499 | -4139.9925 | -4140.0037 | -4140.0442 |
| ν_0_(cm^-1^) | 14.3068 | 238.0485i | 20.4547 | 19.2548 |

**Table S2 Electronic DFT B3LYP energy (E^B3LYP^), zero-point energy (E^ZP^), total internal energy (E^B3LYP + ZP^), and the fundamental vibrational state (ν_0_) are reported for the optimized geometries of the O-linked intermediate of PaAsta according in its reactant, transition state (TSE1), intermediate, transition state (TSE2) and product states, respectively, according to the elimination mechanism. Energies are in Hartree, the vibrational states are in cm^-1^**

| **Energy (Hartree)** | **Reactant** | **Transition state (TSE1)** | **intermediate** | **Transition state (TSE2)** | **Product** |
| --- | --- | --- | --- | --- | --- |
| E^B3LYP^ | -4141.2818 | -4141.2662 | -4141.2718 | -4141.2212 | -4141.2609 |
| E^ZP^ | 1.2330 | 1.2285 | 1.2318 | 1.2269 | 1.2309 |
| E^B3LYP + ZP^ | -4140.0488 | -4140.0377 | -4140.0400 | -4139.9943 | -4140.0300 |
| ν_0_(cm^-1^) | 20.9360 | -726.0547i | 16.9023 | 979.2141i | 21.4987 |

**Table S3**. **Selected ligand-receptor contact distances in the highest ranked poses generated when PhPh-O-SO_3_ and PhPh-O-SO_2_-NH_2_ are docked on PaAtsA. The reported distances are measured between the atoms underlined in bold. The atom numbers specified refer to the numbering of Scheme in Figure S2.**

| PhPh-O-SO_3_ – PaAtsA | | PhPh-O-SO_2_-NH_2_ – PaAtsA | |
| --- | --- | --- | --- |
| Contact | Distance (Å) | Contact | Distance (Å) |
| Fgly51-**Cβ**---O_3_**S**-O-PhPh | 3.8 | Fgly51-**Cβ**---NH_2_-O_2_**S**-O-PhPh | 3.5 |
| K113-**N**H_3_^+^--- O_3_**S**-O-PhPh | 4.3 | K113-**N**H_3_^+^--- NH_2_-O_2_**S**-O-PhPh | 4.5 |
| K375-**N**H_3_^+^--- O_3_**S**-O-PhPh | 3.7 | K375-**N**H_3_^+^--- NH_2_-O_2_**S**-O-PhPh | 3.3 |
| H115-Nε2--- O_3_**S**-O-PhPh | 4.4 | H115-Nε2--- NH_2_-O_2_**S**-O-PhPh | 6.4 |
| W212-CH2---(**C1**)O_3_S-O-PhPh | 3.5 | W212-CH2---(**C3**) NH_2_-O_2_**S**-O-PhPh | 3.5 |
| M72-Me---(**C5**)O_3_S-O-PhPh | 4.4 | M72-Me---(**C5**) NH_2_-O_2_**S**-O-PhPh | 4.6 |

| **Collection statistics** | BT1636 LacNAc | BT1636 3S-LacNAc | BT1636 3S-LewisA | BT1636 3S-LewisX |
| --- | --- | --- | --- | --- |
| Wavelength (Å) | 0.98 | 0.77 | 0.98 | 0.98 |
| Resolution (Å) | 56.74-1.40 (1.42-1.40) | 38.13-1.45 (1.47-1.45) | 46.089-1.41 (1.43-1.41) | 49.78-1.42 (1.44-1.42) |
| Space group | P2_1_2_1_2_1_ | P2_1_2_1_2_1_ | P2_1_2_1_2_1_ | P2_1_2_1_2_1_ |
| Unit-cell parameters |  |  |  |  |
| a, b, c (Å) | 74.37, 87.77, 103.12 | 74.390, 87.600, 103.030 | 75.06, 77.06, 89.40 | 74.50, 87.77, 103.41 |
| α, β, γ (°) | 90,90,90 | 90,90,90 | 90,90,90 | 90,90,90 |
| No. of measured reflections | 895772 (44298) | 1654713 (78239) | 675849 (33491) | 866542 (42775) |
| No. of independent reflections | 133132 (6538) | 119693 (5879) | 100356 (4883) | 128193 (6248) |
| Completeness (%) | 100 (100) | 100 (100) | 99.9 (99.7) | 100 (100) |
| Redundancy | 6.7 (6.8) | 13.8 (13.3) | 6.7 (6.9) | 6.8 (6.8) |
| <I>/<σ(I)> | 15.4 (1.4) | 15.4 (0.9) | 13.5 (1.5) | 14.9 (1.4) |
| CC(1/2) | 0.999 (0.658) | 1.00 (0.58) | 0.999 (0.773) | 0.999 (0.557) |
| **Refinement statistics*** |  |  |  |  |
| R_work_/R_free_ (%) | 13/16 | 14/17 | 13/16 | 13/15 |
| No. of non-H atoms |  |  |  |  |
| No. of protein, atoms | 7855 | 7814 | 7781 | 7827 |
| No. of solvent atoms | 404 | 304 | 367 | 421 |
| No. of ligand atoms | 41 | 45 | 23 | 45 |
| r.m.s. deviation from ideal values values |  |  |  |  |
| Bond angle (°) | 1.742 | 1.863 | 1.638 | 1.787 |
| Bond length (Å) | 0.010 | 0.014 | 0.087 | 0.011 |
| Average B factor (Å^2^) |  |  |  |  |
| Protein (by chain) | 23.5 | 25.9 | 22.9 | 23.1 |
| Solvent | 35.3 | 32.6 | 32.8 | 35.3 |
| Ligand (by chain) | 30.9 | 30.6 | 17.6 | 27.4 |
| Ramachandran plot^+^,  most favoured regions (%) | 96.9 | 96.7 | 97.4 | 96.5 |
| Molprobity score | 1.27 | 1.21 | 1.03 | 1.32 |
| PDB code | 33HO/pdb_000033ho  D_1292160063 | 33HP/pdb_000033hp  D_1292160065 | 33IL/pdb_000033il  D_1292160066 | 33HR/pdb_000033hr  D_1292160067 |

**Table S4. Crystallographic statistics table** Values in parentheses are for the highest resolution shell. R_free_ was calculated using a set (5%) of randomly selected reflections that were excluded from refinement.

| **Collection statistics** | PaAsta^Ser51^ Sulfate adduct | PaAsta Ar14 adduct  *O*-linked | PaAsta Irosustat adduct *O*-linked |
| --- | --- | --- | --- |
| Wavelength (Å) | 0.84 | 0.98 | 0.98 |
| Resolution (Å) | 92.05-1.53 (1.78-1.53) | 52.41-1.50 (1.53-1.50) | 50.79-1.63 (1.63-1.60) |
| Diffraction limits & principal axes of ellipsoid fitted to diffraction cut-off surface | 2.228 0.9876, 0, -0.1569, a  1.761 0, 1,0 b  1.571 0.1569, 0, 0.9876, c | N/A | N/A |
| Space group | C2 | C2 | P3_2_2 1 |
| Unit-cell parameters |  |  |  |
| a, b, c (Å) | 184.46, 65.90 89.64 | 189.04, 67.45, 89.72 | 59.67 59.67 275.16 |
| α, β, γ (°) | 90, 93.63, 90 | 90, 94.24, 90 | 90, 90, 120 |
| No. of measured reflections | 631141 (88277) | 1239291 (179674) | 1537307 (78083) |
| No. of independent reflections | 31498 (4414) | 60844 (8748) | 76659 (3754) |
| Completeness (%)  Spherical  Ellipsoid | 54.8 (7.6)  93.0 (57.9) | 100 (99.3)  N/A | 99.9 (100) |
| Redundancy | 7.1 (7.1) | 6.9 (7.0) | 20.1 (20.8) |
| <I>/<σ(I)> | 5.7 (1.5) | 6.7 (0.5) | 10.4 (0.5) |
| CC(1/2) | 0.980 (0.534) | 0.991 (0.405) | 0.999 (0.439) |
| **Refinement statistics*** |  |  |  |
| R_work_/R_free_ (%) | 24/26 | 25/27 | 16/23 |
| No. of non-H atoms | 16,999 | 16,892 |  |
| No. of protein, atoms | 16,615 | 16,304 | 8350 |
| No. of solvent atoms | 374 | 574 | 464 |
| No. of ligand atoms | 8 | 8 | 98 |
| r.m.s. deviation from ideal values |  |  |  |
| Bond angle (°) | 1.607 | 1.615 | 1.589 |
| Bond length (Å) | 0.007 | 0.007 | 0.007 |
| Average B factor (Å^2^) |  |  |  |
| Protein (by chain) | 13.5, 13.6 | 19.7, 19.7 | 33.4 |
| Solvent | 12.5 | 22.2 | 43.9 |
| Ligand (by chain) | 7.9, 7.5 | 18.1, 17.7 | 30.0, 46.2 |
| Ramachandran plot^+^,  most favoured regions (%) | 96.7 | 97.7 | 97.2 |
| Molprobity score | 1.62 | 1.58 | 1.46 |
| PDB code | 33IO/pdb_000033io  D_1292160084 | 33ID/pdb_000033id  D_1292160068 | 33II/pdb_000033ii  D_1292160069 |

**Table S5. Crystallographic statistics table** Values in parentheses are for the highest resolution shell. R_free_ was calculated using a set (5%) of randomly selected reflections that were excluded from refinement.

### General procedures

NMR spectra were recorded at either 295 K (300 MHz) or 300 K (600 MHz) at either Bruker AV 300 (300 MHz, 75 MHz) or Bruker AV 600 (600 MHz, 151 MHz) spectrometers. Chemical shifts are reported in ppm (δ) referenced to residual solvent peak such as DMSO (^1^H NMR: 2.50 ppm) and CHCl_3_ (^1^H NMR: 7.26 ppm). LC/MS analysis was performed on an Agilent InfinityLab series LC/MS HPLC system with an Agilent Infinity II 1260 DAD coupled to an Agilent 6120 single quadrupole mass spectrometer (ESISQ) equipped with a Thermo Fisher Scientific Accucore C18 column, 2.1 x 30 mm, 2.6 μm. Method: ESI+, flux: 0.8 ml/min, 5-95% ACN in H_2_O + 0.1% formic acid (FA), total runtime: 2.5 min or 8 min. High resolution mass spectra were recorded on an Agilent 6530 accurate-mass Q-ToF LC/MS linked to Agilent Technologies HPLC 1260 Infinity II. The HPLC was equipped with an Agilent Poroshell 120, C18 column, 2.1 x 100 mm, 1.8 µm. Method: ESI+, flux: 0.6 ml/min, 5-99% ACN in H_2_O + 0.1% FA, total runtime: 4.5 min. Preparative HPLC was performed on a Gilson PLC 2250 with a Macherey-Nagel VP 250/21 Nucleodur 100-7 C18Ec column (30 mL/min flow). Purity and characterization of all final compounds was established by a combination of LC-MS, LC-HRMS and NMR analytical techniques. All tested compounds were found to be >95% pure by LC/MS and HRMS analysis.

#### ****Table S6. Calculated or literature known pKa values of alcohol leaving group.****

| **Compound** | **Structure** | **pKa lit** | **pKa cal*** | **PaAtsa**  **Fractional activity in assay** | **BT1636^3S-Gal^**  **Fractional activity in assay** |
| --- | --- | --- | --- | --- | --- |
| **1** | 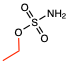 | 16 | 14.63 | 0.93 | 0.96 |
| **2** | 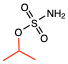 | 17.1 | 15.04 | 0.56 | 0.92 |
| **3** | 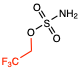 | 12.5 | 10.73 | 0.91 | 1.40 |
| **4** | 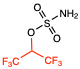 | 9.3 | 6.92 | 0.013 | 0.90 |
| **5** | 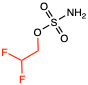 | 13 | 12.31 | 0.75 | 1.35 |
| **6** | 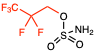 | 10.5 | 10.65 | 0.36 | 1.17 |
| **7** | 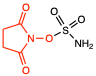 | 6 | 7.88 | 0.05 | 0.58 |
| **8** | 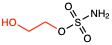 | 14.22 | 14.4 | 0.54 | 0.90 |
| **9** | 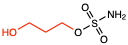 | 15.6 | 12.7 | 0.76 | 1.19 |
| **10** | 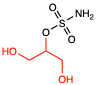 |  | 12.6 | 0.21 | 0.53 |
| **11** | 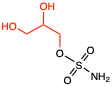 |  | 12.22 | 0.74 | 1.17 |
| **12** | 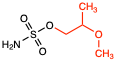 |  | 15.55 | 0.88 | 1.32 |
| **13** | 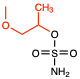 |  | 15.87 | 0.64 | 1.18 |

Ref for lit value is given as a ref

*Calculated using Rowen (cite: <https://doi.org/10.26434/chemrxiv-2024-8489b>)

#### ****General Procedure for Sulfamate Ester Synthesis A****

To a round-bottom flask equipped with a stir bar under N_2_, 2 mL of CH_3_CN and 1 mL of chlorosulfonyl isocyanate (2.20 eq.) were added, and the solution was cooled to 0 °C. Next, 0.45 mL of formic acid (2.24 eq.) was added dropwise via syringe and gas evolution was observed immediately. The resulting mixture was stirred at 0 °C for 30 min, then diluted with 16 mL of CH_3_CN and allowed to gradually warm to room temperature, stirring for an additional 2 h. The mixture was then cooled to 0 °C, and 0.5mL of alcohol in 2 mL of DMA was added dropwise. After 10 min, the reaction was warmed to room temperature and stirred for 16 h. The reaction was quenched by the addition of 20 mL of H_2_O and transferred to a separatory funnel. The aqueous layer was extracted with EtOAc (2 × 50 mL). The combined organic layers were washed successively with H_2_O (3 × 20 mL) and brine (1 × 20 mL), dried over Na_2_SO_4_, decanted, and concentrated *in vacuo*. The product was purified by silica gel flash chromatography to give the desired sulfamate ester product.

#### ****General Procedure for Sulfamate Ester Synthesis B****

To a round-bottom flask equipped with a stir bar under N_2_, 2 mL of CH3CN and 1 mL of chlorosulfonyl isocyanate (2.20 eq.) were added, and the solution was cooled to 0 °C. Next, 0.45 mL of formic acid (2.24 equiv) was added dropwise via syringe and gas evolution was observed immediately. The resulting mixture was stirred at 0 °C for 30 min, then diluted with 16 mL of CH3CN and allowed to gradually warm to room temperature, stirring for an additional 2 h. The mixture was then cooled to 0 °C, and 0.5mL of alcohol in 2 mL of DMA was added dropwise. After 10 min, the reaction was warmed to room temperature and stirred for 16 h. The reaction was quenched by the addition of 20 mL of H_2_O and transferred to a separatory funnel. The aqueous layer was extracted with EtOAc (2 × 50 mL). The combined organic layers were washed successively with H_2_O (3 × 20 mL) and brine (1 × 20 mL), dried over Na_2_SO_4_, decanted, and concentrated *in vacuo*. The product was purified by silica gel flash chromatography to give the desired sulfamate ester intermediate. The intermediate was dissolved in methanol with a drop of HCl (1M), and stirred at room temperature until cleavage of the acid labile protecting groups was confirmed by LCMS, the sample was concentrated *in vacuo* to yield the sulfamate ester product after silica gel flash chromatography.

### Compound Characterisation

#### Ethyl sulfamate

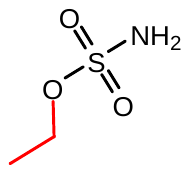

Following the general procedure A with ethanol yielded the title sulfamate ester (630 mg, 34%). **^1^H NMR** (300 MHz, CDCl_3_) δ 4.29 (q, *J* = 7.1 Hz, 2H), 1.41 (t, *J* = 7.1 Hz, 3H).**^13^C NMR** (75 MHz, CDCl_3_) δ 67.85, 14.74. **HRMS** QTOF-MS: calcd. C_2_H_7_NO_3_S for [M+Na]^+^: cald. 148.0039, found 148.0045

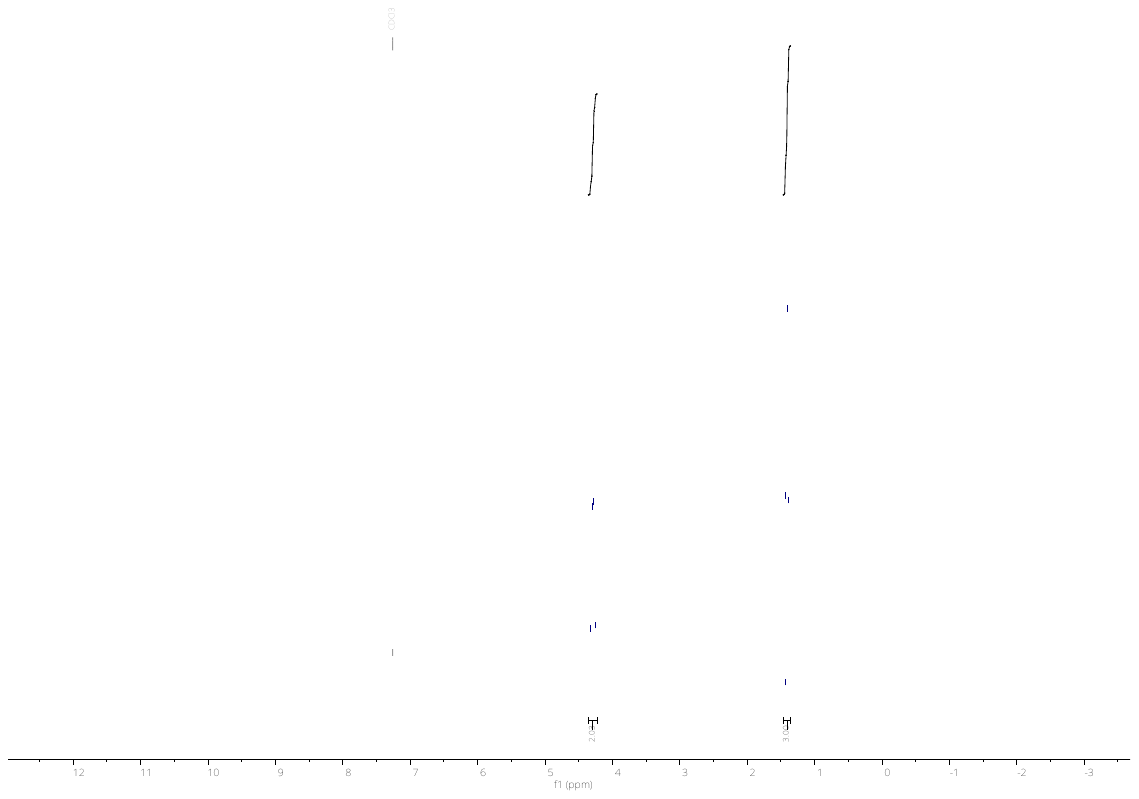
**^1^H NMR**

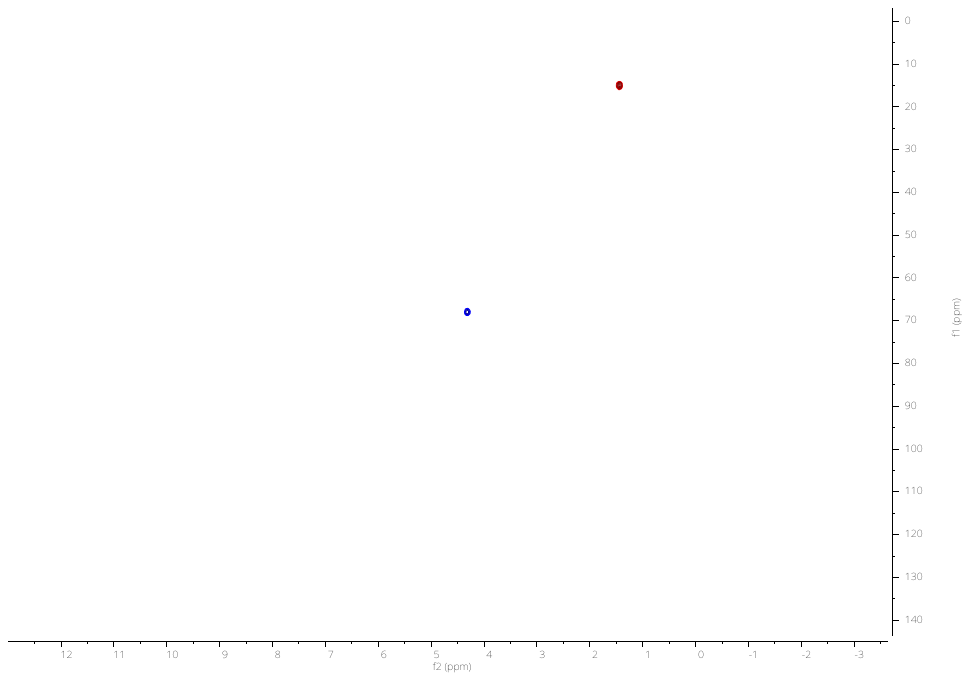

**^1^H-^13^C HSQC NMR**

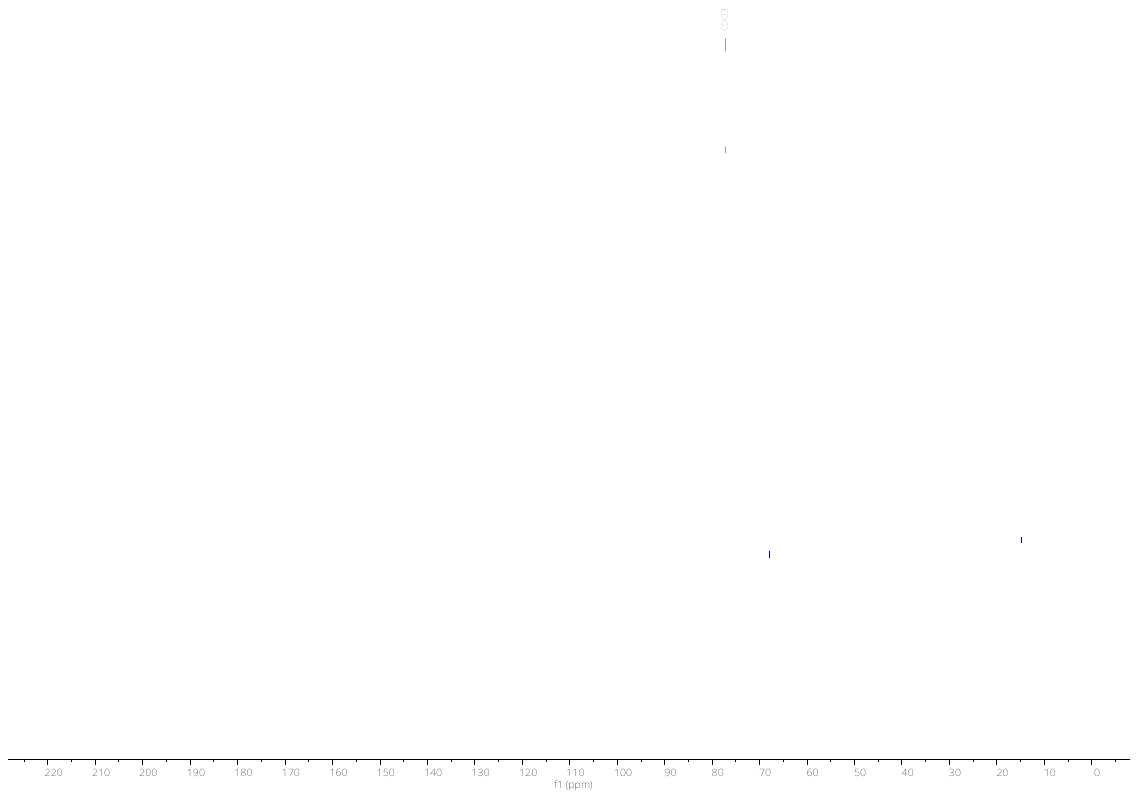

**^13^C NMR**

#### isopropyl sulfamate

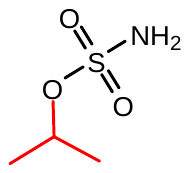

Following the general procedure A with isopropanol yielded the title sulfamate ester (250 mg, 61%). **^1^H NMR** (300 MHz, CDCl_3_) δ 4.90 – 4.72 (m, 1H), 1.44 – 1.31 (m, 6H).**^13^C NMR** (75 MHz, CDCl_3_) δ 78.16, 22.62. **HRMS** QTOF-MS: calcd. C_3_H_9_NO_3_S for [M+Na]^+^: cald. 162.0195, found 162.0192

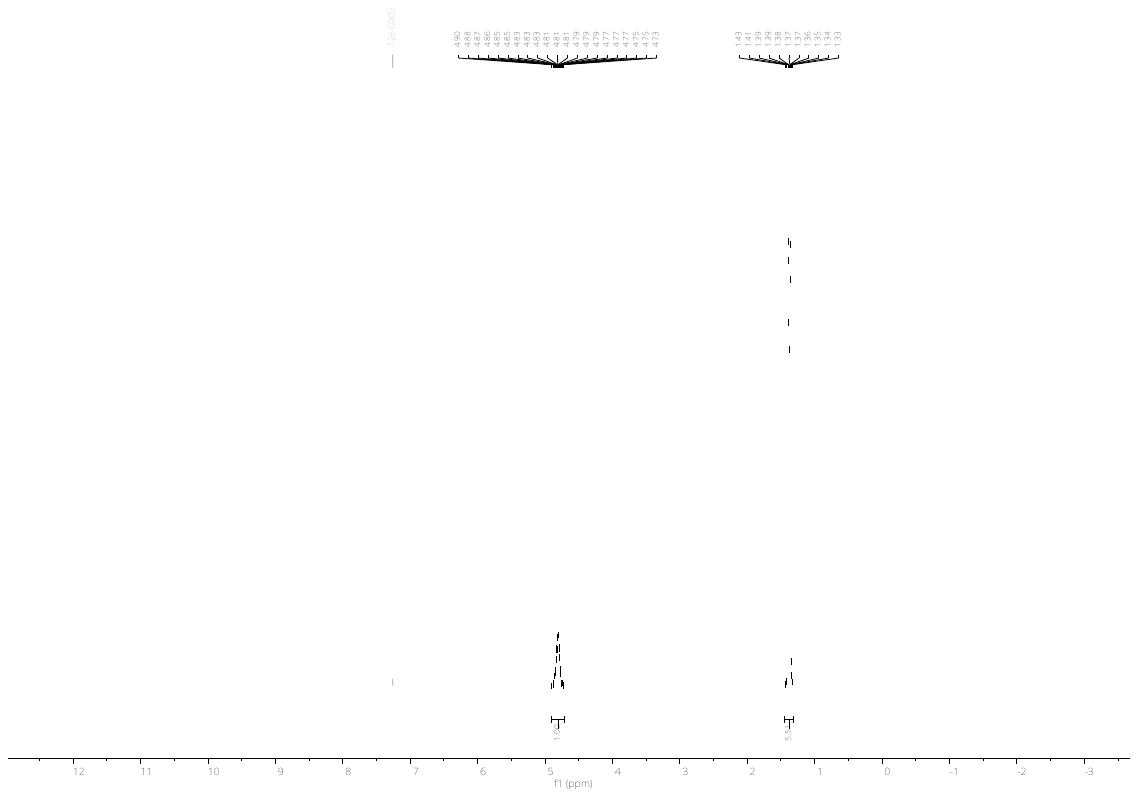

**^1^H NMR**

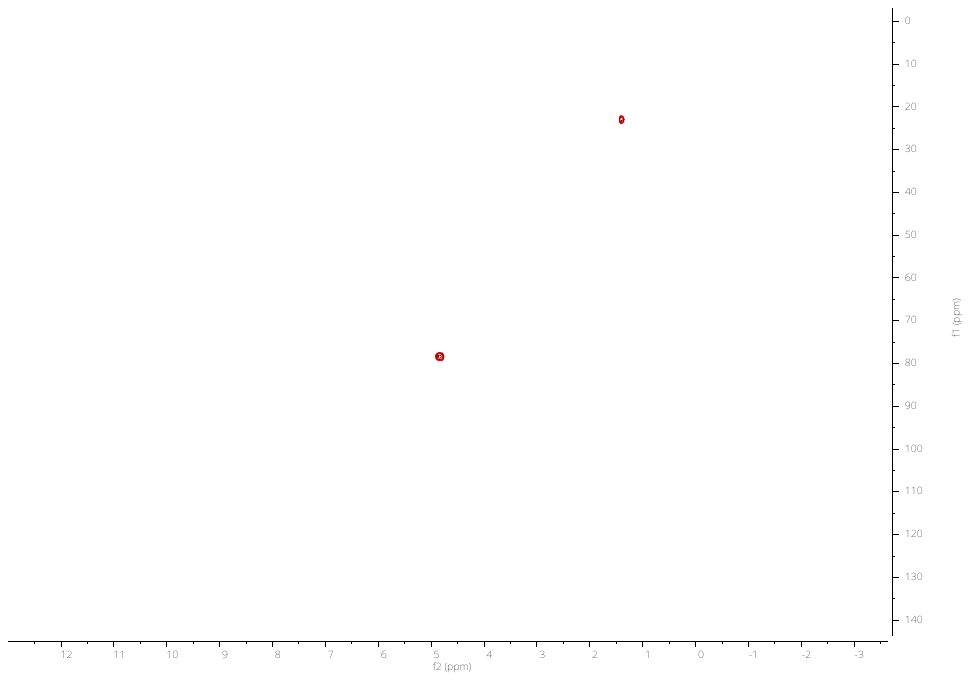

**^1^H-^13^C HSQC NMR**

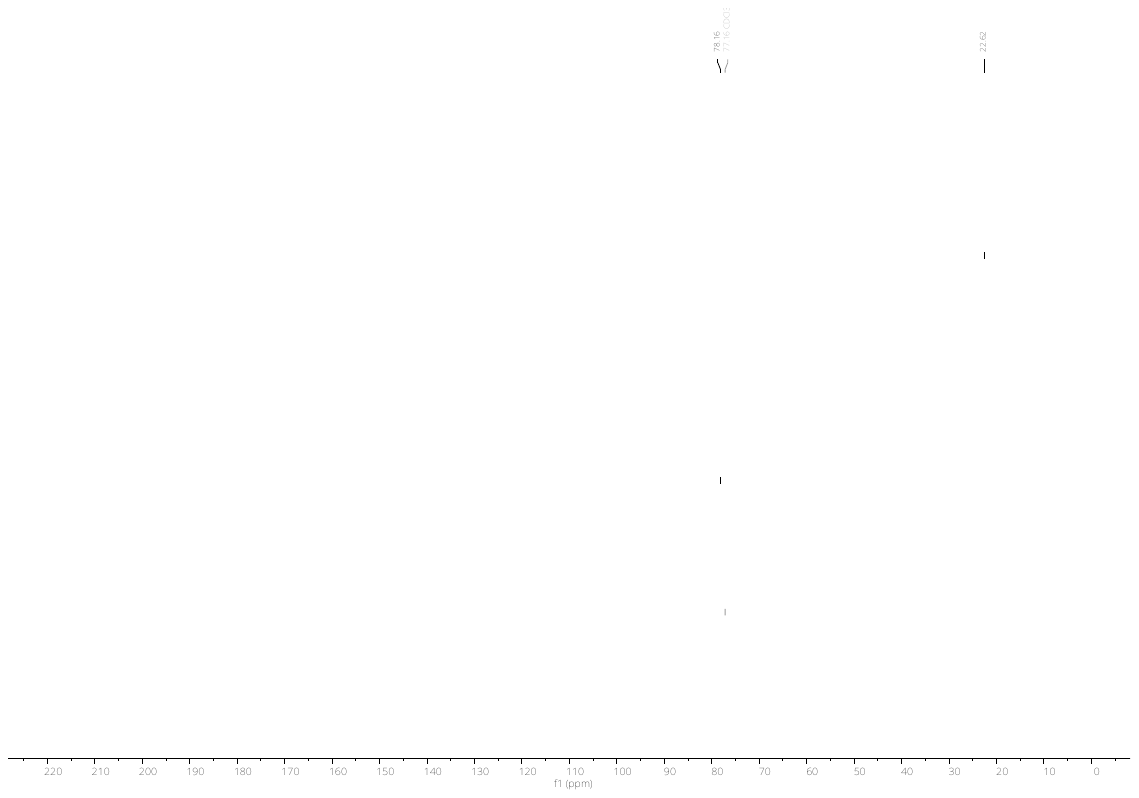

**^13^C NMR**

#### Sulfamate *N*-hydroxysuccinimide ester

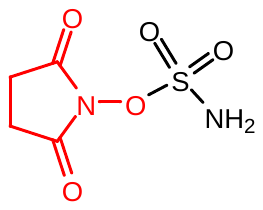

Following the general procedure A with *N*-hydroxysuccinimide yielded the title sulfamate ester (50 mg, 12%). **^1^H NMR** (600 MHz, MeOD) δ 4.83 (s, 2H), 2.68 (s, 4H). **^13^C NMR** (151 MHz, MeOD) δ 174.85, 26.28. **HRMS** QTOF-MS: calcd. C_4_H_6_N_2_O_5_S for [M-H]^-^ : cald. 192.9925, found 192.9924.

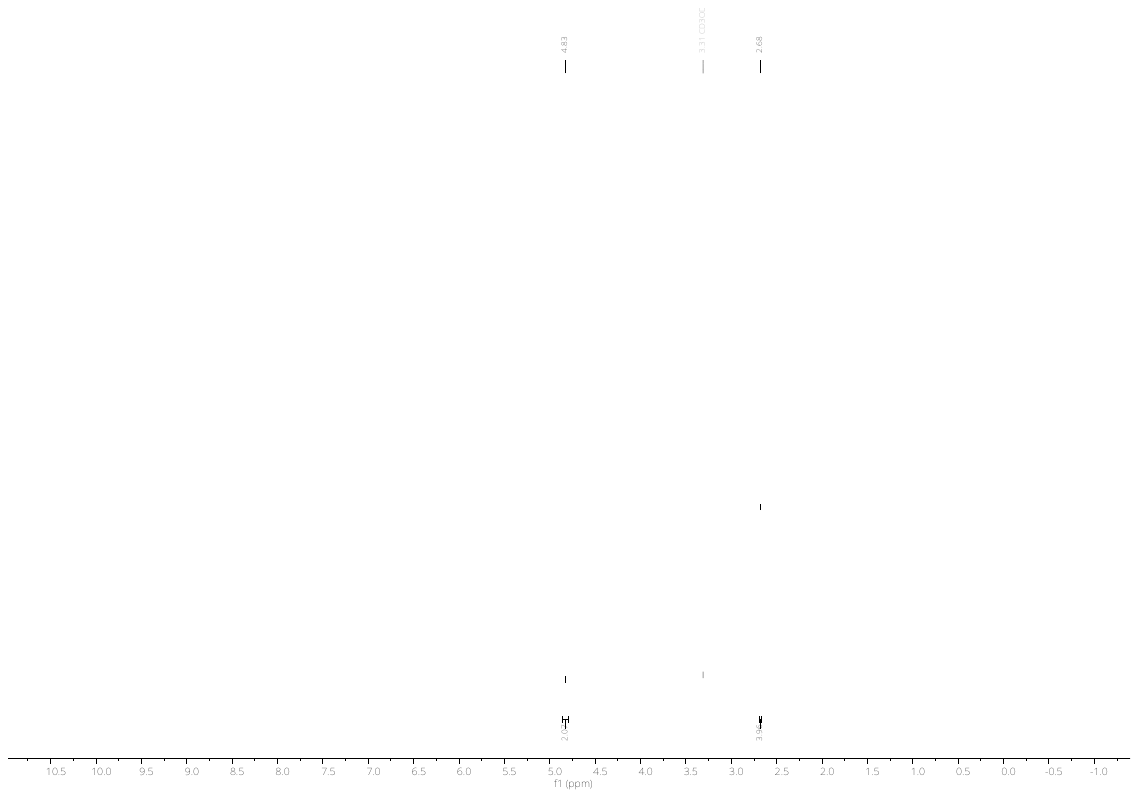

**^1^H NMR**

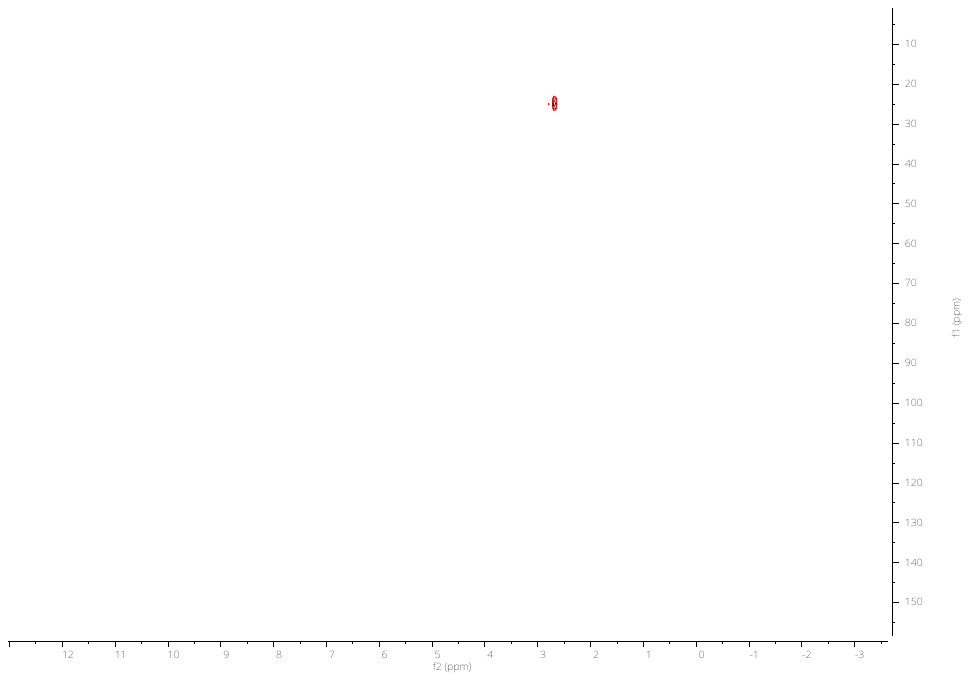

**^1^H-^13^C HSQC NMR**

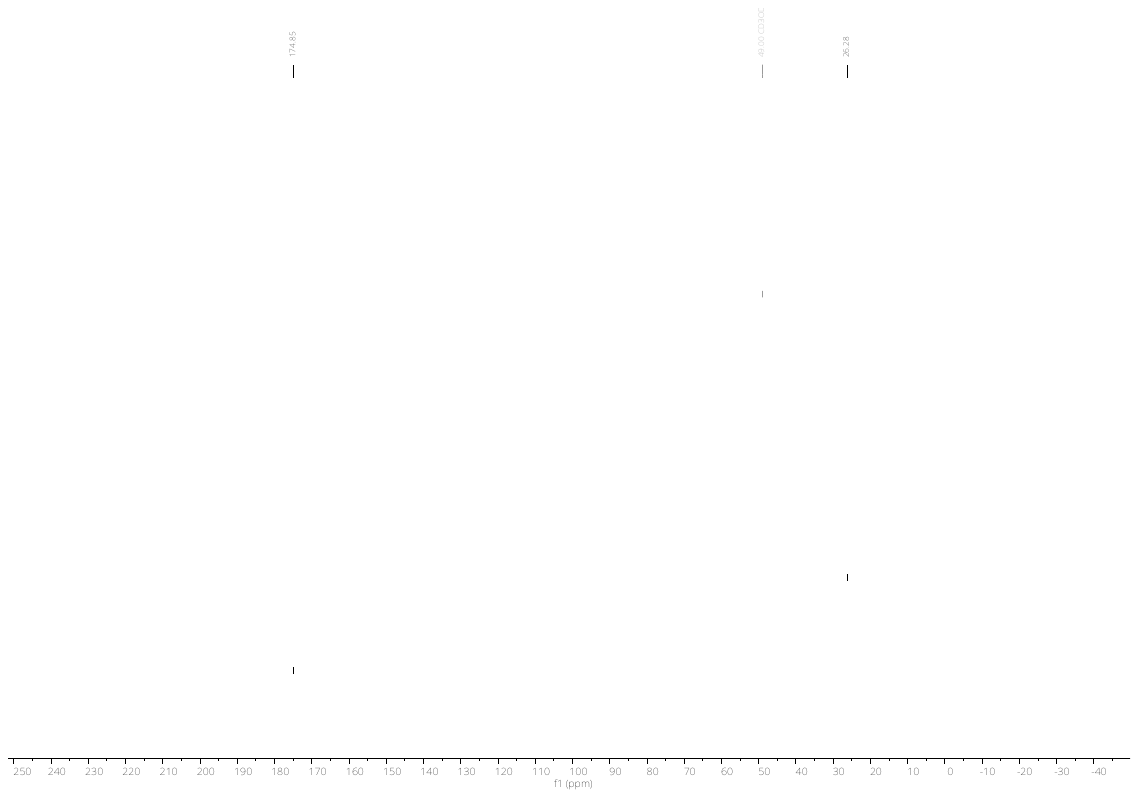

**^13^C NMR**

#### 2,2,2-trifluoroethyl sulfamate

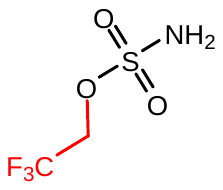

Following the general procedure A with 2,2,2-trifluoroethanol yielded the title sulfamate ester (340 mg, 53%). **^1^H NMR** (300 MHz, CDCl_3_) δ 4.53, 4.50, 4.49, 4.48, 4.45.**^13^C NMR** ^13^C NMR (75 MHz, CDCl_3_) δ 65.70 (q, *J* = 38.0 Hz).**^19^F NMR** (282 MHz, CDCl_3_) δ 3.76. **HRMS** QTOF-MS: calcd. C_2_H_4_F_3_NO_3_S for [M-H]^-^: cald. 177.9791, found 177.9800

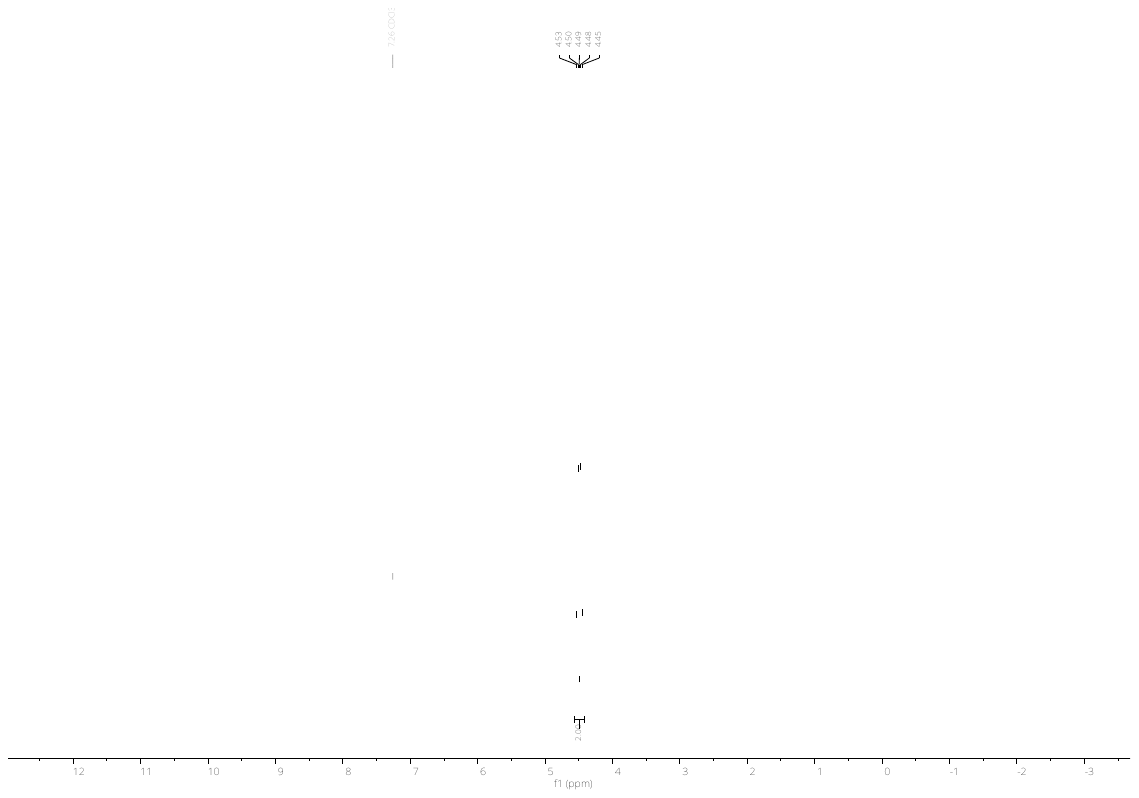

**^1^H NMR**

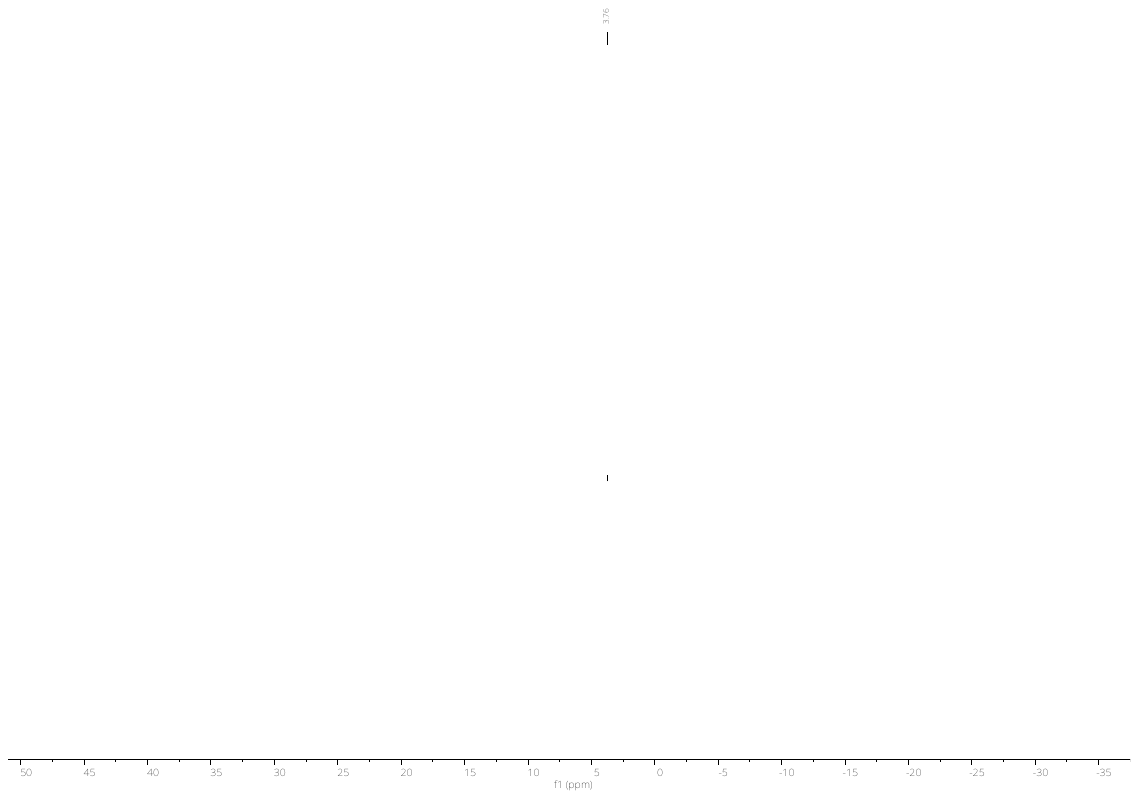

**^19^NMR**

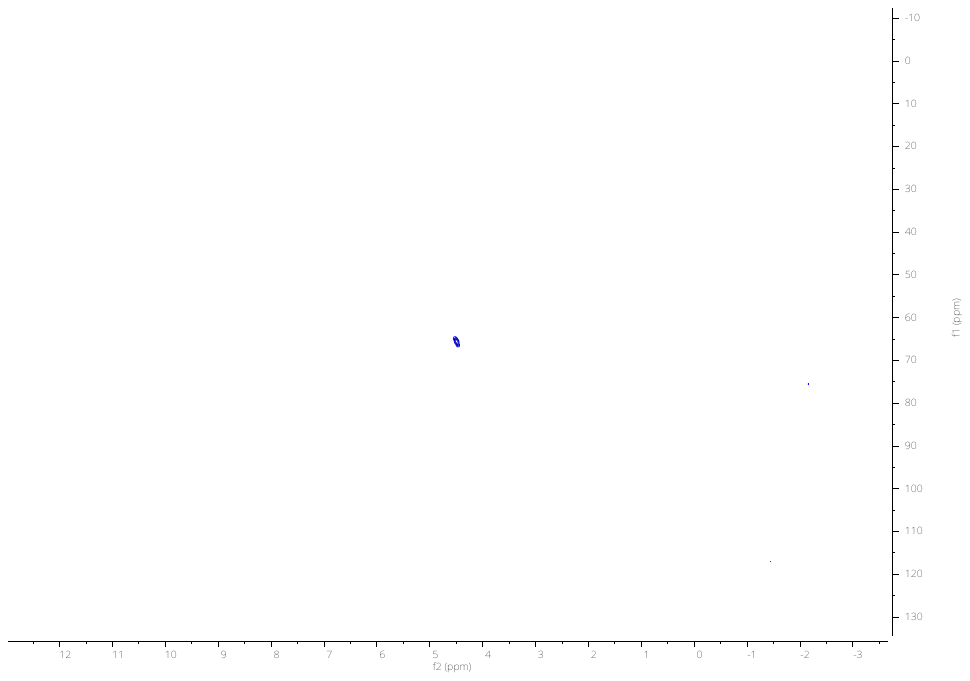

**^1^H-^13^C HSQC NMR**

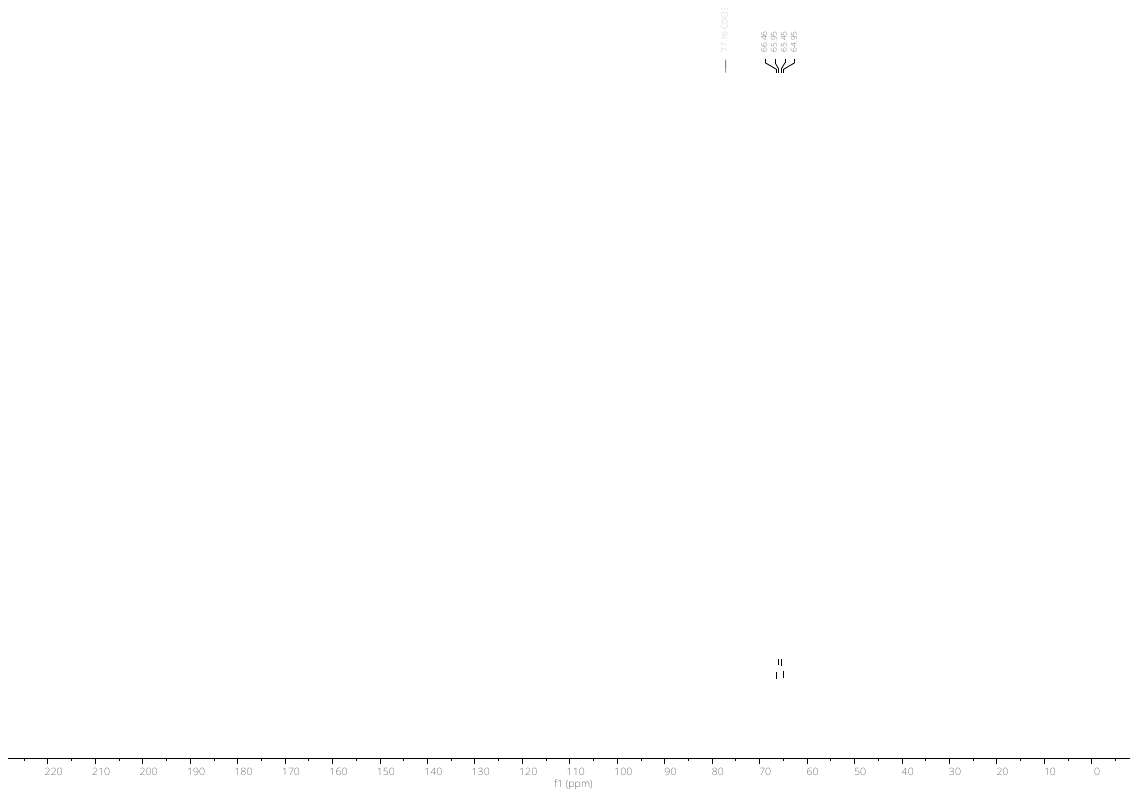

**^13^C NMR**

#### 1,1,1,3,3,3-hexafluoropropan-2-yl sulfamate

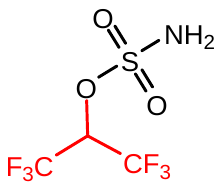

Following the general procedure A with 1,1,1,3,3,3-hexafluoro-2-propanol yielded the title sulfamate ester (450 mg, 48%). **^1^H NMR** (300 MHz, Acetone) δ 7.70 (s, 2H), 5.85 (hept, *J* = 5.9 Hz, 1H). **^13^C NMR** (75 MHz, Acetone) δ 206.27 (tt, *J* = 1.8, 1.0 Hz), 129.40 – 113.59 (m), 72.74 (dt, *J* = 68.8, 34.4 Hz). **^19^F NMR** (564 MHz, CDCl_3_) δ -73.18 (s, 1F). **HRMS** QTOF-MS: calcd. C_3_H_3_F_6_NO_3_S for [M-H]^-^: cald. 245.9665, found 245.9655.

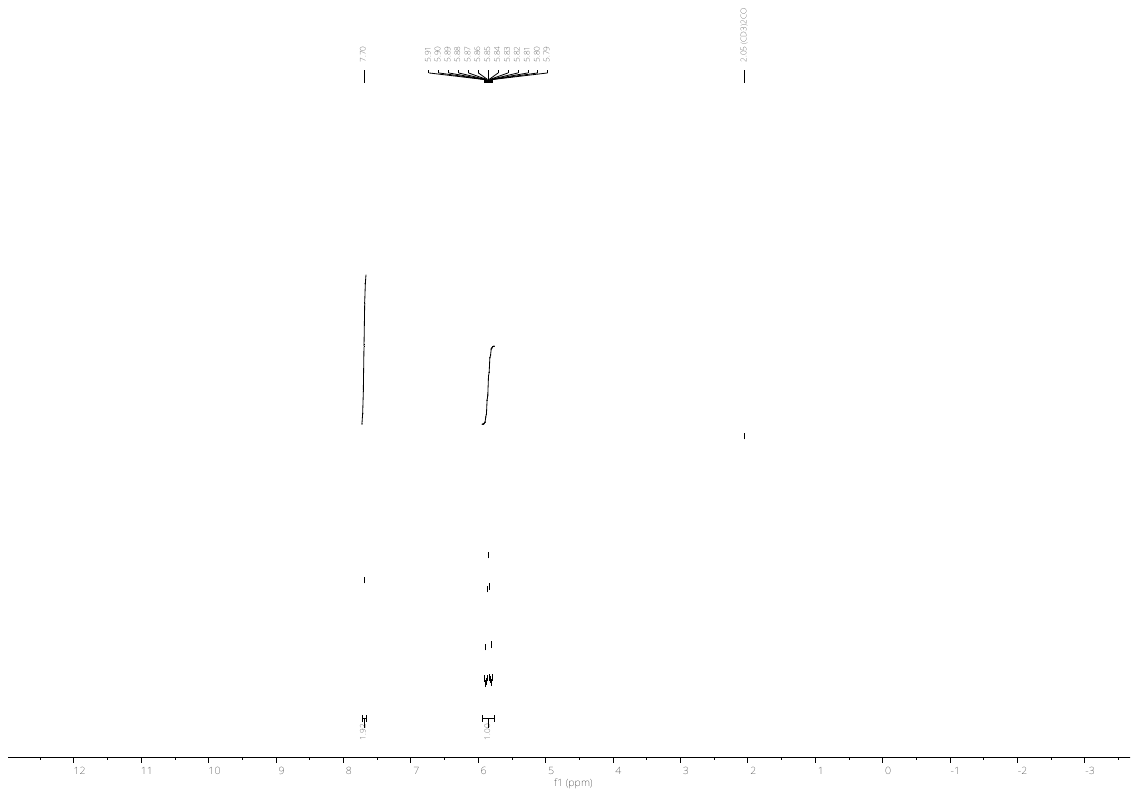

**^1^H NMR**

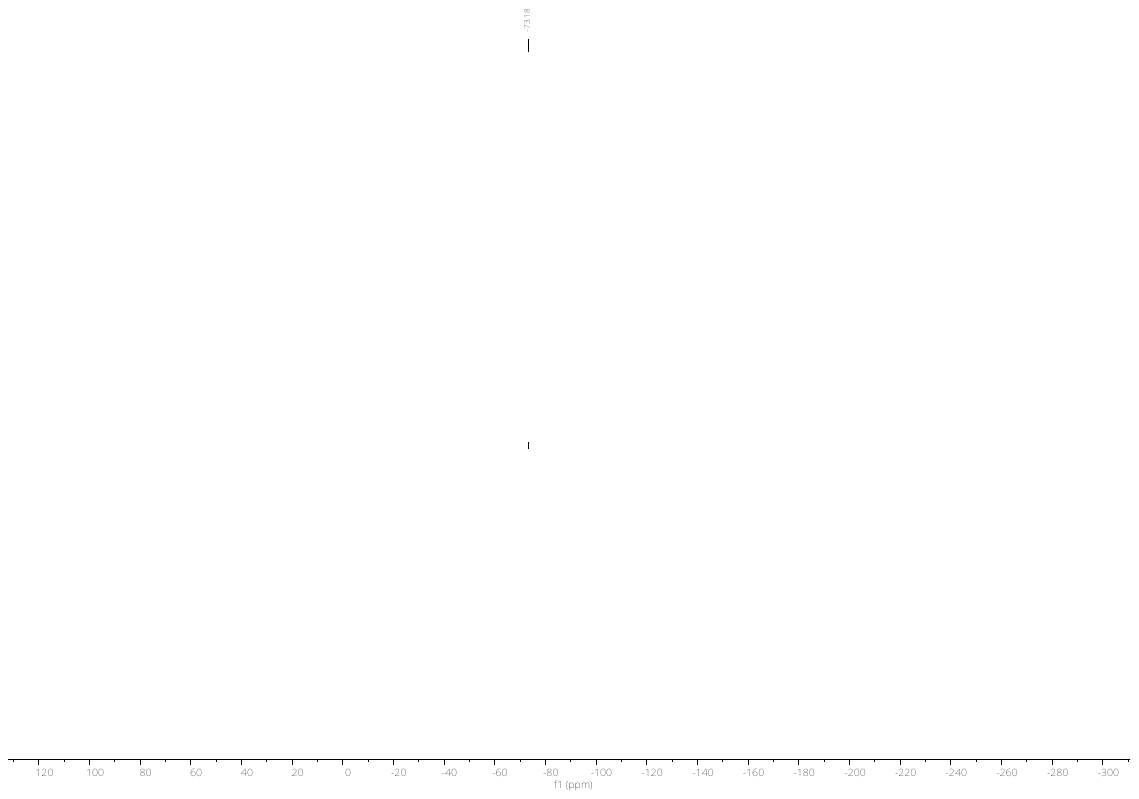

**^19^F NMR**

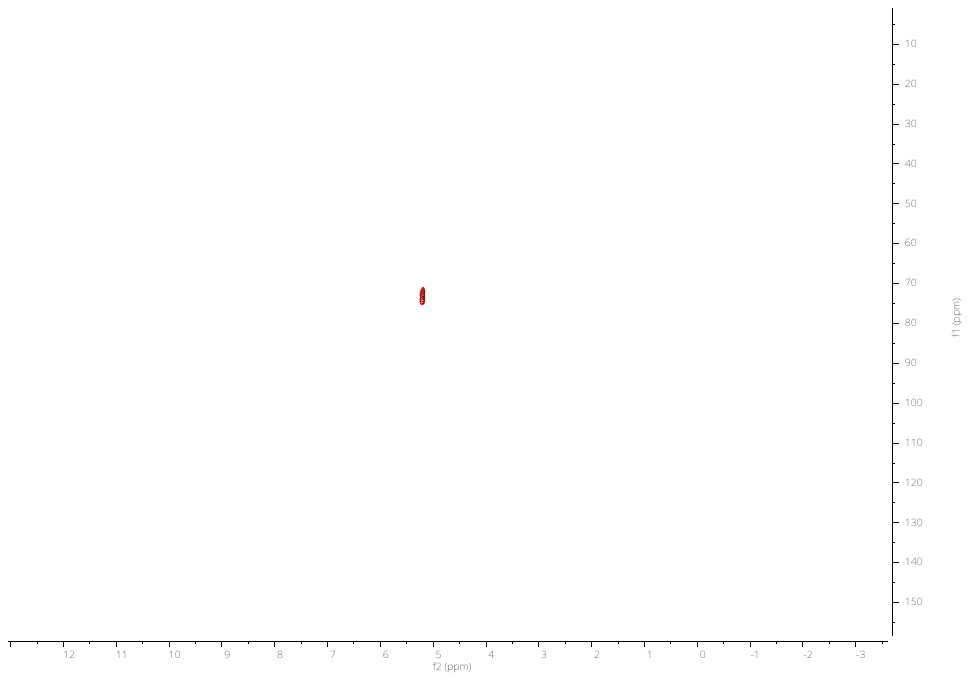

**^1^H-^13^C HSQC NMR**

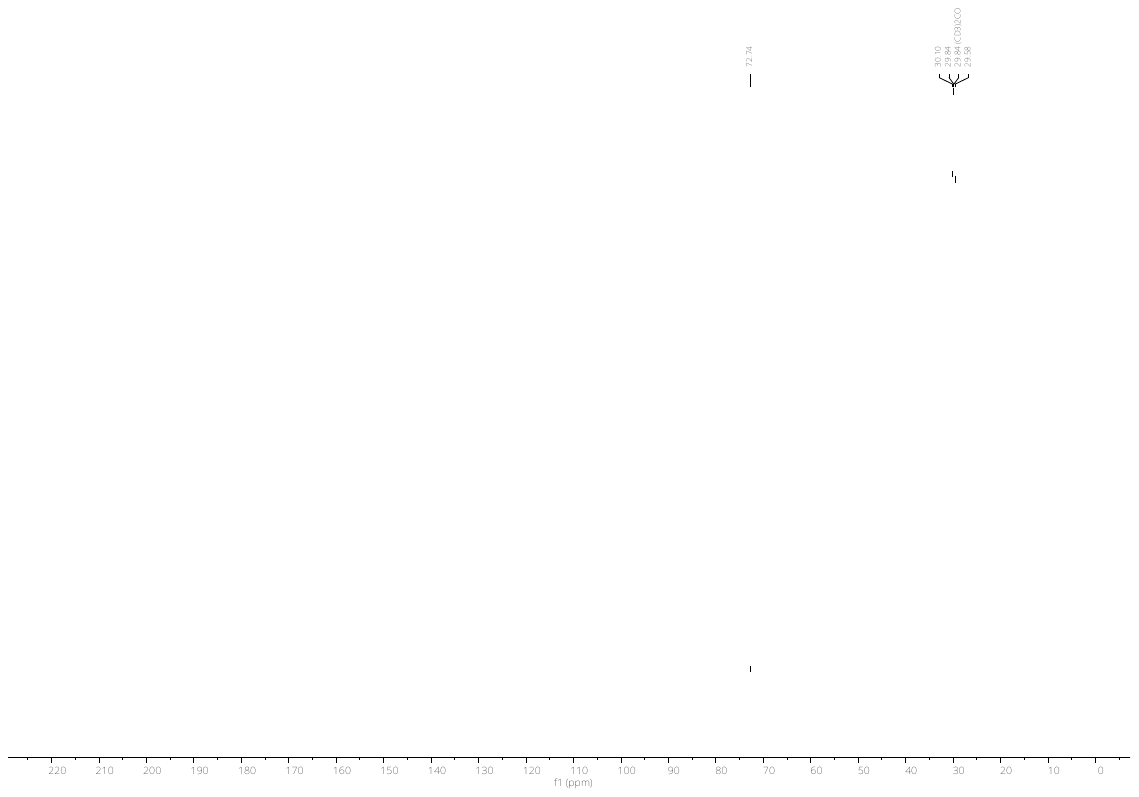

**^13^C NMR**

#### 2,2-difluoroethyl sulfamate

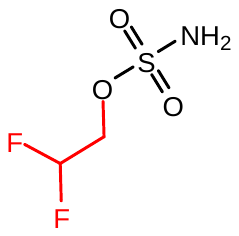

Following the general procedure A with 2,2-difluoroethanol yielded the title sulfamate ester (402 mg, 55%). **^1^H NMR** (300 MHz, MeOD) δ 6.10 (tt, *J* = 54.7, 3.8 Hz, 1H), 4.25 (td, *J* = 13.7, 3.8 Hz, 2H).**^13^C NMR** (75 MHz, MeOD) δ 117.45, 114.27, 111.09, 68.18, 67.79, 67.40.**^19^F NMR** (282 MHz, MeOD) δ -50.62. **HRMS** QTOF-MS: calcd. C_2_H_5_F_2_NO_3_S for [M-H]^-^: cald. 159.9885, found 159.9886

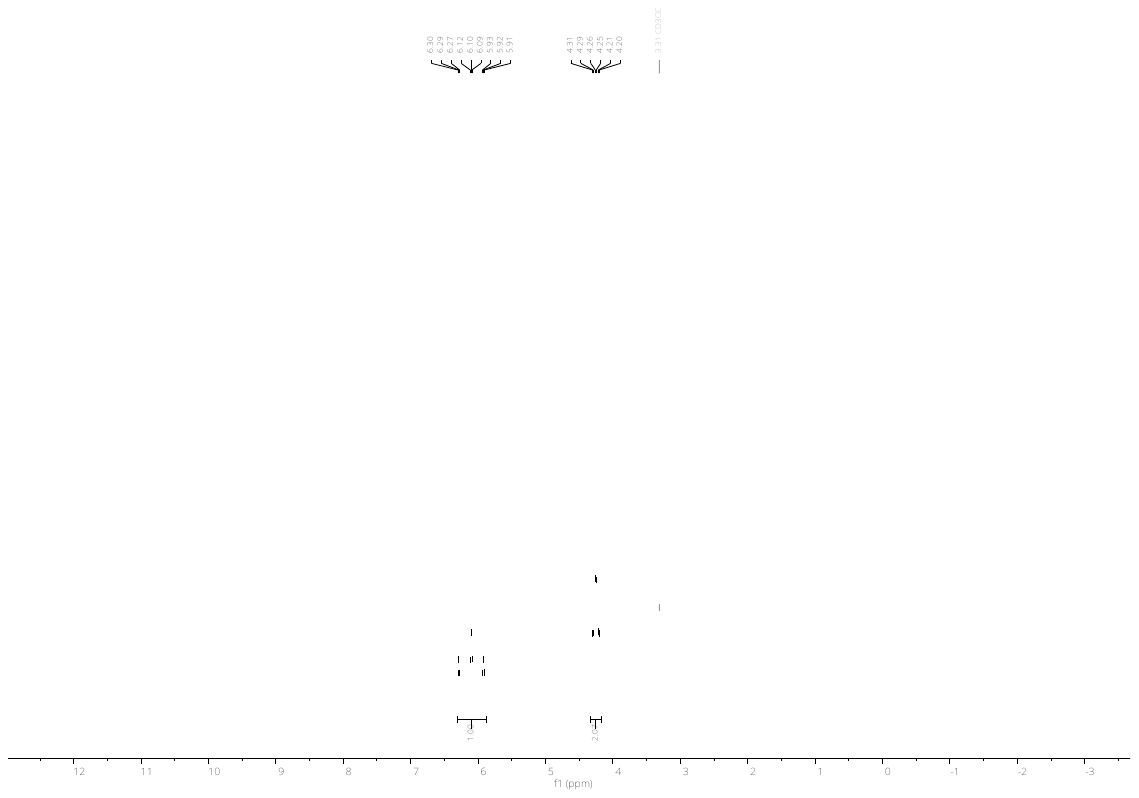

**^1^H NMR**

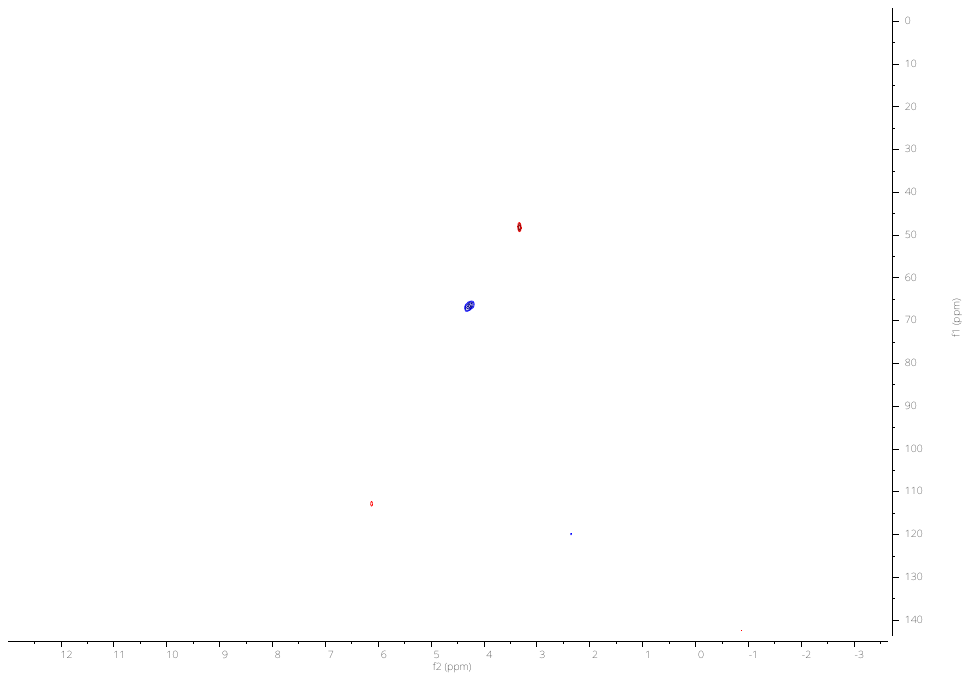

**^1^H-^13^C HSQC NMR**

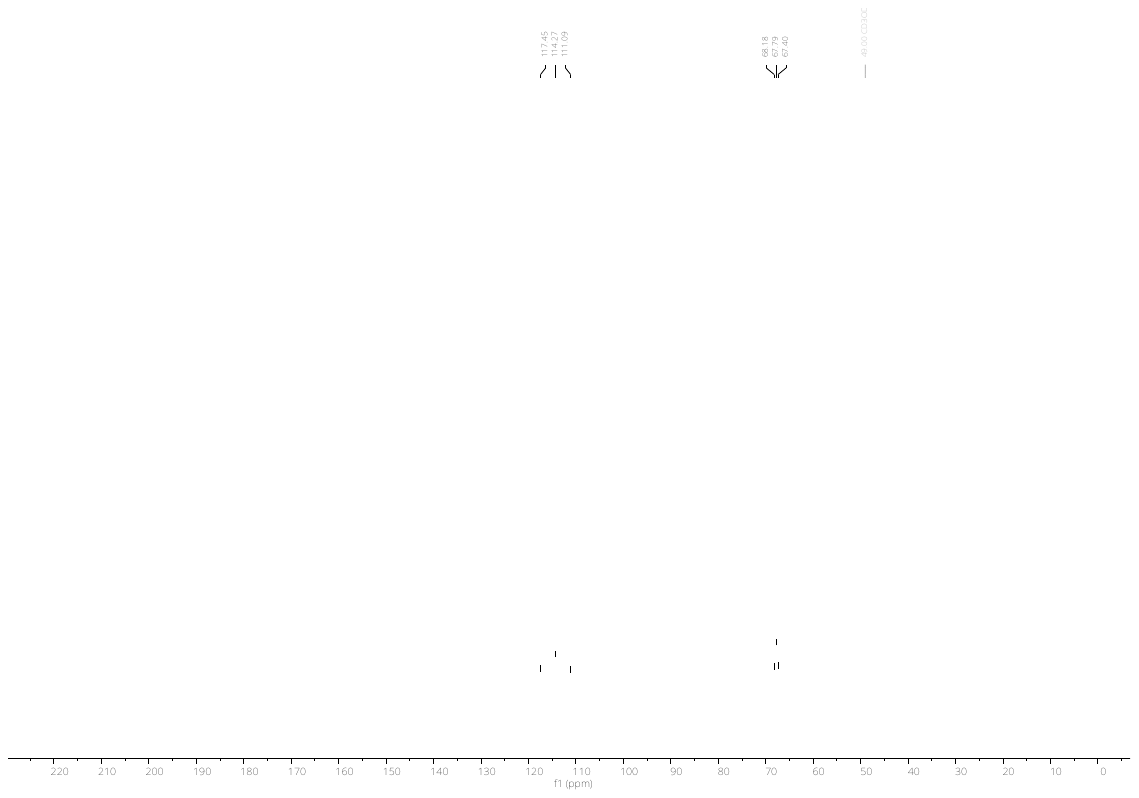

**^13^C NMR**

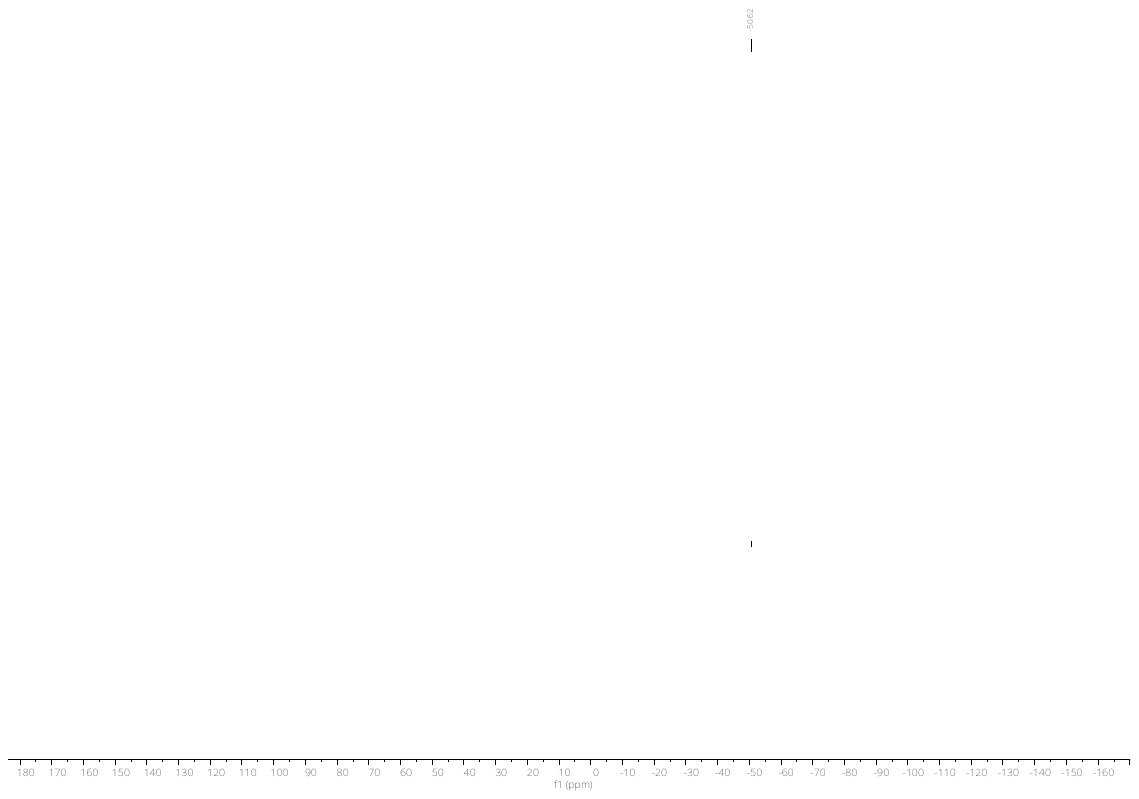

**^19^F NMR**

#### 2,2,3,3,3-pentafluoropropyl sulfamate

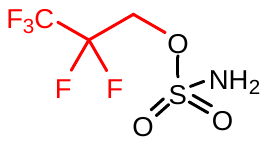

Following the general procedure A with 2,2,3,3,3-pentafluoropropanol yielded the title sulfamate ester (190 mg, 38%). **^1^H NMR** (300 MHz, MeOD) δ 4.70 (tq, *J* = 13.2, 1.1 Hz, 2H). **^13^C NMR** (75 MHz, MeOD) δ 152.05, 61.03, 60.66, 60.30. **^19^F NMR** (282 MHz, MeOD) δ -7.52, -47.28. **HRMS** QTOF-MS: calcd. C_3_H_4_F_5_NO_3_S for [M-H]^-^: cald. 227.9759, found 227.9760

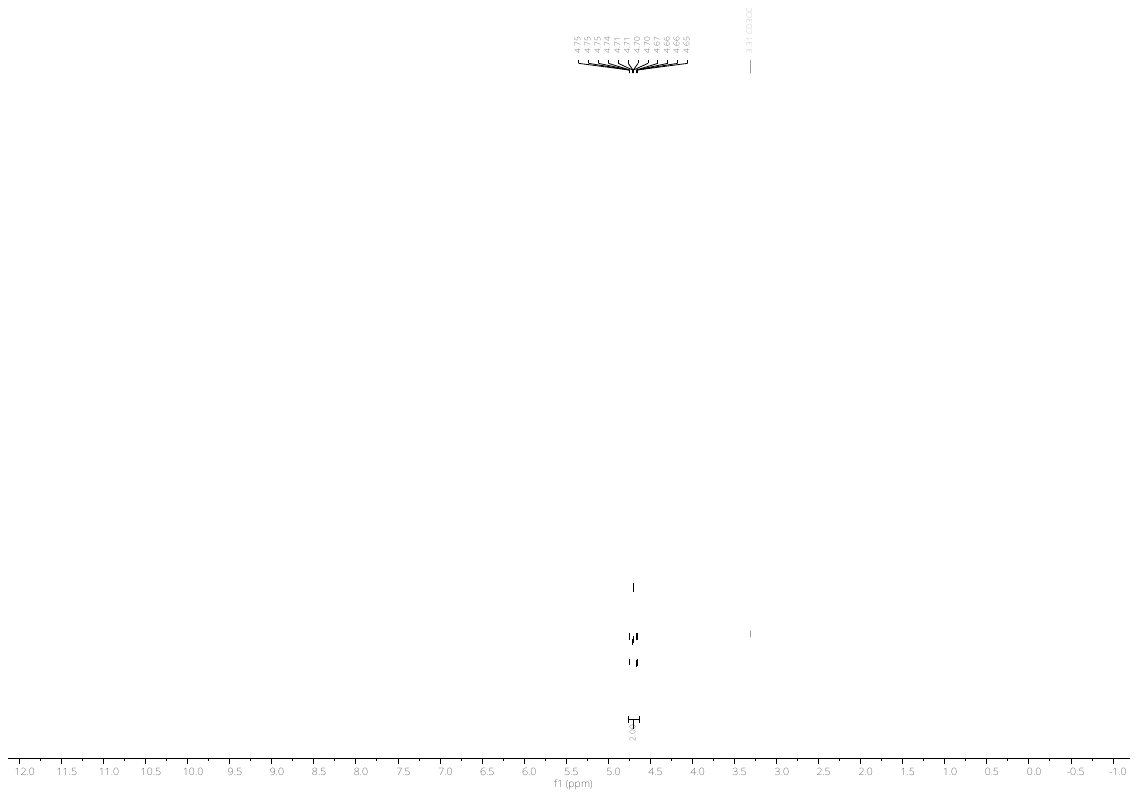

**^1^H NMR**

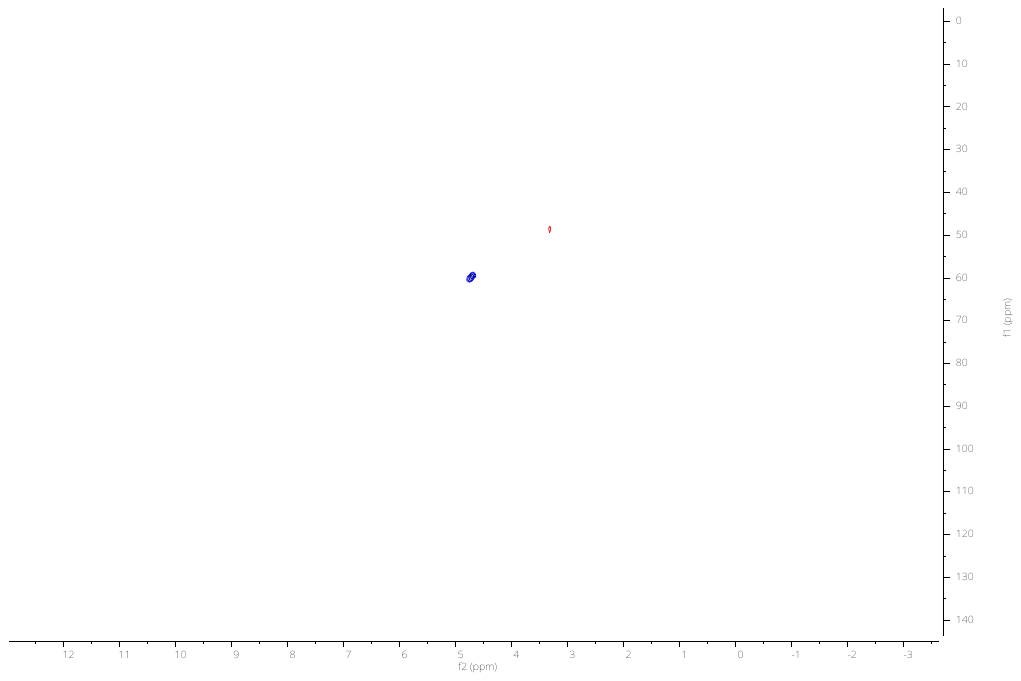

**^1^H-^13^C HSQC NMR**

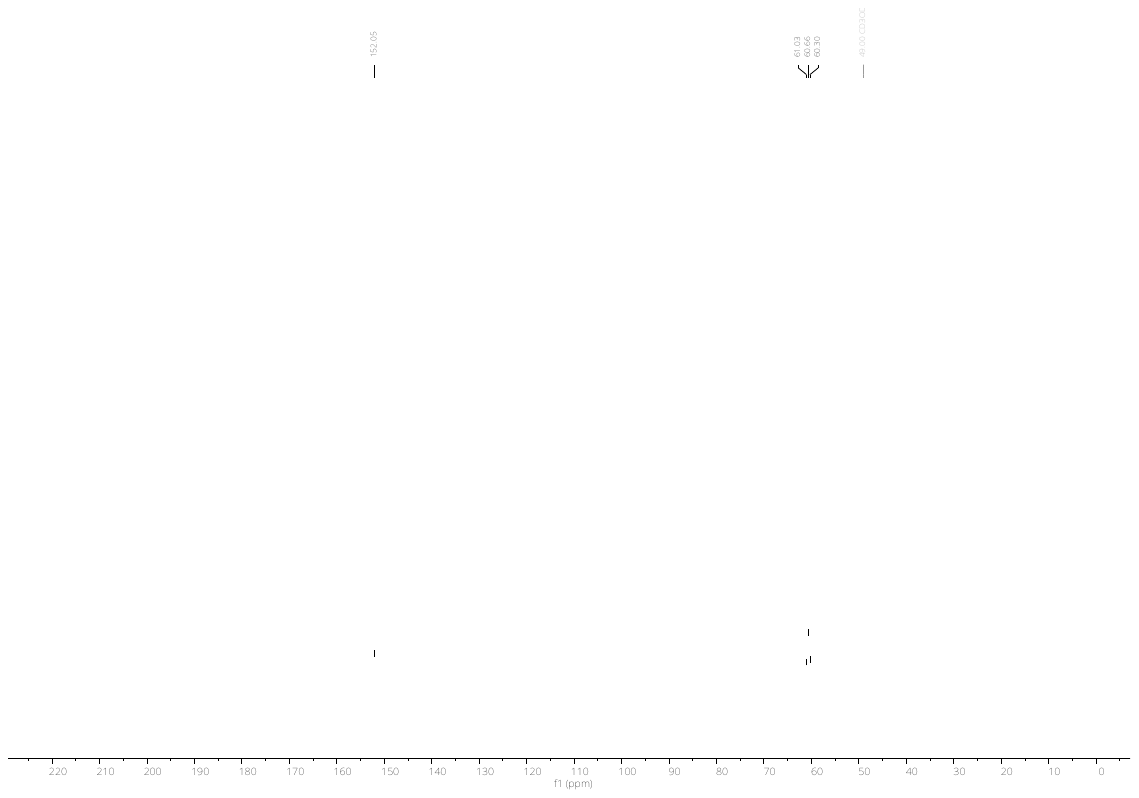

**^13^C NMR**

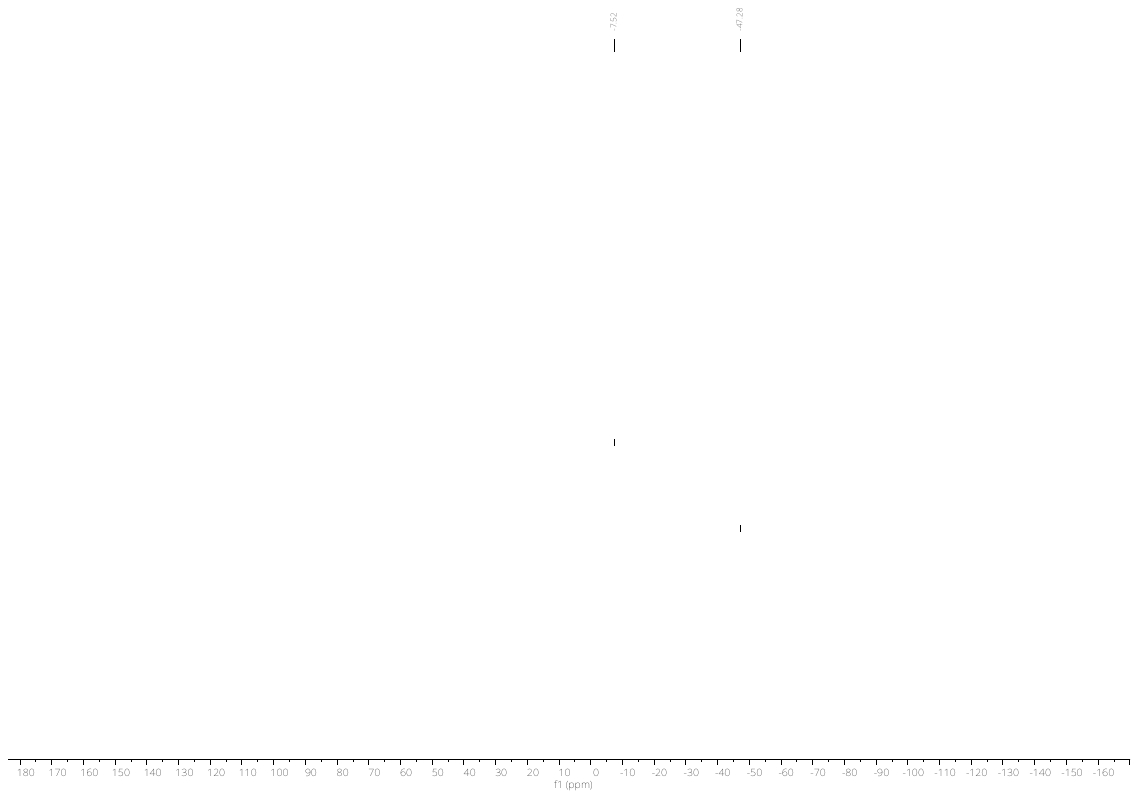

**^19^F NMR**

#### 3-hydroxypropyl sulfamate

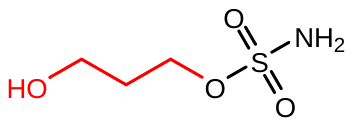

Following the general procedure A with 1,3-propanediol yielded the title sulfamate ester (74 mg, 26%). **^1^H NMR** (300 MHz, MeOD) δ 4.75, 4.29, 4.27, 4.25, 3.70, 3.68, 3.66, 2.20, 2.18, 2.16, 2.16, 2.14, 2.12. **^13^C NMR** (75 MHz, MeOD) δ 67.51, 41.50, 32.83. **HRMS** QTOF-MS: calcd. C_3_H_9_NO_4_S for [M-H]^+^: cald. 154.0180, found 154.0183

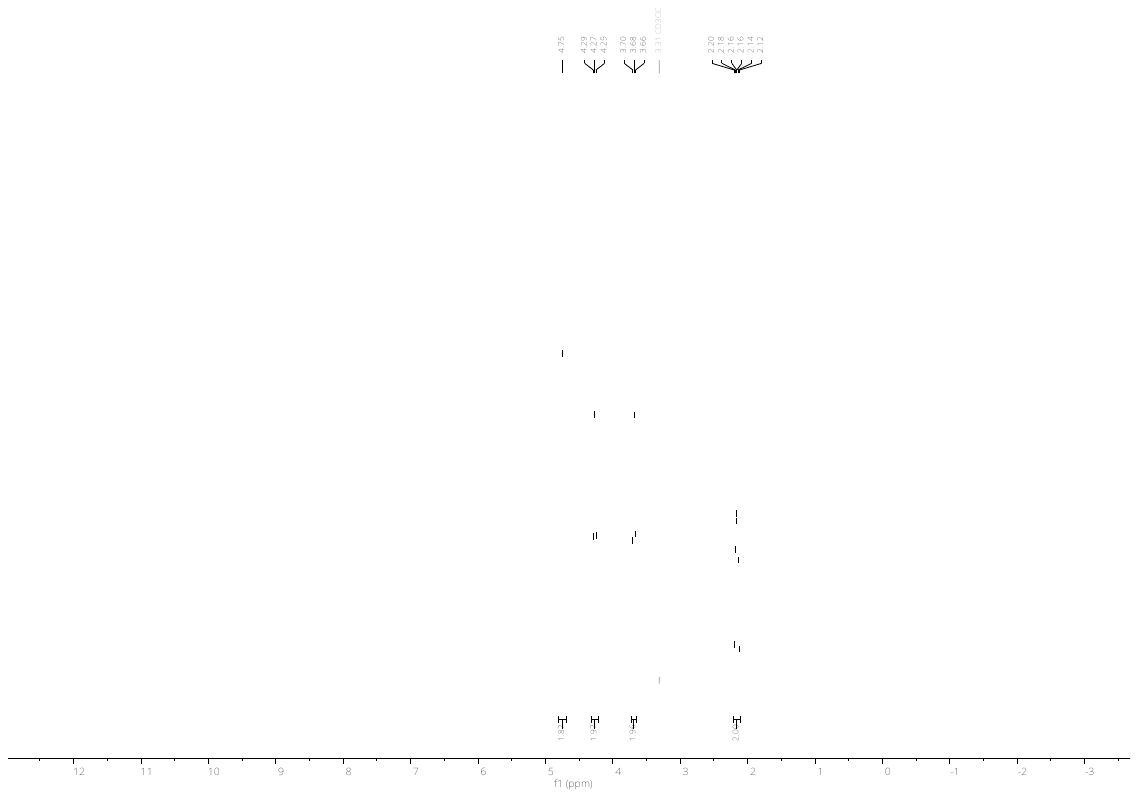

**^1^H NMR**

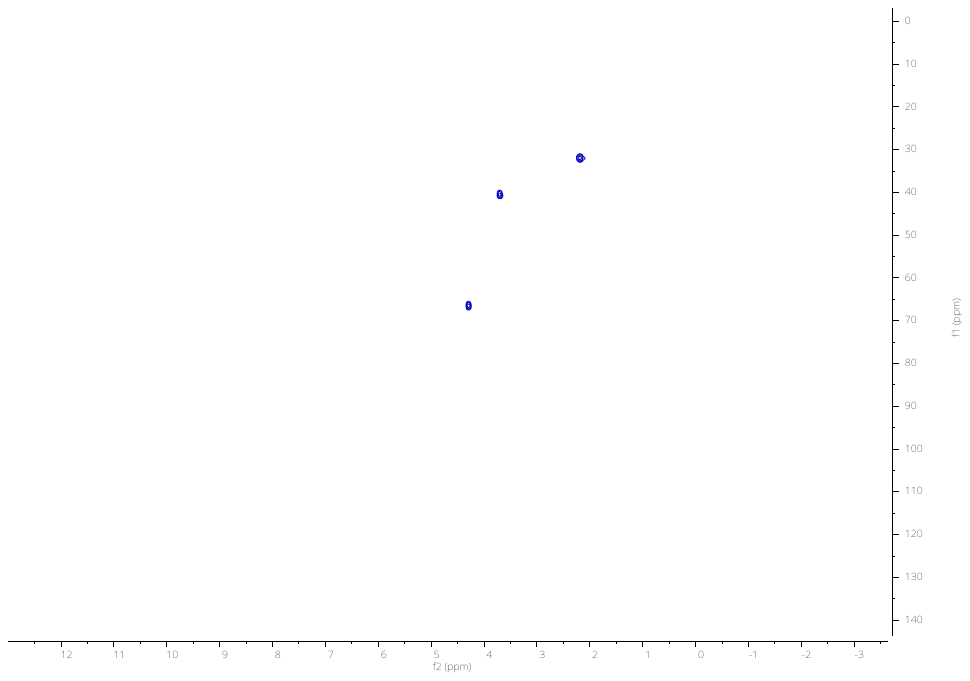

**^1^H-^13^C HSQC NMR**

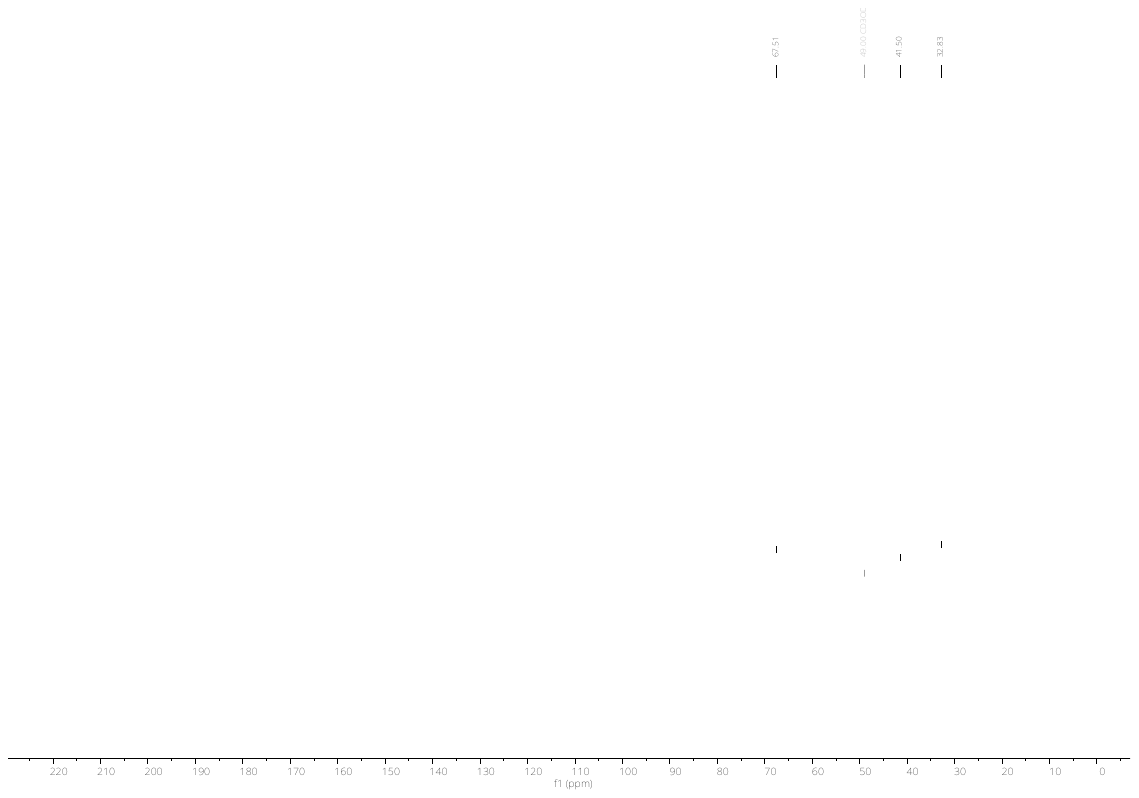

**^13^C NMR**

#### 1,3-dihydroxypropan-2-yl sulfamate

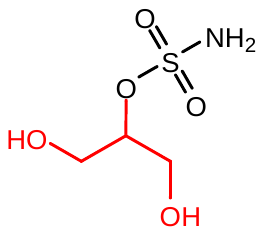

Following the general procedure B with 2,2,3,3,9,9,10,10-octamethyl-4,8-dioxa-3,9-disilaundecan-6-ol yielded the title sulfamate ester (89 mg, 31%). **^1^H NMR** (300 MHz, MeOD) δ 4.67 (qd, *J* = 5.1, 4.3 Hz, 1H), 4.45 – 4.24 (m, 2H), 3.90 – 3.73 (m, 2H).**^13^C NMR** (75 MHz, MeOD) δ 79.67, 68.26, 61.21. **HRMS** QTOF-MS: calcd. C_3_H_9_NO_5_S for [M-H]^-^ : cald. 170.0129, found 170.0131.

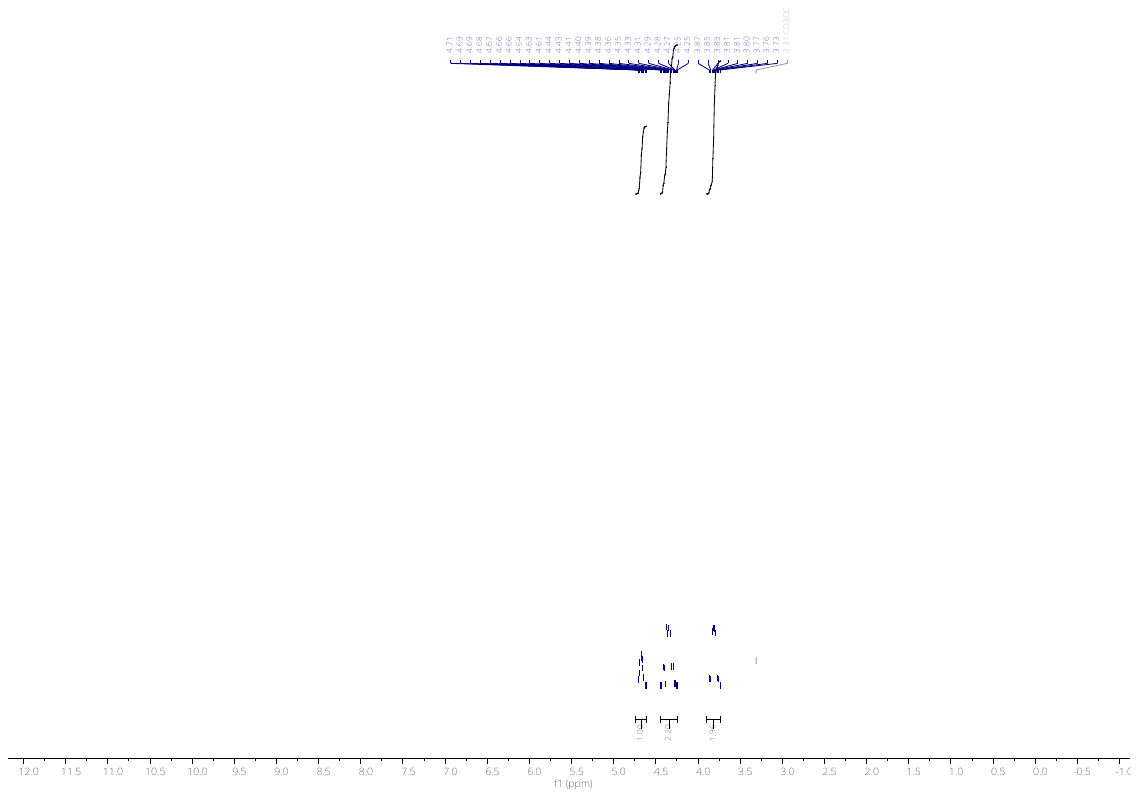

**^1^H NMR**

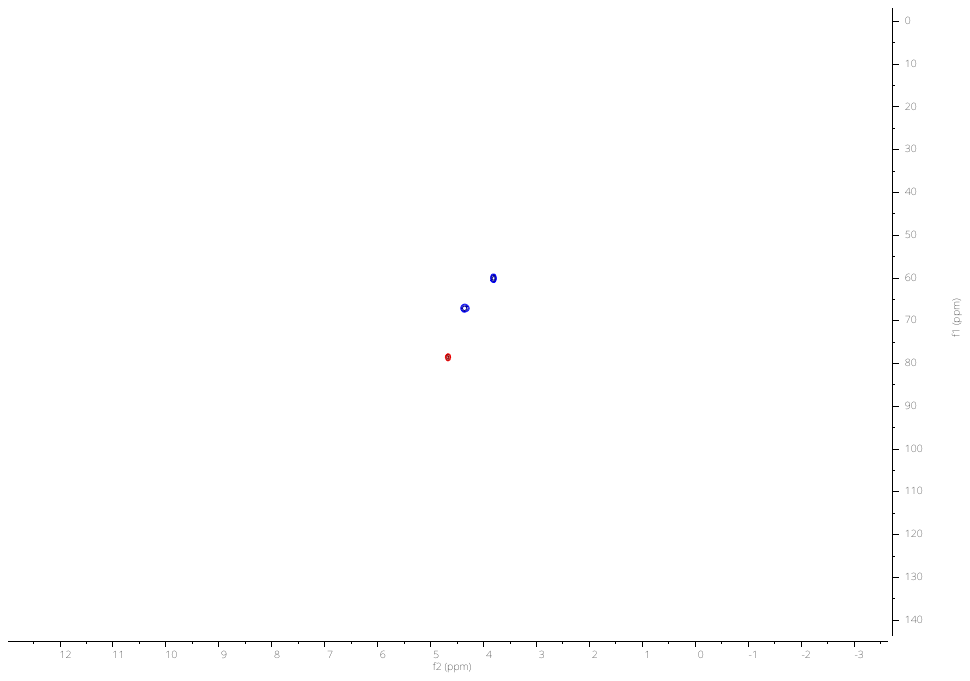

**^1^H-^13^C HSQC NMR**

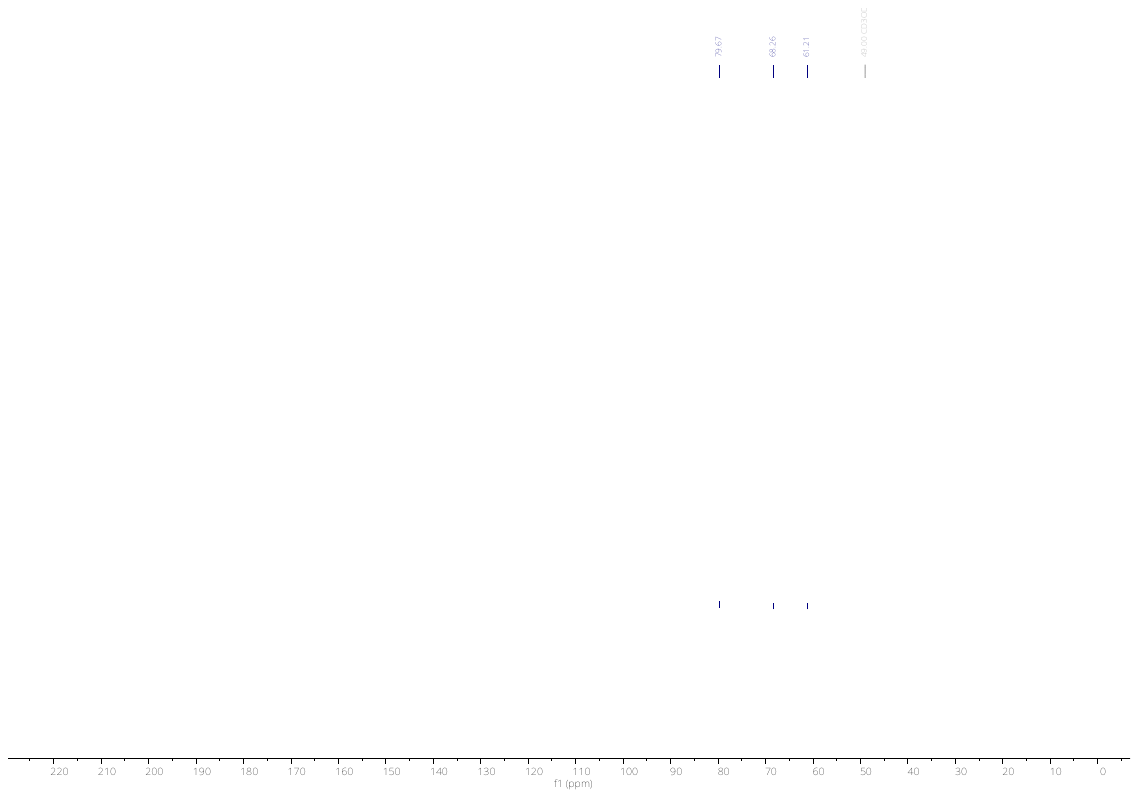

**^13^C NMR**

#### 2,3-dihydroxypropyl sulfamate

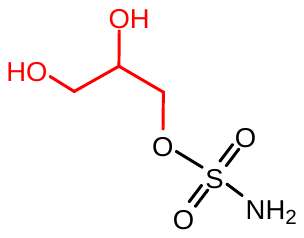

Following the general procedure B with solketal yielded the title sulfamate ester (36 mg, 43%). **^1^H NMR** (300 MHz, MeOD) δ 4.23 – 4.02 (m, 2H), 3.89 (qd, *J* = 5.8, 2.9 Hz, 1H), 3.62 – 3.55 (m, 2H). **^13^C NMR** (75 MHz, MeOD) δ 71.56, 70.95, 63.56. **HRMS** QTOF-MS: calcd. C_3_H_9_NO_3_S for [M+Na]^+^ : cald. 194.0094, found 194.0085.

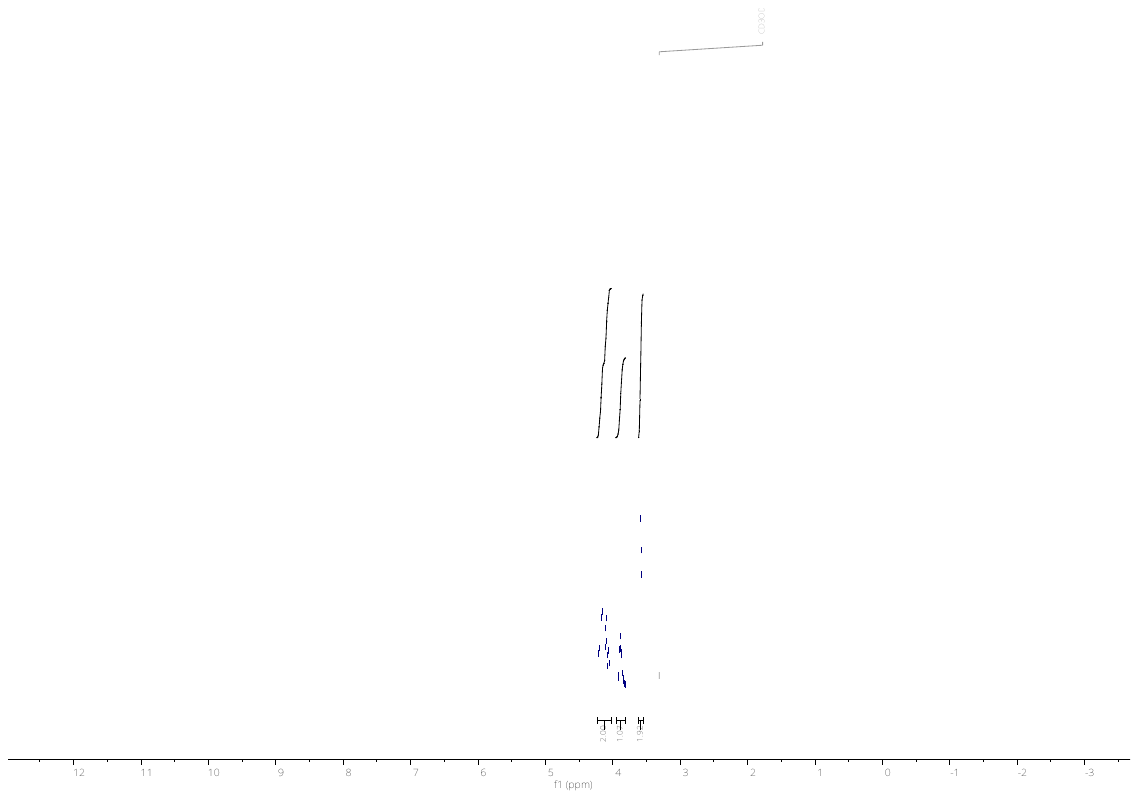

**^1^H NMR**

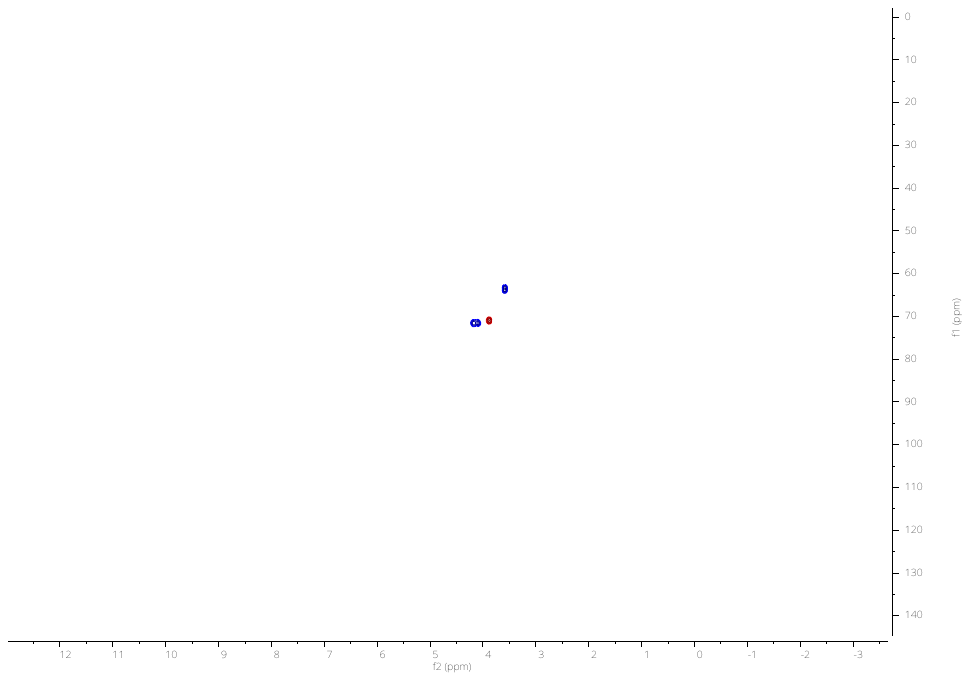

**^1^H-^13^C HSQC NMR**

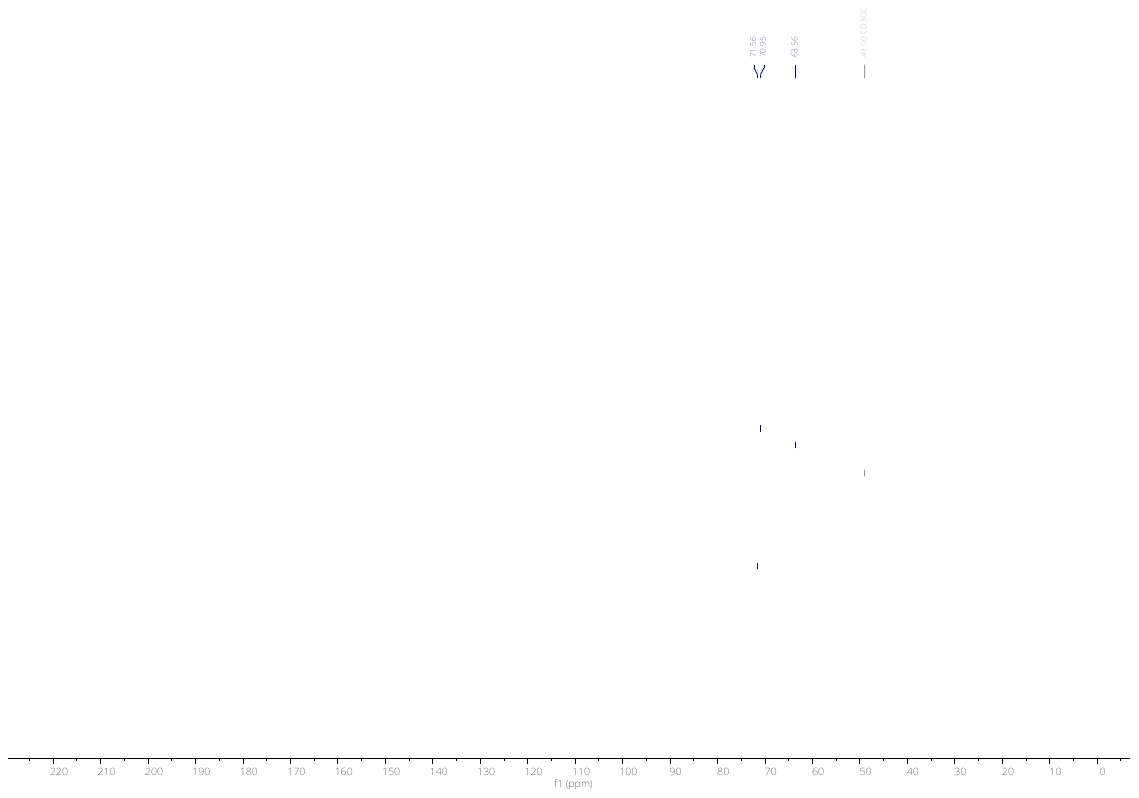

**^13^C NMR**

#### 2-hydroxyethyl sulfamate

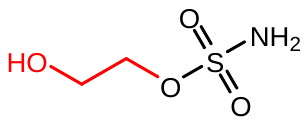

Following the general procedure B with 2-((tert-butyldimethylsilyl)oxy)ethanol yielded the title sulfamate ester (232 mg, 53%). **^1^H NMR** (300 MHz, MeOD) δ 4.22, 4.19, 4.17, 4.16, 4.13, 4.10, 4.09, 3.81, 3.80, 3.79, 3.78. **^13^C NMR** (75 MHz, MeOD) δ 72.23, 60.88. **HRMS** QTOF-MS: calcd. C_2_H_7_NO_4_S for [M+H]^+^ : cald. 142.0169, found 142.0174

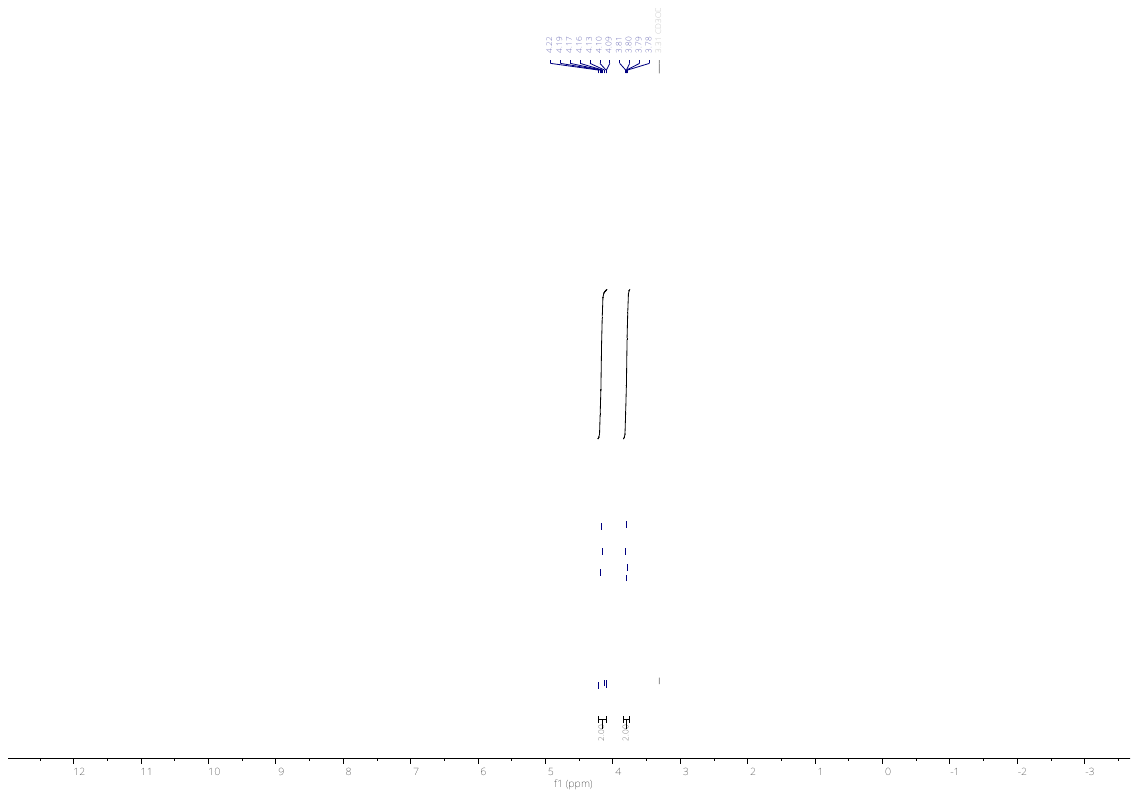

**^1^H NMR**

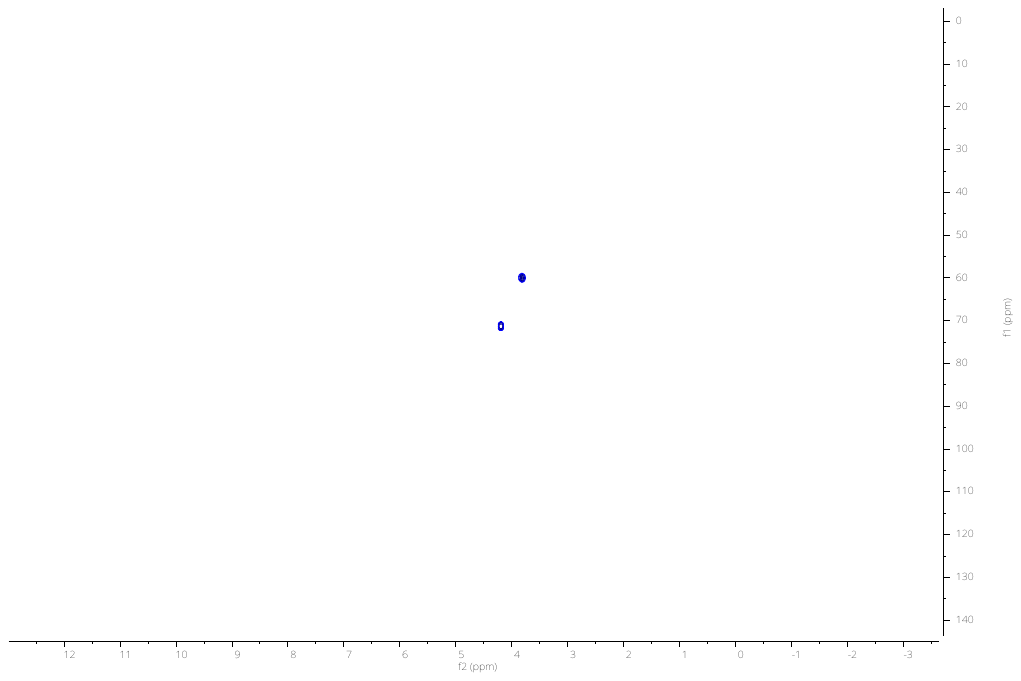

**^1^H-^13^C HSQC NMR**

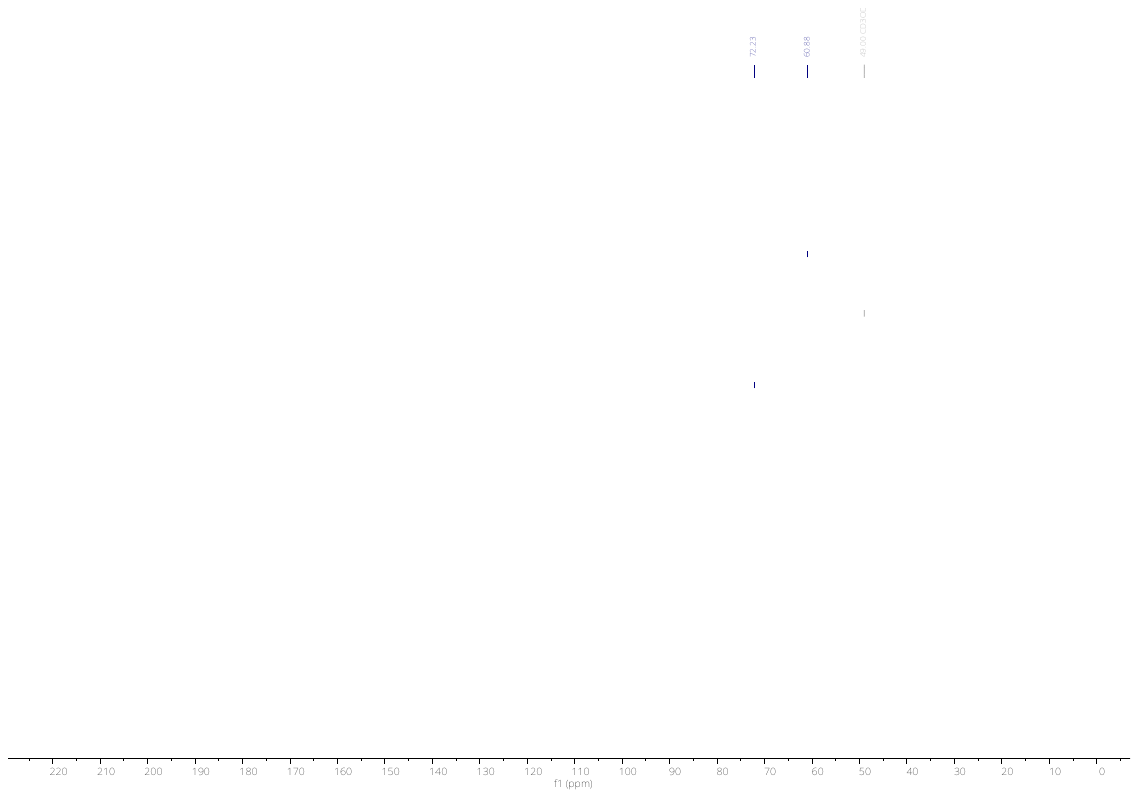

**^13^C NMR**

#### 2-methoxypropyl sulfamate

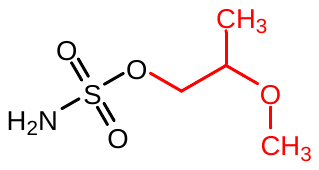

Following the general procedure A with 2-methoxypropan-1-ol yielded the title sulfamate ester (168 mg, 64%). **^1^H NMR** ^1^H NMR (300 MHz, MeOD) δ 4.82 (d, *J* = 4.4 Hz, 2H), 4.15 – 3.96 (m, 2H), 3.73 – 3.57 (m, 1H), 3.39 (s, 3H), 1.19 (d, *J* = 6.4 Hz, 3H). **^13^C NMR** (75 MHz, MeOD) δ 75.91, 72.77, 57.07, 16.35. **HRMS** QTOF-MS: calcd. C_4_H_11_NO_4_S for [M+H]^+^ : cald. 170.0482, found 170.0477.

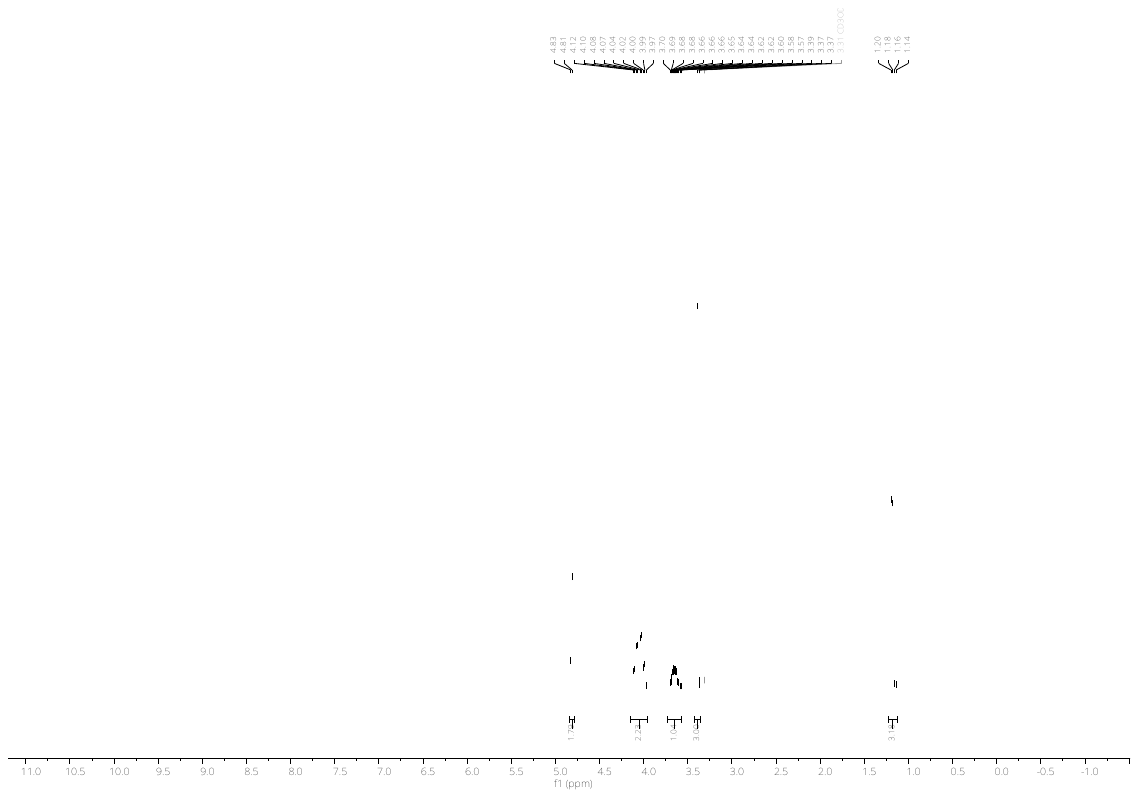

**^1^H NMR**

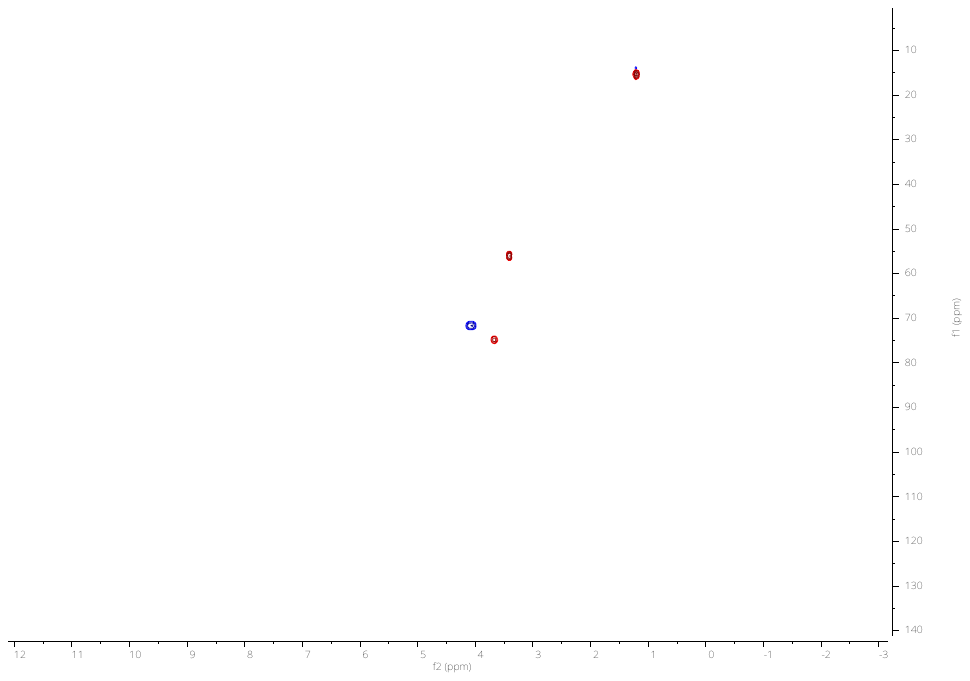

**^1^H-^13^C HSQC NMR**

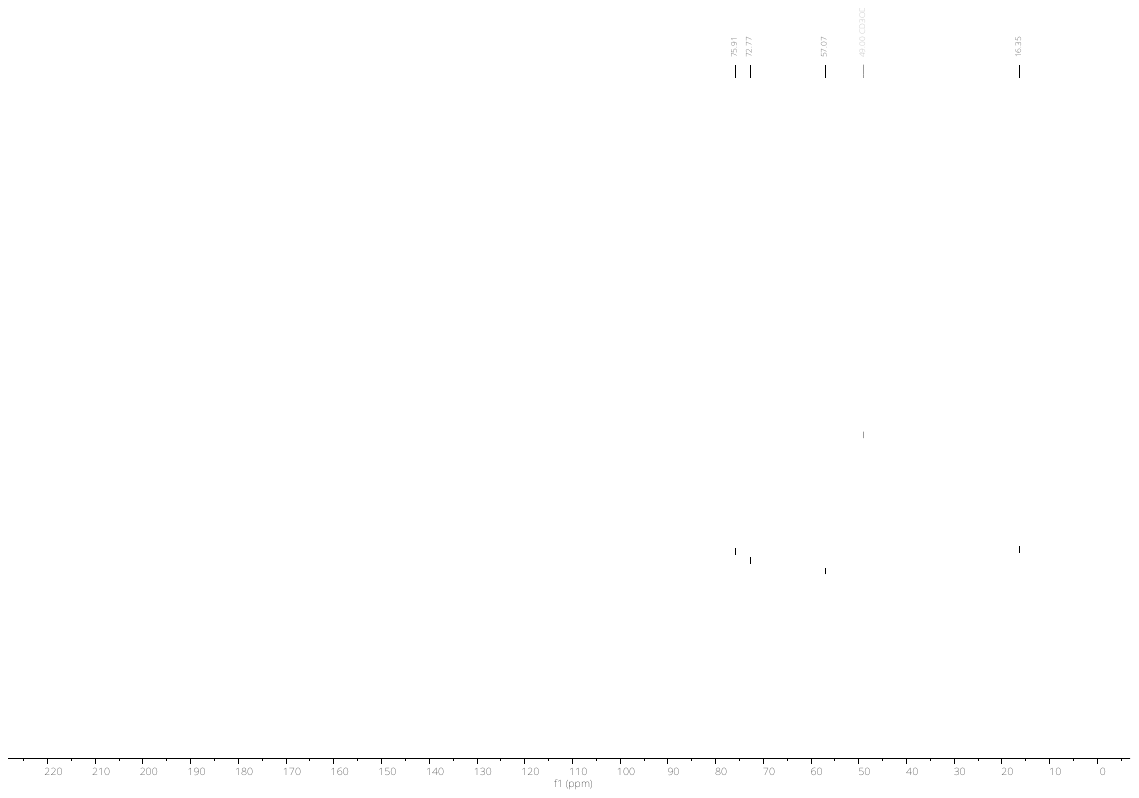

**^13^C NMR**

#### 1-methoxypropan-2-yl sulfamate

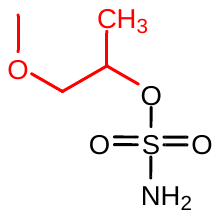

Following the general procedure A with 1-methoxypropan-2-ol yielded the title sulfamate ester (83 mg, 48%). **^1^H NMR** (300 MHz, MeOD) δ 4.74, 4.73, 4.72, 4.72, 4.71, 4.70, 4.70, 4.70, 4.69, 4.68, 4.68, 4.68, 4.66, 4.66, 4.66, 4.64, 3.52, 3.50, 3.50, 3.49, 3.38, 1.36, 1.34.**^13^C NMR** (75 MHz, MeOD) δ 78.02, 76.06, 59.38, 17.60. **HRMS** QTOF-MS: calcd. C_4_H_11_NO_4_S for [M+H]^+^ : cald. 170.0482, found 170.0474

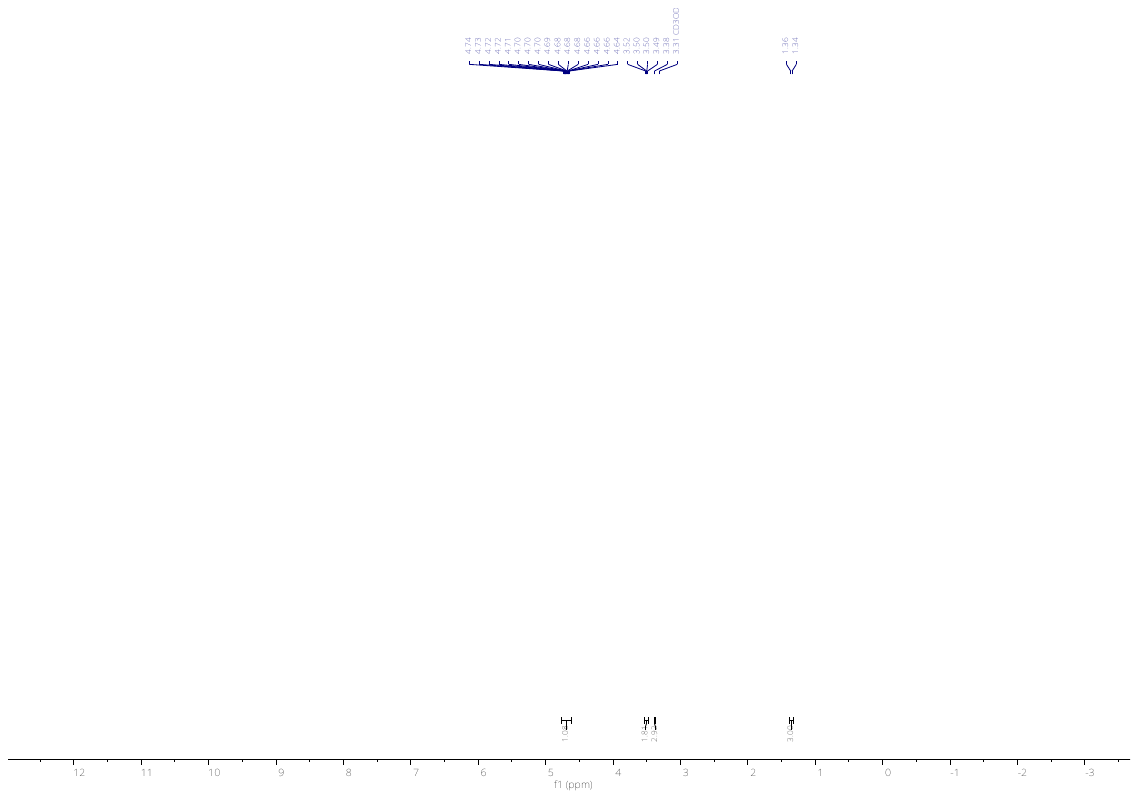

**^1^H NMR**

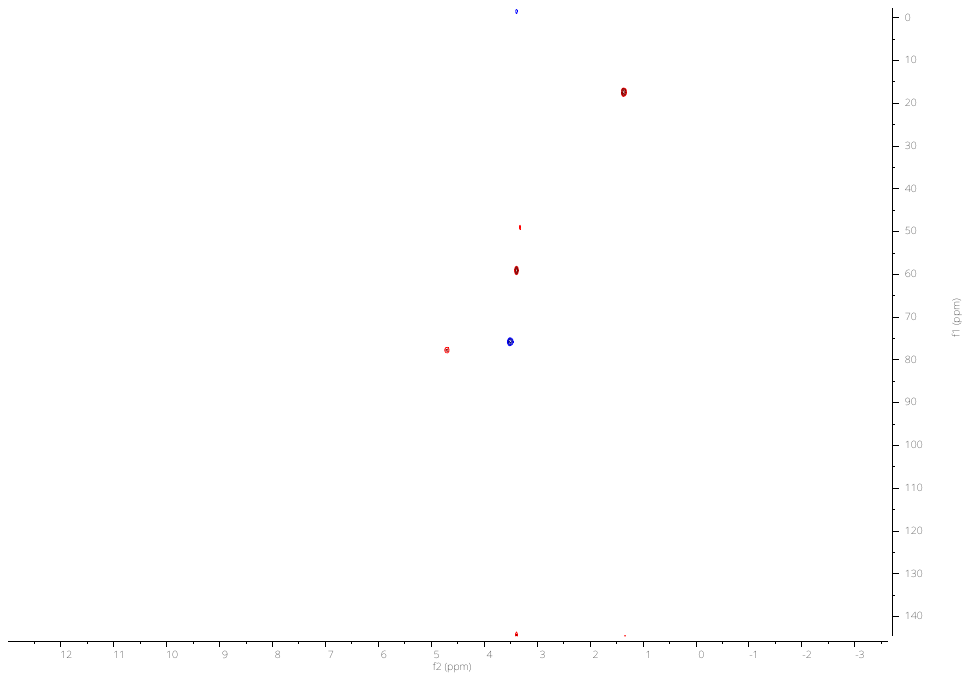

**^1^H-^13^C HSQC NMR**

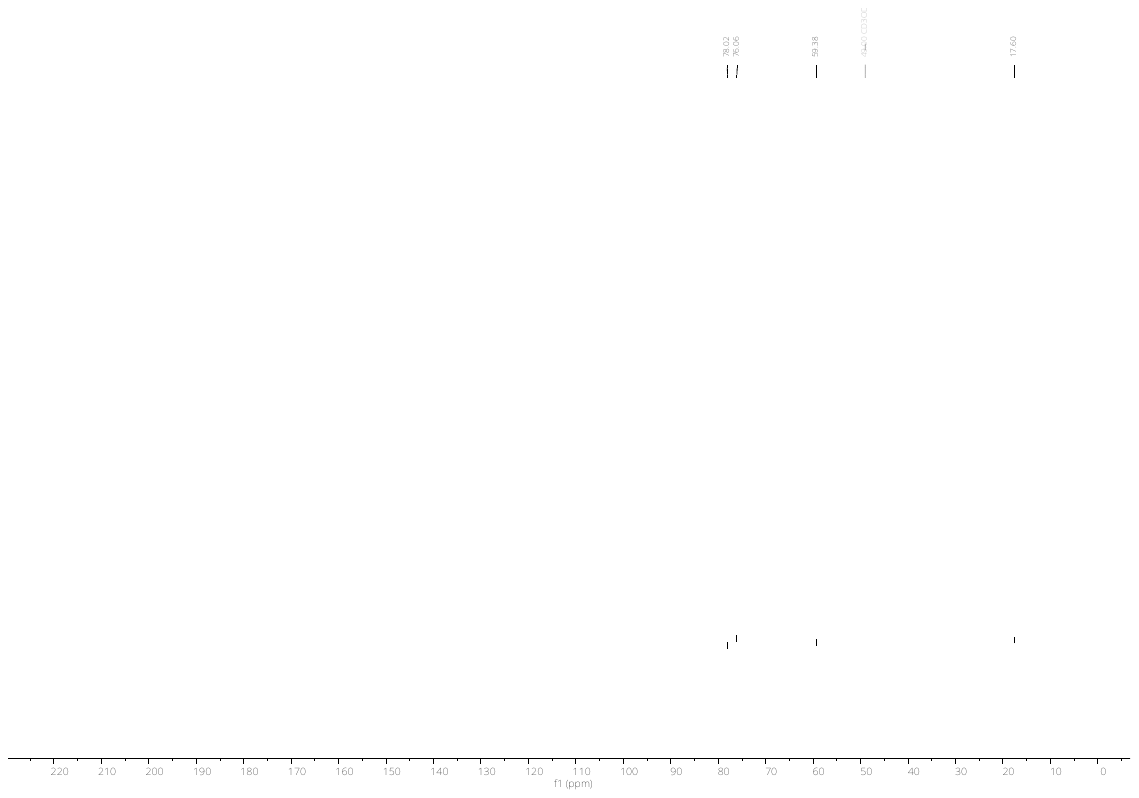

**^13^C NMR**
